# Regularized partial correlation provides reliable functional connectivity estimates while correcting for widespread confounding

**DOI:** 10.1101/2023.09.16.558065

**Authors:** Kirsten L. Peterson, Ruben Sanchez-Romero, Ravi D. Mill, Michael W. Cole

**Affiliations:** Center for Molecular and Behavioral Neuroscience, Rutgers University, Newark, NJ, 07102; Graduate Program in Neuroscience, Rutgers University, Newark, NJ, 07102

**Keywords:** Network neuroscience, fMRI, Regularization, Structural connectivity, Diffusion MRI, Individual differences

## Abstract

Functional connectivity (FC) has been invaluable for understanding the brain’s communication network, with strong potential for enhanced FC approaches to yield additional insights. Unlike with the fMRI field-standard method of pairwise correlation, theory suggests that partial correlation can estimate FC without confounded and indirect connections. However, partial correlation FC can also display low repeat reliability, impairing the accuracy of individual estimates. We hypothesized that reliability would be increased by adding regularization, which can reduce overfitting to noise in regression-based approaches like partial correlation. We therefore tested several regularized alternatives – graphical lasso, graphical ridge, and principal component regression – against unregularized partial and pairwise correlation, applying them to empirical resting-state fMRI and simulated data. As hypothesized, regularization vastly improved reliability, quantified using between-session similarity and intraclass correlation. This enhanced reliability then granted substantially more accurate individual FC estimates when validated against structural connectivity (empirical data) and ground truth networks (simulations). Graphical lasso showed especially high accuracy among regularized approaches, seemingly by maintaining more valid underlying network structures. We additionally found graphical lasso to be robust to noise levels, data quantity, and subject motion – common fMRI error sources. Lastly, we demonstrated that resting-state graphical lasso FC can effectively predict fMRI task activations and individual differences in behavior, further establishing its reliability, external validity, and ability to characterize task-related functionality. We recommend graphical lasso or similar regularized methods for calculating FC, as they can yield more valid estimates of unconfounded connectivity than field-standard pairwise correlation, while overcoming the poor reliability of unregularized partial correlation.

## 1. Introduction

The brain is a complex system, and to fully understand it we must understand how its components interact. Interactions between brain regions or other neural entities (the network “nodes”) are typically investigated using functional/effective connectivity (FC) methods, which quantify network connections (or “edges”) as the statistical relationships between nodes’ neural activities. The most common FC method used with functional magnetic resonance imaging (fMRI) has been, by far, pairwise Pearson correlation (Biswal et al., 1995; Zalesky et al., 2012), with similar pairwise measures being used with other neuroimaging approaches (e.g., coherence with electroencephalography [EEG] or magnetoencephalography [MEG]; Srinivasan et al., 2007). Pairwise correlation FC is easy to interpret and compute, and it can indicate the overall amount of activity that is common between nodes. However, it is flawed as a measure of direct interactions, which is desirable when modeling network topology, causal relationships, and the specific pathways over which information is spread. If FC is intended to represent direct functional interactions, as we define it here, then pairwise correlation severely overestimates FC, reporting not only direct connections but numerous confounded and indirect connections as well (for review see Friston, 2011; Reid et al., 2019). These false positives can be reduced by instead using multivariate FC methods such as partial correlation and multiple regression (Cole et al., 2016; Marrelec et al., 2006; Reid et al., 2019; Smith et al., 2011). Such methods improve upon pairwise correlation by conditioning on the time series of all other measured nodes (Figure 1A), allowing them to resolve confounding and indirect influences to more validly estimate direct connectivity. In other words, pairwise correlation measures the total activity shared between two regions, which may be fully or partly mediated by other regions, while partial correlation measures the activity shared by only the two regions of interest that are not accounted for by other regions. Note that there may be cases in which the total activity shared between two neural populations is desired, in which case pairwise correlation should be preferred. However, there are a wide variety of circumstances (e.g., modeling network topology, causal relationships, and the specific pathways over which information is spread) in which estimating direct FC may be preferred.

**Figure 1.**
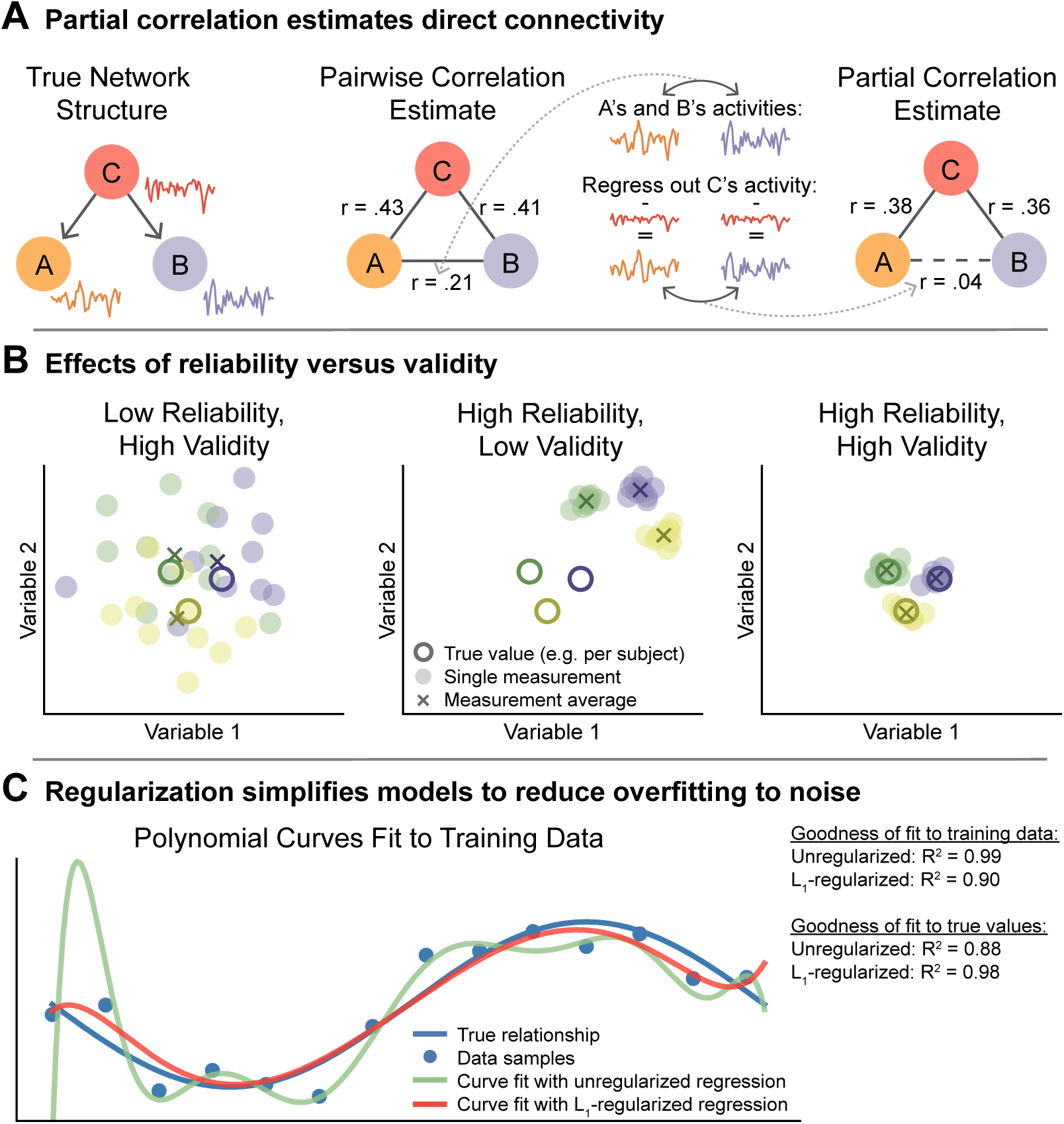
– Partial correlation corrects for confounds but suffers from low reliability, likely due to overfitting, which can be reduced with regularization. **A)** Partial correlation can estimate true direct connectivity more validly than pairwise correlation (the field standard for fMRI) in the presence of confounders (node *C*) (Reid et al., 2019). Here, regions *A* and *B* show a strong pairwise correlation despite not interacting, due to the third region *C* influencing (i.e. confounding) both *A* and *B*. Partial correlation takes region *C* into account to properly estimate the relationship between *A* and *B*. Empirical tests suggest confounder connectivity patterns are widespread throughout the brain, leading to thousands of false connections in many fMRI FC studies (Sanchez-Romero and Cole, 2021). Note that, while the example true network contains asymmetric edges that act in a single direction, all FC methods tested here estimate symmetric, undirected edges. **B)** An FC method must be both reliable and valid for its individual measurements to be consistently accurate (right panel). Recent studies have suggested that pairwise correlation is reliable yet invalid as a measure of direct FC (center) while partial correlation is unreliable but largely valid (left; Mahadevan et al. 2021; Fiecas et al. 2013). Note that colors represent distinct entities such as subjects or task states, measured across multiple sessions. **C)** We hypothesized that partial correlation and related multivariate FC methods may show low reliability due to overfitting to noise, since estimated model coefficients that are more readily influenced by noise will be less stable with repeated sampling. We illustrate the issue of overfitting by fitting polynomial curves to simulated data (blue dots; true curve plus noise), where differing regression models were used to estimate the coefficients of polynomial terms. Note that multiple regression is the statistical basis for computing partial correlation. The curve produced by unregularized multiple regression (green line) is overly complex and specific to the noise in the training data. It differs substantially from the true underlying relationship (blue line) and would also differ from new curves fit to new data. L1-regularization constrains the flexibility of the model and results in a simpler but more accurate curve (red line).

While partial correlation and related FC methods have a clear theoretical advantage over pairwise correlation in reporting direct functional connections, they have also been shown to have worse repeat reliability (Fiecas et al., 2013; Mahadevan et al., 2021). Reliability gauges the stability of measurements and is essential to the accuracy of individual estimates. However, validity is an even more important criterion, assessing how well a method measures what it intends to measure. In this study, we define validity as being independent of reliability, evaluating the systematic correctness of a method, or how close the average of many repeated estimates is to the true value. Validity and reliability as used here can therefore be considered the analogues of bias and variance, concepts in statistics and machine learning. We then define individual measurement accuracy (or just “accuracy”) as reflecting the closeness of individual estimates to the true value, a function of both validity and reliability that measures overall correctness. If a method is valid but unreliable (Figure 1B, left panel), as has been shown for partial correlation FC (Fiecas et al., 2013; Mahadevan et al., 2021), then its individual measurements would be frequently dissimilar from the true values due to their high variability, although they may approximate the truth in aggregate (e.g., after averaging over many sessions from a single subject, or many subjects in a population). Alternatively, a method can be reliable but not valid (Figure 1B, center), which more resembles pairwise correlation FC when aiming to measure direct connections (Fiecas et al., 2013; Mahadevan et al., 2021; Sanchez-Romero & Cole, 2021). Such resulting measurements would be close to each other but far from the truth, meaning that none of the individual measurements were accurate representations. An ideal method would be reliable and valid (Figure 1B, right), producing stable measurements that are each close to the truth.

Given its otherwise high validity, partial correlation has the potential to be an extremely useful FC method if its low reliability were overcome. We hypothesized that the reported instability of partial correlation and related multiple regression-based FC methods occurs from overfitting to noise and could be ameliorated by regularization techniques. Overfitting to noise often results from a model’s excessive complexity, and it can be exacerbated by factors such as low quantity and poor quality of the data being fit (Blum et al., 2020; Hastie et al., 2009; Ying, 2019). Such complexity arises in partial correlation and multiple regression FC with fMRI as the models fit more variables to include all measured nodes (brain regions or voxels). Increasing the complexity of a model allows it to better fit the specific training data by accounting for more variance, including noise. By capturing arbitrary patterns in the training data, such an overfit model will not generalize to independent data or accurately estimate coefficients (Blum et al., 2020; Hastie et al., 2009; Lever et al., 2016; Ying, 2019). We illustrate this problem in Figure 1C by fitting a polynomial regression model to noisy data, demonstrating that a more complex model (i.e., a model with more variables) overfits to the noise, reducing model reliability as it will fit to unique noise with every new data sample. In the case of FC estimation, overfit regression models will adjust connectivity coefficients to incorporate chance similarities in regions’ activities, resulting in unstable coefficient weights that seldom reflect true connectivity values.

Regularization is a common strategy for reducing model complexity, and therefore reducing overfitting to noise, in both statistics and machine learning (Blum et al., 2020; Hastie et al., 2009; Ying, 2019). A variety of regularization techniques have been developed, such as explicitly penalizing complexity during model fitting. Two common methods are L_1_ (lasso) and L_2_ (ridge) regularization, which simplify models by penalizing them by the summed absolute values and squares of their coefficients, respectively (Hoerl & Kennard, 1970; Tibshirani, 1996). Another regularization method – principle component (PC) regression – works by fitting the model to a subset of PCs, reducing the number of variables (and hence model complexity) but keeping much of the presumed signal (Jolliffe, 1982).

While applying regularized multivariate methods to FC estimation is not entirely novel, the practice remains substantially under-used relative to pairwise correlation. Indeed, several studies have tested and recommended regularized multivariate FC (Brier et al., 2015; Duff et al., 2013; Pervaiz et al., 2020; Smith et al., 2013), with some even introducing new implementations (Mejia et al., 2018; Nie et al., 2017; Ryali et al., 2012; Varoquaux et al., 2010). Still, many other studies that utilized regularization did not clearly recommend the extra step (Fiecas et al., 2013; Mahadevan et al., 2021; Smith et al., 2011), and some analyses that used FC for prediction did not demonstrate an unequivocal benefit of regularized multivariate methods over pairwise correlation (Duff et al., 2013; Sala-Llonch et al., 2019). In addition, while there are many regularized methods available, few studies have tested the differences between them and even then typically to a limited extent (Brier et al., 2015; Mejia et al., 2018; Nie et al., 2017; Pervaiz et al., 2020; Ryali et al., 2012; Varoquaux et al., 2010). This lack of comparison between methods can leave researchers uncertain of which regularization approach to implement. Such ambiguities, as well as a lack of understanding of these regularized multivariate methods, may prevent researchers from adopting methods that would otherwise benefit their analyses.

The purpose of this study is to test the suitability of regularized partial correlation and similar multivariate methods for estimating direct, more causally and mechanistically valid FC. To do this, we compared the performances of three established regularized methods – graphical lasso, graphical ridge, and PC regression – with unregularized partial correlation and the field-standard pairwise correlation. The use of three regularized methods allowed us to generalize utility of the fundamental concept of regularization across techniques, as well as test for differences between them.

We began by separately testing the reliability, validity, and individual measurement accuracy of the FC methods, where we define validity as being independent of reliability while accuracy is not (see above). Reliability was quantified using between-session similarity and intraclass correlation, with repeated measures being compared to each other. Since we cannot access the ground truth of empirical FC, validity and accuracy were quantified by comparing FC estimates against two different informative but imperfect targets. First, we compared empirical FC estimates with the corresponding subjects’ structural connectivity (SC), computed from diffusion MRI data. Although FC and SC reflect different phenomena, direct functional connections should require a white matter link, and the measures should therefore show a degree of convergence. As a second strategy, we compared FC estimated from simulated data to the ground truth simulation networks. In both cases, individual measurement accuracy was quantified as the closeness of individual FC estimates to the target, while validity was the closeness of the collective, group-averaged FC estimates. We then extended our assessments of the FC methods to their resilience against fMRI practical pitfalls: short scan lengths, scanner noise (Blum et al., 2020; Hastie et al., 2009; Ying, 2019), and subject head movement artifacts (Power et al., 2015). Lastly, we further broadened our assessment of FC accuracy to task-related functionality, using FC estimates to generate held-out task activations with activity flow modeling (Cole et al., 2016) as well as to predict individual differences in subject age and intelligence. These comprehensive tests show the effectiveness of regularized partial correlation in estimating direct FC and its advantages over the field-standard pairwise correlation FC approach.

## 2. Methods

### 2.1. Functional connectivity estimation methods

This study compared the performance of five FC methods: pairwise Pearson correlation, partial correlation, graphical lasso, graphical ridge, and PC regression. All methods estimated FC from the same empirical resting state or simulated timeseries, which were always z-scored (Hastie et al., 2009). Each FC matrix was calculated independently from a single session of data, coming from a single subject or simulated network.

Note that we have released code to implement graphical lasso FC – including hyperparameter selection and FC estimation – as part of the Brain Activity Flow Toolbox (https://colelab.github.io/ActflowToolbox/) (Cocuzza et al., 2022).

#### 2.1.1. Pairwise correlation

Pairwise correlation FC was computed as the Pearson correlation between each pair of nodes’ timeseries.

#### 2.1.2. Partial correlation

Partial correlation can be calculated in two ways with near identical results. The more intuitive approach involves regressing the timeseries of all other nodes (the conditioning variables) from the two target nodes’ timeseries, and then computing the Pearson correlation between the two targets’ residuals (Figure 1A). This study instead used the inverse covariance approach, which is advantageous because it is less computationally expensive. First the covariance matrix of all nodes is inverted, giving the precision matrix *P*. Then the partial correlation coefficients are calculated as:

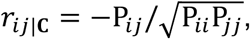

for nodes *i* and *j* conditioned on set **C**.

#### 2.1.3. Graphical lasso

The first of our regularized methods, graphical lasso (”glasso”) implements partial correlation with an L_1_ penalty to limit model complexity (Friedman et al., 2008; Tibshirani, 1996). The penalty is applied when computing the precision matrix, and this regularized precision matrix is then transformed into the partial correlation matrix. L_1_ regularization works by adding to the model cost function the term:

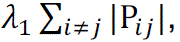

which is proportional to the summed absolute values of entries in the estimated precision matrix (*P*), not including the diagonal. The amount of regularization is scaled by the hyperparameter λ_1_. L_1_ regularization tends to drive less informative coefficients to exactly zero, producing a sparse result. In this way, L_1_ regularization can also perform feature selection. We implemented graphical lasso using the Python package GGLasso (Schaipp et al., 2021).

#### 2.1.4. Graphical ridge

Graphical ridge applies L_2_ regularization (also called Tikhonov regularization) to partial correlation (Hoerl & Kennard, 1970). It applies the L_2_ penalty term:

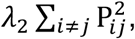

which is the summed square values of entries in the estimated precision matrix (*P*) multiplied by the hyperparameter λ_2_. While L_2_ regularization also encourages model simplification (i.e., by bringing coefficients closer to zero), it does not cause sparsity to the extent that L_1_ regularization does (shrinking less meaningful coefficients to exactly zero). Because of the square term, L_2_ regularization exerts uneven pressure on edges based on their coefficient weights, with low coefficients contributing disproportionately small amounts to the penalty term. As coefficients are shrunk, their penalties become increasingly negligible so that they are seldom brought to exactly zero as occurs with L_1_ regularization. Meanwhile, high coefficients produce comparatively much larger penalties, causing the model to be biased against higher weights. In this way, L_2_ regularization encourages a narrower range of weights that are shared more evenly across coefficients (see Figure 2). We implemented graphical ridge using the R package rags2ridges (Peeters et al., 2022).

**Figure 2.**
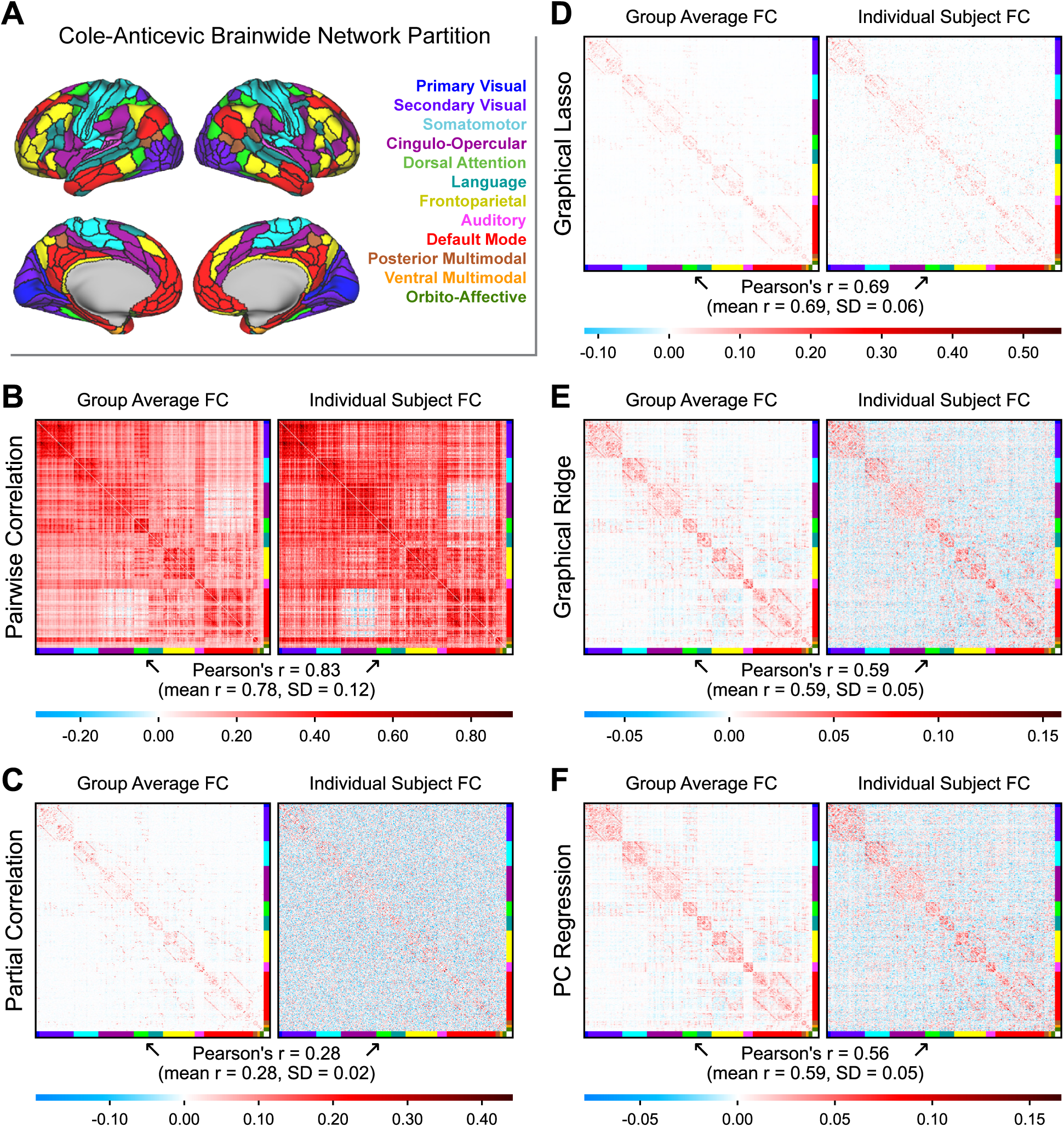
– Empirical FC matrices produced by different methods. The nodes were ordered according to the Cole-Anticevic Brain-wide Network Partition **(A)** to visualize the network structure (Ji et al., 2019). Group-averaged connectivity matrices (N=236; left) are shown beside a random subject’s individual matrices (right). The Pearson correlation between the individual FC matrix and the group average is printed below each set of matrices. **B)** Pairwise correlation (the field standard), **C)** partial correlation, **D)** graphical lasso, **E)** graphical ridge, and **F)** PC regression.

#### 2.1.5. Principal components regression

PC regression combines multiple regression with principal component analysis (PCA) to induce regularization (Hastie et al., 2009; Jolliffe, 1982). To construct an FC matrix using any multiple regression method, a regression model is fit once for each node, with that target node’s timeseries being predicted by the timeseries of all other nodes. The row of the FC matrix that corresponds with the target node is filled in with the beta coefficients from all predictor nodes, in the corresponding columns. The FC matrix is filled row by row in this way. To perform PC regression for a target node (single row of matrix), PCA is applied to the predictor nodes’ timeseries and only a subset of the PCs are used as predictor variables in the regression model, usually selected as the *n* PCs accounting for the highest variance. The number of PCs to include is a hyperparameter which determines the amount of regularization, where using fewer PCs leads to more regularization. This reduces overfitting to noise by reducing the number of variables in the regression model and presumably discarding noisier or less relevant dimensions of data. After fitting the PCs to the original to-be-predicted target timeseries, the PC coefficients are transformed to reflect the predictor nodes’ contributions, which become entries in the FC matrix. We implemented this method using the LinearRegression and PCA functions from Python package Scikit-learn. Using multiple regression to calculate FC independently for each target node creates an asymmetrical matrix, as the contribution of a node *X* to predicting target *Y* is unlikely to be identical to the contribution of *Y* when *X* is the target. This does not necessarily represent directionality of connections as FC asymmetry usually implies, since additional noise in predictor node *X*’s time series would reduce its FC value without there being a change in causal influence. Therefore, we manually symmetrized each PC regression FC matrix by averaging the original with its transpose.

#### 2.1.6. Hyperparameter selection

Regularization often requires the selection of hyperparameters, the choice of which can greatly impact the resulting FC estimates. We therefore had to optimize the hyperparameters within each regularized method before making comparisons between methods. In the methods we tested, the hyperparameters are λ_1_ for graphical lasso, λ_2_ for graphical ridge, and number of PCs for PC regression. For graphical lasso and graphical ridge, lambda values of zero would coincide with unregularized partial correlation and higher values would lead to a greater degree of regularization. For PC regression, using all components would yield unregularized multiple regression while using fewer components would produce more regularization. If the hyperparameter of any method does not induce enough regularization, then (according to our hypothesis) the model will remain overfit, making the FC estimates unreliable (i.e., having high variance). If the hyperparameter induces too much regularization, however, the model will be underfit, discarding relevant information such that the FC estimates are less valid (i.e., having high bias; Hastie et al., 2009; Lever et al., 2016). The optimal value would balance these two tendencies.

We sought a measure of model fit that could be applied in the same way for all FC methods that we tested, to limit the possibility of model fit metrics biasing FC method performance. Our solution was to determine optimal hyperparameter values as those whose models can best predict held-out timeseries data. This form of cross-validation is a standard machine learning approach for testing a regression model’s accuracy in the context of fitting time series (Pardoe, 2020). Resting-state fMRI time series were used for both model fitting and testing with held-out data. For each row of an FC matrix (representing all connections to a single node), the connectivity weights were treated as beta coefficients in a regression model to predict that single node’s held-out activity from the concurrent activities of all other nodes. Prediction accuracy was then calculated as the coefficient of determination (R^2^) between predicted and actual activities. This was applied with 10-fold cross-validation within each individual session timeseries, where each fold served as held-out, to-be-predicted data for one iteration while all other folds were used to compute the FC matrix. We made sure to use a similar number of timepoints to produce these cross-validation matrices as we would for the final matrices (9 of 10 folds means 90% of timepoints), as we found that models fit with less data tend to prefer a greater level of regularization. This was repeated for all tested hyperparameter values, and the optimal hyperparameter for each session was that which produced the highest R^2^ value averaged over all folds. If the parameter value selected for a given session was the smallest or largest in the range being tested, then we expanded the parameter range and retested within the new range. This helped us find the local maximum in the parameter space. However, we did not allow PC regression to use fewer than 10 PCs. We tested at increments of 0.005 λ_1_ for graphical lasso, 0.1 λ_2_ for graphical ridge, and 5 PCs for PC regression.

### 2.2. Empirical MRI data and processing

Our empirical analyses used the Human Connectome Project in Aging (HCP-A) dataset (Bookheimer et al., 2019; Harms et al., 2018), which is publicly available through the NIMH Data Archive. This extensive dataset includes behavioral measures and high-quality multimodal MRI for 1200+ participants sampled across the adult lifespan (36-100+). The present study used resting-state fMRI to calculate FC, with additional measures used to validate those estimates (SC from diffusion MRI, task fMRI activations, subject age and fluid intelligence scores). Participants were recruited from the areas surrounding the four acquisition sites (Washington University St. Louis, University of Minnesota, Massachusetts General Hospital, and University of California, Los Angeles). All participants gave informed consent through the institutional review board associated with each recruitment site. We excluded from our analyses any participants who were noted to have quality control issues or who were missing any of the MRI scans or behavioral measures that we utilized in this study. This left us with 472 subjects, divided evenly between discovery (n = 236 subjects, 141 females; mean age = 56.9 years, SD = 13.95) and replication (n = 236 subjects, 134 females; mean age = 58.2 years, SD = 14.35) datasets.

Our analyses utilized resting-state fMRI, task fMRI, and diffusion MRI from HCP-A (Harms et al., 2018). Across the four sites, data was collected using a Siemens 3T Prisma scanner. fMRI scans were acquired with TR = 800 ms, 72 slices, and 2.0 mm isotropic voxels. The resting-state fMRI data were collected in four runs over two days, each run containing 488 volumes and lasting 6 min 41 s. For task fMRI data we analyzed the Go/No-go task, which contained 300 volumes and lasted 4 min 11 s. We used the publicly available minimally preprocessed fMRI data, processed using the HCP minimal preprocessing pipeline (Glasser et al., 2013). The preprocessed cortical surface data were parcellated into 360 brain regions according to the multimodal Glasser parcellation (Glasser et al., 2016). We did not apply ICA-FIX denoising in favor of our own nuisance regression procedures, previously described by Ito et al. (2020). We first removed the first 5 frames from each run and demeaned and linearly detrended the timeseries. We then performed nuisance regression as described by Ciric et al. (2017) with 24 motion regressors and 40 physiological noise regressors, the latter being modeled from white matter and ventricle timeseries components using aCompCor (Behzadi et al., 2007). Note that – as in Cole et al. (2021) – aCompCor was used in place of global signal regression, given evidence that it has similar benefits as global signal regression for removing artifacts (Power et al., 2018) but without regressing gray matter signals (mixed with other gray matter signals) from themselves, which may result in false correlations (Murphy et al., 2009; Power, Laumann, et al., 2017). The cleaned resting state data was then partitioned into two sessions of 956 TRs (12 min 45 s), the concatenations of runs 1-2 and runs 3-4. Those two sessions of resting-state data were used to produce two FC matrices per subject for all FC methods (see section 2.1).

The preprocessed task fMRI timeseries were used to estimate task-evoked activations. We used the Go/No go task, in which 92 stimuli (simple geometric shapes) were presented. We estimated the mean activation values separately for “hit”, “correct rejection”, “false alarm”, and “miss” events by fitting a general linear model with 4 separate regressors (canonical hemodynamic response function (HRF) convolved with event onsets). Our analyses only included the “hit” and “correct rejection” regressors, as these were the events where subjects responded correctly. Task regression was performed concurrently with the nuisance regression described above.

Diffusion MRI was used to estimate SC, or the brain’s network of white matter tracts. It measures the diffusion of water molecules in each voxel in different directions. Where the molecules are only able to move freely in certain directions, this typically corresponds with a white matter tract. HCP-A acquired diffusion MRI with 1.5 mm isotropic voxels, sampling 92-93 directions in each of two shells (b = 1500 and 3000 s/mm^2^) and repeating the entire acquisition in both AP and PA phase encoding directions (Harms et al., 2018). This large quantity of data aids in making accurate estimations of SC. We preprocessed the data using the HCP diffusion pipeline (Sotiropoulos et al., 2013). Tractography was performed using DSI Studio (Yeh et al., 2013), as it was shown to be among the most accurate pipelines tested by Maier-Hein et al. (2017). Note that there are a variety of other diffusion MRI structural connectivity approaches, which are known to vary in their relationship with FC (Nelson et al., 2023). We first estimated local fiber orientations using generalized Q-sampling imaging. We then transformed each subject’s individualized cortical parcellation into volume space from surface space to use with DSI Studio. Tractography was performed with a deterministic algorithm, by placing thousands of seeds throughout the white matter volume and letting them spread as streamlines from the initial point, directed through 3-dimensional space by the local fiber orientation vector field. Structural connection weights between each pair of regions were determined as the normalized count of streamlines that overlap with voxels in both regions. These streamlines estimate the presence and size of white matter tracts physically linking the brain areas.

In addition to the neuroimaging data, this study also used out-of-scanner behavioral measures to gauge individual differences in intelligence. We used factor analysis to estimate psychometric *g*, or the general intelligence factor, for each subject (Johnson et al., 2008; McCormick et al., 2022). We analyzed data from all available measures of fluid intelligence (Bookheimer et al., 2019), which consisted of the Picture Sequence Memory Test, Dimensional Change Card Sort Test, Flanker Task Control and Attention Test, Pattern Completion Processing Speed Test, and List Sorting Working Memory Test from the NIH Toolbox (Weintraub et al., 2013), as well as the Rey Auditory Verbal Learning Test (Rey, 1941) and Trail Making Test B (Bowie & Harvey, 2006). Factor analysis was applied to all subjects’ test scores (separately for discovery and replication datasets) to estimate the unitary factor loadings underlying all cognitive measures. The *g* scores were then calculated for each subject, reflecting general intelligence in a single value. We implemented factor analysis using the R package psych (Revelle, 2017).

### 2.3. Simulated networks and timeseries

Our simulation analyses involved creating random network connections, simulating activity timeseries for all nodes using linear modeling, estimating FC on those simulated data using each FC method, and then comparing the resulting FC matrices. The generated networks were directed and asymmetric. They were designed to be modular (organized into communities), small-world (showing high clustering but efficient paths between all nodes), roughly scale-free (containing densely connected hub nodes), and rich-club (having hubs highly connected to each other), because the brain network has been shown to have these characteristics (Bullmore & Sporns, 2009; van den Heuvel & Sporns, 2011).

Networks were comprised of 100 nodes organized into 5 modules of 20 nodes, with edges added based on the Barabási-Albert model (Albert & Barabási, 2002). For each module, we started with 4 connected nodes and added one at a time until reaching the full size of 20. Each node came with 4 attached edges (binary, undirected), which attached to existing nodes with preference for those with high degree. That is, the probability of the new edge attaching to any node *i* over all existing nodes *j* was:

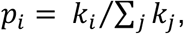

where *k* is node degree, or the number of edges connected to that node. We then added extra-modular edges, adding them one at a time until there were 25% as many edges outside of modules as within modules (6.25% percent the density). The nodes to be connected were again chosen with higher preference given to nodes with high degree, with each of the two nodes having the above probability of being attached. This resulted in all networks having 886 directed (or 443 bidirectional) edges (8.95% of all possible edges; 700 or 36.84% of intra-modular edges; 186 or 2.33% of extra-modular edges). After establishing this binary backbone, we randomly assigned weights to each directed edge from a normal distribution (SD = 1, intra-modular edge mean = 1, extra-modular edge mean = 0.5). One of the five modules was designated as an inhibitory network, its extra-modular edges multiplied by –1. All weights were then normalized by the mean node inputs, and they were scaled by 0.6 to prevent resulting activities from exploding (having extremely large variance). 50 networks were generated for the main analyses and an additional 50 networks were used when testing the impact of the amount of data and noise level.

Activity timeseries were simulated from linear models as described by Sanchez-Romero and Cole (2021) and Sanchez-Romero et al. (2023). The relationships of variables in the network were described by the linear model:

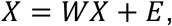

where *X* is a dataset of *p* nodes with *n* datapoints, *E* is a dataset of *p* normally distributed, independent intrinsic noise terms with *n* datapoints, and *W* is the matrix of connectivity weights between all *p* nodes, which must have its diagonal equal to zero. The activities in *X* were calculated by expressing the linear model as:

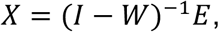

where *I* is the identity matrix. Additional measurement noise, scaled by noise level, was then added to each timeseries. For the main analyses, we simulated 250 datapoints per session and 100 sessions for each of the 50 networks. Each had a noise level of 0.5, meaning that the added measurement noise had roughly half the amplitude of the original signal. When simulating data with varying amounts of data and noise levels, we simulated one session for each combination of 50, 100, 200, 300, 400, 500, 1000, and 10000 datapoints and 0.25, 0.50, and 1.00 noise levels for each of the 50 additional networks. All FC methods were applied to each session of simulated data (see section 2.1).

While we attempted to capture several hypothesized properties of the brain’s functional network, we do not know the absolute true brain network, and some features of the above simulations could potentially bias results. We therefore repeated the main analyses with two supplemental manipulations. First, we generated 25 networks with twice the density of the originals (weights now scaled by 0.65). That is, each module began with 8 fully connected nodes, and each added node had 8 attached edges. We again simulated 100 sessions for each network. Then, we generated another 25 networks like the originals with 100 sessions each, but with an HRF convolution step. Nodes’ intrinsic activities were instead a binary timeseries of “up” and “down” states, created by a Poisson process that determined the probability of switching states (mean 2.5 s per “up” phase and 10 s per “down” phase). Neural noise was then added with a standard deviation of 5% the difference between binary states (Smith et al., 2011). The slower, more prominent activations produced by this method would be less attenuated by HRF convolution, which can smooth over fast fluctuations. The above linear model equations were applied to produce connectivity effects in the neural data, which was then convolved with a canonical HRF. The neural timeseries were initially generated with a sampling rate of 0.05 s but were down-sampled to a TR of 2 s after convolution. Measurement noise terms were then added.

### 2.4. Analyses

#### 2.4.1. Between-session similarity

Our primary measure of repeat reliability was between-session similarity, which we calculated as the Pearson correlation of all edge weights (vectorized upper triangle) between the FC matrices of two sessions. For empirical data, this was calculated between the two sessions’ FC matrices for each subject (n = 236 within-subject pairs), and for simulated data, this was calculated between the first two sessions’ FC matrices for each network (n = 100 within-network pairs). This controls for subject differences in empirical FC. While there may still be some legitimate differences in FC between sessions, we expect this contribution to be relatively small. FC estimated from simulated data should show no between-session differences, as they were generated from the exact same network. A strong correlation indicates a small degree of variability (high reliability), while a weak correlation indicates that noise is obscuring the underlying edge weights.

#### 2.4.2. Intraclass correlation

The intraclass correlation coefficient (ICC) has previously been used in FC studies to quantify repeat reliability (Fiecas et al., 2013; Mahadevan et al., 2021; Mejia et al., 2018). Unlike the other metrics presented here, it is calculated for each edge rather than for each subject. We calculated for each edge one-way random-effects ICC, or ICC(1,1), according to Shrout and Fleiss (1979):

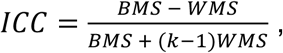

where BMS is the mean squares between subjects (variability in subject-averaged edge weights across the group), WMS is the mean squares within subjects (variability in session edge weights within each subject), and *k* is the number of sessions (k = 2 for this study). BMS should include true variability across subjects plus noise variability, while WMS should primarily contain noise variability.

While intraclass correlation is a generally valid metric, we observed that it is biased against sparse FC methods if applied to all edges. As can be seen from the above formula, ICC is zero when WMS and BMS are equal, which happens when there is only noise and no variability specific to individual differences. When estimating connectivity in the brain, many FC methods yield some null edges that systematically approximate zero for all subjects and only vary from zero due to noise. These null edges would accordingly have ICCs distributed around zero, and sparse FC methods would then have ICCs around zero for the majority of their edges. To avoid this automatic penalty for null edges, we therefore only analyzed edges which were systematically assigned nonzero weights, leaving the potential for between-subject variability. To accommodate a high degree of sparsity, we chose these edges as those which had group-averaged weights in the 98th percentile for every FC method being tested, giving us 543 edges.

#### 2.4.3. Structural-functional similarity

SC was used as a validation benchmark for empirical FC estimates because it offers an independent measure of direct connections between brain regions. SC is not a perfect ground truth for direct FC because strength of white matter tracts does not exactly correspond to functional influence. For instance, FC is also impacted by aggregate effects of microanatomy (e.g., synaptic strengths) that are not captured by diffusion MRI, and FC can fluctuate with cognitive task state or arousal level while SC is static over short timescales. However, given that a direct functional connection can only exist if a structural connection exists as well, the measures are expected to overlap. SC estimated from diffusion MRI is also prone to its own measurement error and does not perfectly represent the white matter network. While diffusion tractography has replicated tract tracing results with above-chance accuracy (Donahue et al., 2016), it has also been shown to produce many false positive and false negative connections (Maier-Hein et al., 2017; Rheault et al., 2020; Sotiropoulos & Zalesky, 2019). However, because SC is calculated from an entirely different MRI modality from FC, it has highly distinct biases and error sources, and it should not be sensitive to the same false connections we wish to test for in FC.

We calculated structural-functional similarity as the Pearson correlation between functional and structural connectivity weights. We included all possible edges, regardless of whether they had zero or nonzero weights in either modality. This was computed from both group-averaged and individual FC to assess validity and individual measurement accuracy, respectively. For validity, the group-averaged session 1 FC matrix was compared with the group-averaged SC matrix. For individual measurement accuracy, we compared each session 1 FC matrix with the SC matrix from the same subject (n = 236 individual matrices). Averaging over many estimates can nullify much of the measurement noise, leaving only the underlying connectivity structures. The group-averaged structural-functional similarity should therefore reflect the validity of each FC method without the influence of reliability, while the individual similarities depend on both.

One characteristic of our SC matrices that could have biased FC method performance is their sparsity. The high degree of sparsity observed here may closely reflect true white matter connectivity, but it is also possible that the tractography method induces greater sparsity than actually occurs in the brain. For instance, deterministic tractography algorithms, such as was used in this study, tend to produce sparser results than probabilistic tractography algorithms. To test whether FC density, regardless of edge accuracy, could be giving some FC methods an advantage in SC similarity, we recalculated the measures after thresholding the FC matrices to match the SC densities. The SC density (percent nonzero edges) was calculated for each subject, and for each FC matrix from the same subject, only the edges with weights above that percentile were left while all others were brought to zero. To avoid large gaps between zero and nonzero values, we also subtracted the minimum nonzero value from all nonzero weights. Group-averaged and individual structural-functional similarity were then recomputed using these artificially sparse FC matrices.

#### 2.4.4. Ground truth similarity

For simulated data, we computed ground truth similarity as the Pearson correlation between each FC matrix and the corresponding ground truth weights matrix. Since all FC methods that we implemented estimate undirected connectivity, we calculated correlations with symmetrized versions of the ground truth matrices. As we did with structural-functional similarity, we computed ground truth similarity using group-averaged as well as individual FC matrices, as averaging reduces the impact of poor reliability to indicate just group-level validity. For the main simulation analyses, we calculated ground truth similarity of group-averaged matrices using FC matrices averaged over 100 sessions for each network (n = 50 group-averaged matrices) and individual matrices using only the session 1 FC matrix from each network (n = 50 individual matrices). When analyzing the effect of amount of data and noise level, we calculated ground truth similarity of individual FC matrices for each network (n = 50 individual matrices) for all conditions.

#### 2.4.5. Correlation with subject motion

Head motion artifacts in fMRI timeseries can lead to spurious, systematic changes in FC estimates, often dependent on the distance between regions (Power et al., 2015). We quantified the influence of subject motion on FC weights using quality control-functional connectivity (QC-FC) correlation, as described by Ciric et al. (2017). QC-FC correlation was calculated for each edge as the partial correlation between subjects’ estimated edge weights and mean relative root mean square (RMS) displacement, conditioned on subject age and sex. We calculated median QC-FC correlation and percentage of edges with significant correlations (p < .05 following false discovery rate correction; Benjamini & Hochberg, 1995) for each FC method. We additionally calculated QC-FC correlation while controlling for FC sparsity. This was done by re-calculating each QC-FC correlation while only including subjects with nonzero weights (|*w*| > .01) for the given edge. Only edges where at least 80% of subjects had nonzero weights were analyzed.

#### 2.4.6. Task activation prediction

Task activation prediction measures how accurately the FC matrices relate to independent functional activity. For this we used the task-evoked activations for “hit” and “correct rejection” events in the Go/No go task. Predictions were generated using activity flow modeling, which simulates the generation of activity in a target node from the activity flows of all other nodes over their connections with the target (Cole et al., 2016). The predicted activity of each target node was calculated as:

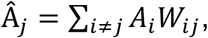

where *A_i_* is the activity of source node *i* and *W_ij_* is the FC coefficient between the target *j* and source node *i*. Then, the similarity of the predicted activity (*Â_j_*) to actual activity (*A_j_*) was quantified using Pearson correlation.

#### 2.4.7. Age and intelligence prediction

Lastly, we tested how well the FC weights from different methods could predict subject age and intelligence (psychometric *g* score). As fluid intelligence often declines with age (Harada et al., 2013; Salthouse, 2010), we first regressed subject age from the *g* scores so that we could examine their relationship with FC independently. We predicted age and age-adjusted *g* scores separately by fitting ridge regression models within 10-fold cross-validation. 9 folds of subjects were used to train each regression model, with FC edge weights (session 1 only) as predictor variables. We used ridge regression because the number of variables (n = 64620 edges) is much larger than the number of observations (n = 212 or 213 subjects in the 9 folds). The hyperparameter values were selected through nested cross-validation by Scikit-learn’s RidgeCV function. We then applied each model to the held-out fold of subjects to calculate their predicted *g* scores and ages. The predicted values were pooled over all folds, and we quantified model accuracy for each FC method as the Pearson correlation between predicted and actual values.

## 3. Results

### 3.1. Visualizing empirical fMRI functional connectivity

Before comparing the different FC methods based on purely quantitative measures, we found it informative to first visualize the resulting FC matrices. We present both group-averaged and individual subject matrices in Figure 2, with the nodes ordered according to the Cole-Anticevic Brain-wide Network Partition to reveal the network architecture (Ji et al., 2019; Figure 2A). For the regularized methods, we measured the mean optimal hyperparameters for both sessions (n = 236 subjects) to be λ_1_ = 0.034 for graphical lasso (SD = 0.006), λ_2_ = 0.404 for graphical ridge (SD = 0.154), and 54.6 PCs for PC regression (SD = 14.9). The group-averaged matrices facilitate comparing the network structures generated by each FC method. Meanwhile, individual subject matrices can expose low measurement reliability if the edge weights vary substantially from the group-averaged weights.

Two contrasting FC methods are pairwise correlation and partial correlation. Pairwise correlation (the field standard) can be seen to produce dense connectivity, as it reflects any linear time series similarity between nodes (Figure 2B). Partial correlation instead creates a sparse network, which is theoretically limited to direct and unconfounded connections (Figure 2C). However, this sparse network structure is obscured by edge weights that vary widely from the group average, possibly reflecting noise. We will investigate this possibility quantitatively in subsequent sections.

We tested several different regularization methods, and displaying their resulting FC matrices illustrates some clear differences between graphical lasso, graphical ridge, and PC regression (Figure 2D-E). Graphical lasso can be seen to produce a similar sparse network structure to partial correlation at the group level. Looking at its individual subject matrix, however, suggests that graphical lasso may produce less edge weight variability. The networks resulting from graphical ridge and PC regression appear similar to each other in that their connectivity graphs are more diffuse than that of graphical lasso, with lower weights spread over more edges. Their structures appear to diverge somewhat from that of partial correlation, which may bode poorly for the methods if partial correlation FC is in fact a valid representation of direct brain connectivity at the group level. Like graphical lasso, graphical ridge and PC regression show less deviation between individual and group-averaged FC.

### 3.2. Reliability of FC with empirical fMRI data

Low repeat reliability has been observed with partial correlation FC (Fiecas et al., 2013; Mahadevan et al., 2021), which may mean that this otherwise effective method is inaccurate at the level of individual measurements. We hypothesized, however, that regularization could solve this problem. We tested the reliability of FC methods in empirical data using between-session similarity and intraclass correlation.

Replicating previously published results, we found partial correlation FC had low between-session similarity (mean r = 0.108, SD = 0.015; Figure 3A) and low ICCs (mean ICC = 0.310, SD = 0.115; Figure 3B). This is especially poor when compared with pairwise correlation FC, which scores significantly higher for both between-session similarity (mean r = 0.804, SD = 0.058; p < .00001) and ICCs (mean ICC = 0.670, SD = 0.053; p < .00001). Our comparison statistics (see Table 1, row 1) confirm that pairwise correlation FC has significantly higher reliability than partial correlation FC.

**Figure 3.**
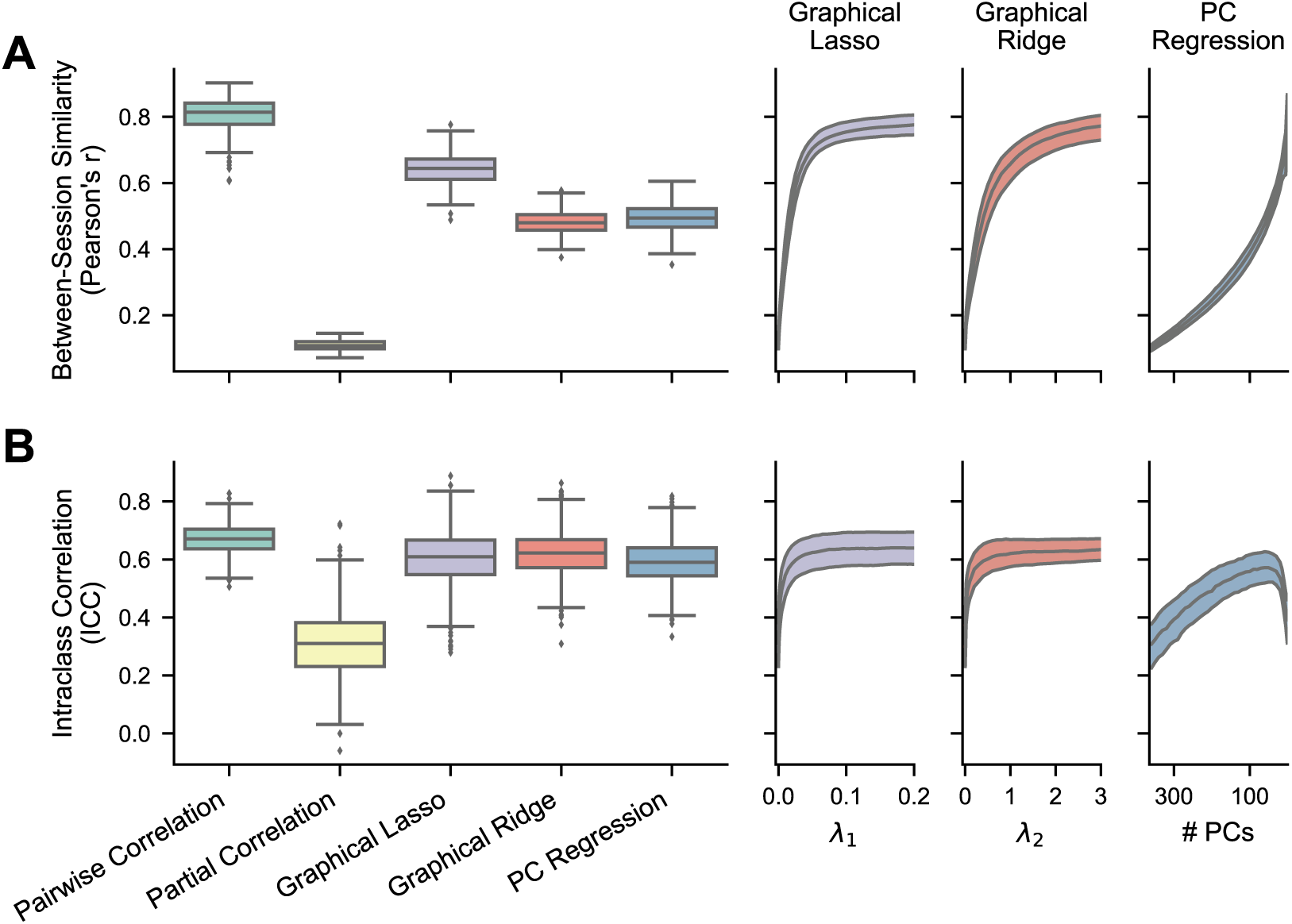
– Reliability of FC methods with empirical fMRI data. The boxplots show results where the regularization hyperparameters have been optimized for each FC matrix, while the right plots show the medians and IQRs across different hyperparameter values for the regularized methods. For PC regression, number of PCs is plotted in descending order because fewer PCs correspond with more regularization. **A)** Between-session similarity, the Pearson correlation between each subject’s session 1 and session 2 FC matrices (n = 236). **B)** Intraclass correlation (Shrout & Fleiss, 1979), calculated for each edge in a conservative subset of nonzero edges (n = 543).

**Table 1.**
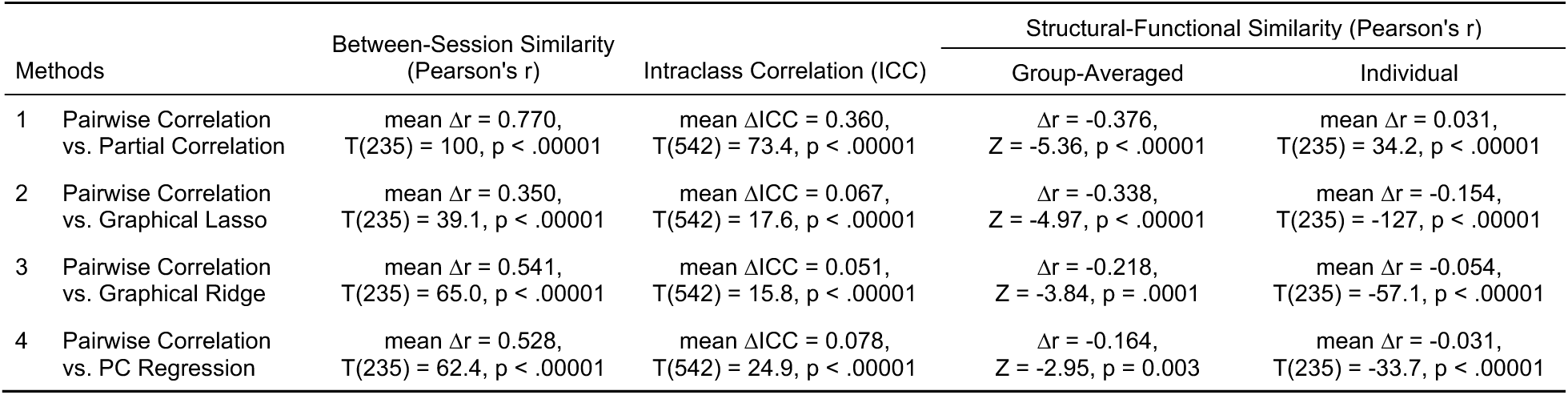
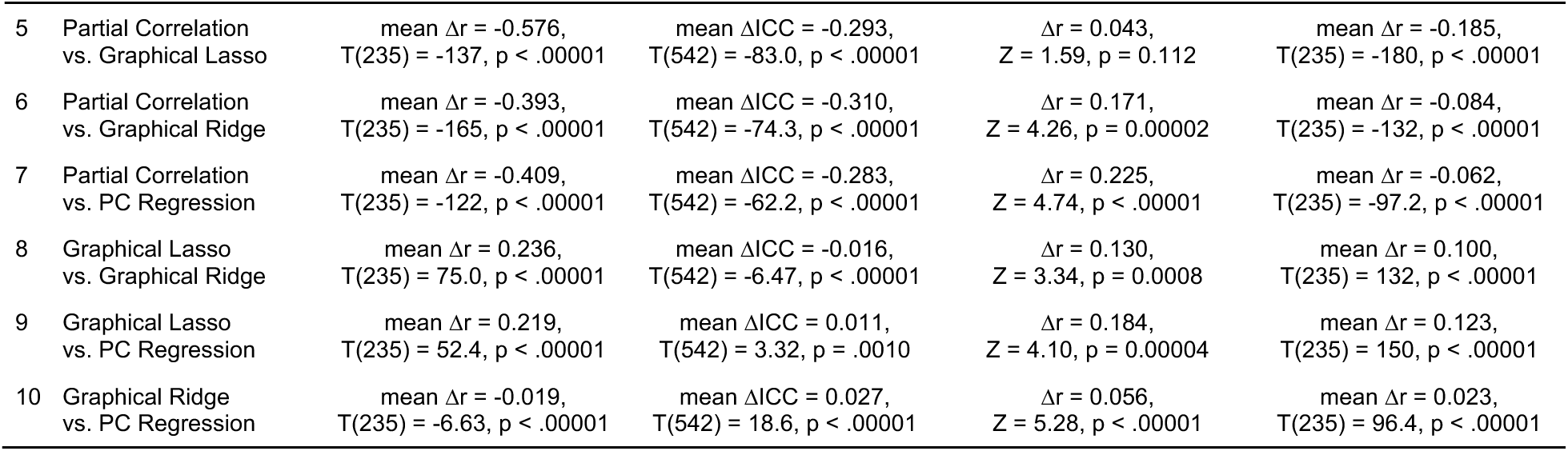
– Statistical tests comparing FC methods on reliability and structural-functional similarity using empirical data. Between-session similarity, intraclass correlation, and individual structural-functional similarity were compared using two-tailed, dependent-sample t-tests. The Pearson’s r values for between-session similarity and individual structural-functional similarity were normalized using Fisher’s z transformation (arctanh) before statistical comparisons. Group-averaged structural-functional similarity (single Pearson’s r per method) was compared using the method described by Meng et al. (1992), which tests for a significant difference between “correlated correlations” (correlations with a shared variable – structural connectivity weights in this case). For all reported correlation differences, correlations were transformed to Fisher’s z, subtracted, and then transformed back to Pearson’s r. Alpha levels were adjusted to .005 (.05/10) following Bonferroni correction to account for the multiple comparisons within each column.

| Methods | Between-Session Similarity<br>(Pearson's $r$) | Intraclass Correlation (ICC) | Structural-Functional Similarity (Pearson's $r$) | |
| --- | --- | --- | --- | --- |
|  |  |  | Group-Averaged | Individual |
| 1 Pairwise Correlation vs. Partial Correlation | mean $\Delta r = 0.770$,<br>$T(235) = 100$, $p < .00001$ | mean $\Delta ICC = 0.360$,<br>$T(542) = 73.4$, $p < .00001$ | $\Delta r = -0.376$,<br>$Z = -5.36$, $p < .00001$ | mean $\Delta r = 0.031$,<br>$T(235) = 34.2$, $p < .00001$ |
| 2 Pairwise Correlation vs. Graphical Lasso | mean $\Delta r = 0.350$,<br>$T(235) = 39.1$, $p < .00001$ | mean $\Delta ICC = 0.067$,<br>$T(542) = 17.6$, $p < .00001$ | $\Delta r = -0.338$,<br>$Z = -4.97$, $p < .00001$ | mean $\Delta r = -0.154$,<br>$T(235) = -127$, $p < .00001$ |
| 3 Pairwise Correlation vs. Graphical Ridge | mean $\Delta r = 0.541$,<br>$T(235) = 65.0$, $p < .00001$ | mean $\Delta ICC = 0.051$,<br>$T(542) = 15.8$, $p < .00001$ | $\Delta r = -0.218$,<br>$Z = -3.84$, $p = .0001$ | mean $\Delta r = -0.054$,<br>$T(235) = -57.1$, $p < .00001$ |
| 4 Pairwise Correlation vs. PC Regression | mean $\Delta r = 0.528$,<br>$T(235) = 62.4$, $p < .00001$ | mean $\Delta ICC = 0.078$,<br>$T(542) = 24.9$, $p < .00001$ | $\Delta r = -0.164$,<br>$Z = -2.95$, $p = 0.003$ | mean $\Delta r = -0.031$,<br>$T(235) = -33.7$, $p < .00001$ |

|  |  |  |  |  |  |
| --- | --- | --- | --- | --- | --- |
| 5 | Partial Correlation<br>vs. Graphical Lasso | mean $\Delta r = -0.576$,<br>T(235) = -137, $p < .00001$ | mean $\Delta ICC = -0.293$,<br>T(542) = -83.0, $p < .00001$ | $\Delta r = 0.043$,<br>Z = 1.59, $p = 0.112$ | mean $\Delta r = -0.185$,<br>T(235) = -180, $p < .00001$ |
| 6 | Partial Correlation<br>vs. Graphical Ridge | mean $\Delta r = -0.393$,<br>T(235) = -165, $p < .00001$ | mean $\Delta ICC = -0.310$,<br>T(542) = -74.3, $p < .00001$ | $\Delta r = 0.171$,<br>Z = 4.26, $p = 0.00002$ | mean $\Delta r = -0.084$,<br>T(235) = -132, $p < .00001$ |
| 7 | Partial Correlation<br>vs. PC Regression | mean $\Delta r = -0.409$,<br>T(235) = -122, $p < .00001$ | mean $\Delta ICC = -0.283$,<br>T(542) = -62.2, $p < .00001$ | $\Delta r = 0.225$,<br>Z = 4.74, $p < .00001$ | mean $\Delta r = -0.062$,<br>T(235) = -97.2, $p < .00001$ |
| 8 | Graphical Lasso<br>vs. Graphical Ridge | mean $\Delta r = 0.236$,<br>T(235) = 75.0, $p < .00001$ | mean $\Delta ICC = -0.016$,<br>T(542) = -6.47, $p < .00001$ | $\Delta r = 0.130$,<br>Z = 3.34, $p = 0.0008$ | mean $\Delta r = 0.100$,<br>T(235) = 132, $p < .00001$ |
| 9 | Graphical Lasso<br>vs. PC Regression | mean $\Delta r = 0.219$,<br>T(235) = 52.4, $p < .00001$ | mean $\Delta ICC = 0.011$,<br>T(542) = 3.32, $p = .0010$ | $\Delta r = 0.184$,<br>Z = 4.10, $p = 0.00004$ | mean $\Delta r = 0.123$,<br>T(235) = 150, $p < .00001$ |
| 10 | Graphical Ridge<br>vs. PC Regression | mean $\Delta r = -0.019$,<br>T(235) = -6.63, $p < .00001$ | mean $\Delta ICC = 0.027$,<br>T(542) = 18.6, $p < .00001$ | $\Delta r = 0.056$,<br>Z = 5.28, $p < .00001$ | mean $\Delta r = 0.023$,<br>T(235) = 96.4, $p < .00001$ |

In accordance with our hypothesis, all three regularized methods significantly improved on the reliability displayed by partial correlation, shown with both between-session similarity (graphical lasso: mean r = 0.642, SD = 0.048; graphical ridge: mean r = 0.480, SD = 0.036; PC regression: mean r = 0.494, SD = 0.040; all p < .00001, Table 1, rows 5-7) and edge ICCs (graphical lasso: mean ICC = 0.603, SD = 0.096; graphical ridge: mean ICC = 0.619, SD = 0.077; PC regression: mean ICC = 0.592, SD = 0.074; all p < .00001, Table 1, rows 5-7). Among the regularized methods, graphical lasso produced the highest between-session similarity (significantly higher than graphical ridge and PC regression; both p < .00001, Table 1, rows 8 and 9), followed by PC regression (significantly higher than graphical ridge; p < .00001, Table 1, last row). Their ICCs were very close, with graphical ridge showing a slight lead (significantly higher than graphical lasso and PC regression; both p < .00001, Table 1, rows 8 and 10) and graphical lasso next highest (significantly higher than PC regression; p = .0010, Table 1, row 9). While the regularized methods did not reach the high scores of pairwise correlation in either between-session similarity or edge ICCs (all p < .00001, Table 1, rows 2-4), they did come much nearer than unregularized partial correlation. Each regularized method could have been made more reliable by implementing a greater degree of regularization, as demonstrated in the right panels of Figure 3A-B. However, we would not recommend optimizing hyperparameters for reliability, as we have found this to trade off against the validity of the resulting FC matrices (see below). All results were also replicated in the replication dataset (Supplementary Figure S1 and Supplementary Table S1).

### 3.3. Similarity of FC to structural connectivity

While it is important for measurement tools to be reliable, it is imperative that they be valid (Figure 1B), meaning that they represent what they are intended to. We approximated the validity and individual measurement accuracy of empirical direct FC by comparing it to an independent estimate of direct brain connections – structural connectivity. SC represents the presence of white matter tracts between regions, and since functional interactions rely on white matter infrastructure, it is expected that FC should largely mirror SC (van den Heuvel et al., 2009). We calculated structural-functional similarity between group-averaged matrices to estimate validity (independent of reliability) and between individual subject matrices to indicate individual measurement accuracy (combined validity and reliability).

We again start by examining pairwise correlation and partial correlation FC. The pairwise correlation FC matrices were not expected to show a high correlation with the SC matrices because pairwise correlation does not distinguish direct from indirect connectivity, producing matrices that are much less sparse. Indeed, pairwise correlation displayed low structural-functional similarity from both its group-averaged (r = 0.250; Figure 4A) and individual matrices (mean r = 0.168, SD = 0.016; Figure 4B). Partial correlation should be expected to show greater similarity because, like SC, it estimates direct connections between brain regions. As expected, the group-averaged partial correlation matrix produced a relatively high degree of structural-functional similarity (r = 0.572) that was significantly greater than that produced by pairwise correlation (p < .00001; Table 1, row 1), reflecting higher validity. However, the individual partial correlation matrices displayed low structural-functional similarity (mean r = 0.137, SD = 0.011), which reflects the impact of noisy connections whose weights were nullified by group averaging. Somewhat surprisingly, the individual similarities for partial correlation were even lower than those from pairwise correlation (p < .00001, Table 1, row 1). This means that, despite partial correlation producing an underlying network structure that is largely valid, the extent of noise renders individual partial correlation FC matrices even less similar to SC than the overly dense pairwise correlation FC. These support the high group-level validity of partial correlation FC for measuring direct connections but also demonstrate how its low reliability can severely undermine the accuracy of individual measurements.

**Figure 4.**
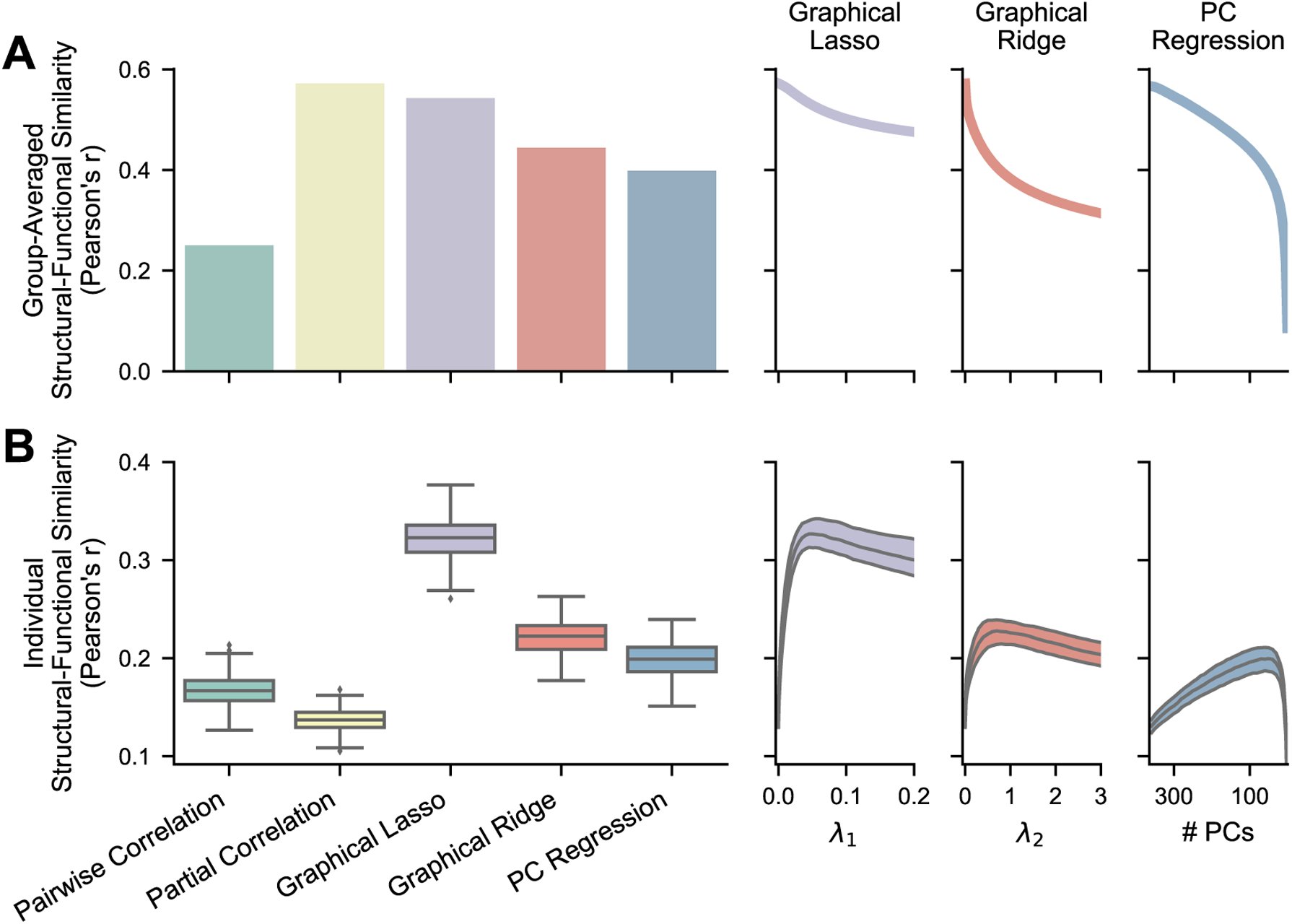
– Similarity between empirical FC and SC (diffusion MRI tractography). The bar and boxplots show results where the regularization hyperparameters have been optimized for each FC matrix, while the right plots show the single values or the medians and IQRs across different hyperparameter values for the regularized methods. For PC regression, number of PCs is plotted in descending order because fewer PCs correspond with more regularization. **A)** Structural-functional similarity between group-averaged FC and structural matrices (n = 1). Averaging nullifies much of the noise in individual connectivity weights to show validity without the effects of low reliability. **B)** Structural-functional similarity between individual FC matrices and the same subjects’ structural matrices (n = 236). Individual measurement accuracy is vastly improved by recovering reliability through regularization.

As expected, the regularized FC methods also showed relatively high levels of structural-functional similarity when using group-averaged matrices (graphical lasso: r = 0.543; graphical ridge: r = 0.444; PC regression: r = 0.398), although unregularized partial correlation scored significantly higher than graphical ridge and PC regression (both p < .00005, Table 1, rows 6 and 7). Graphical lasso did not show a significant difference from partial correlation with group averaging (p = .112, Table 1, row 5). These results indicate that some types of regularization may alter the group-level FC network structures in ways that lessen their validity. This is further illustrated by the decline in group-averaged structural-functional similarity with hyperparameters that induce greater regularization (Figure 4A, right panels). Meanwhile, the regularized FC methods all performed significantly better than basic partial correlation on individual structural-functional similarity (graphical lasso: mean r = 0.322, SD = 0.022; graphical ridge: mean r = 0.222, SD = 0.017; PC regression: mean r = 0.199, SD = 0.017; all p < .00001, Table 1, rows 5-7). Graphical lasso again performed significantly better than graphical ridge and PC regression (both p < .0001, Table 1, rows 8 and 9). This supports our hypothesis that regularization can improve individual measurement accuracy by enhancing reliability. These findings show graphical lasso to be especially promising, as it vastly improves the accuracy of individual FC estimates while preserving high validity of underlying (group-averaged) FC networks seen with partial correlation (Table 1, row 5). Again, these patterns were also demonstrated in held-out subjects (Supplementary Figure S2 and Supplementary Table S1).

In addition, to test whether sparsity alone was driving similarity to SC, we recomputed the same metrics after thresholding each FC matrix (i.e., setting all values under a particular connection strength to 0) to have the same density as that subject’s SC matrix. The average SC density across subjects was 8.83% nonzero edges (SD = 1.026%). The same pattern of results occurred (Supplementary Figure S3 and Supplementary Table S2), except that partial correlation FC, having had many noisy edges brought to zero, now slightly surpassed pairwise correlation FC on individual structural-functional similarity.

### 3.4. Reliability and ground truth similarity of FC with simulated data

Since we do not have access to ground truth empirical FC, we also tested the validity and individual measurement accuracy of FC methods on simulated data whose true underlying connectivity we know. An example simulated network and the FC matrices estimated from a single session are shown in Figure 5. When estimating FC for the regularized methods, the mean optimal hyperparameter values were λ_1_ = 0.112 for graphical lasso (SD = 0.006), λ_2_ = 1.31 for graphical ridge (SD = 0.11), and number of PCs = 11.2 for PC regression (SD = 2.4; number of PCs not allowed to drop below 10). We again quantified the reliability of FC methods using between-session similarity, comparing the first two sessions’ matrices for each of 50 different networks. We evaluated validity using the similarity of network-averaged FC (100 sessions each) with the ground truth networks, and we assessed individual measurement accuracy as the similarity of individual FC estimates (1 session each) with the corresponding true networks.

**Figure 5.**
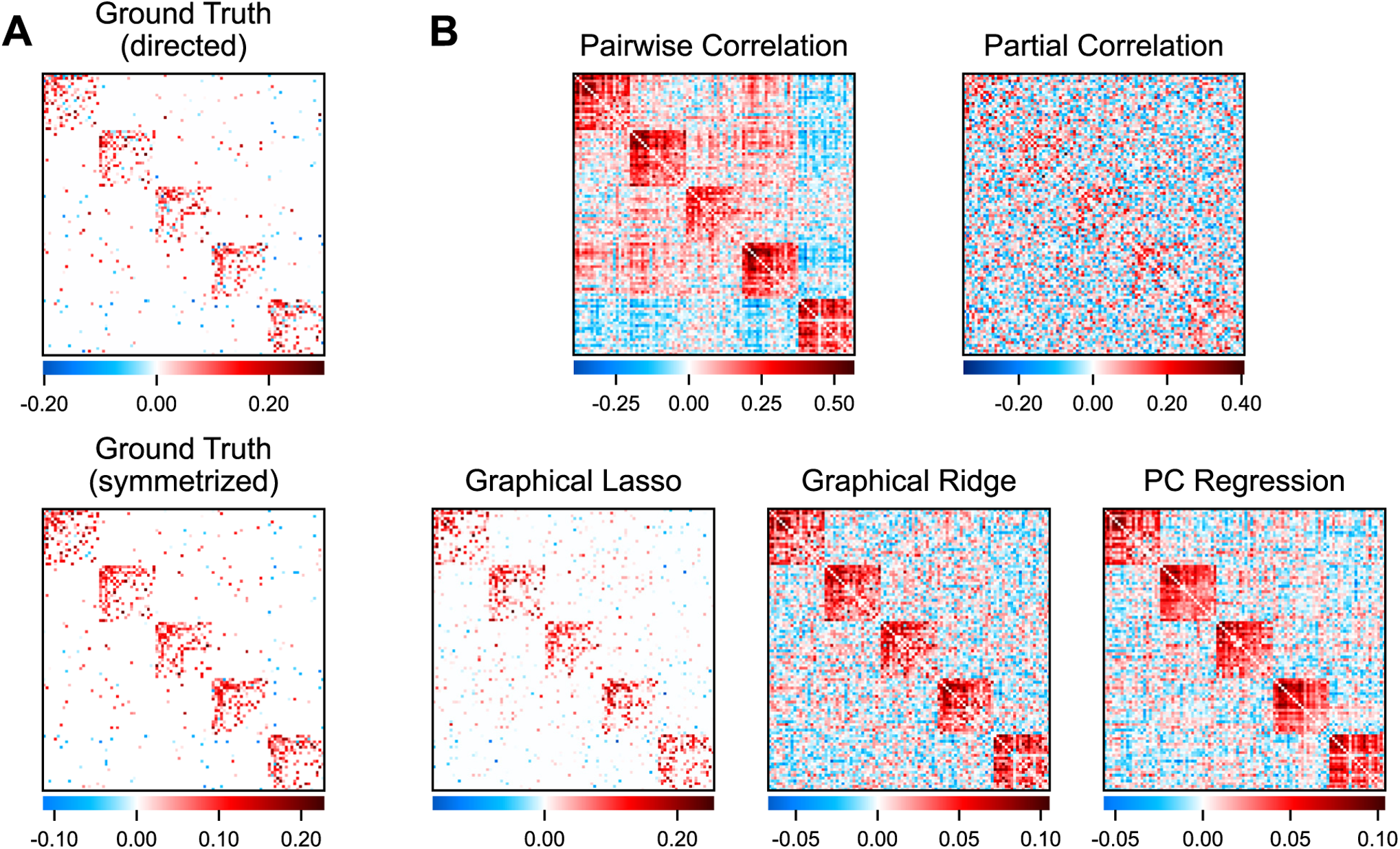
– Simulated networks and FC estimated by different methods. **A)** An example ground truth network. The directed, asymmetric version (top) was used to generate nodes’ timeseries while the symmetrized version (bottom) was compared with FC estimates. **B)** Individual FC matrices estimated from a simulated timeseries using each FC method.

The results using simulated data largely mirror those from empirical fMRI data. In evaluating reliability, partial correlation was again shown to be the least reliable FC method (mean r = 0.158, SD = 0.015; Figure 6A), scoring considerably worse than pairwise correlation (mean r = 0.786, SD = 0.048; p < .00001, Table 2, row 1). Again, all three of the regularized methods improved on the scores of unregularized partial correlation (graphical lasso: mean r = 0.735, SD = 0.018; graphical ridge: mean r = 0.581, SD = 0.024; PC regression: mean r = 0.601, SD = 0.053; all p < .00001, Table 2, rows 5-7). Graphical lasso performed significantly better than graphical ridge and PC regression (both p < .00001, Table 2, rows 8 and 9), and PC regression slightly better than graphical ridge (p = .004, Table 2, row 10). While graphical lasso came close, the regularized methods did not achieve as high reliability as pairwise correlation (all p < .00001, Table 2, rows 2-4). In examining between-session similarity across a range of hyperparameters for the regularized methods, we can again observe that a higher degree of regularization tends to result in greater reliability (Figure 6A, right panels).

**Figure 6.**
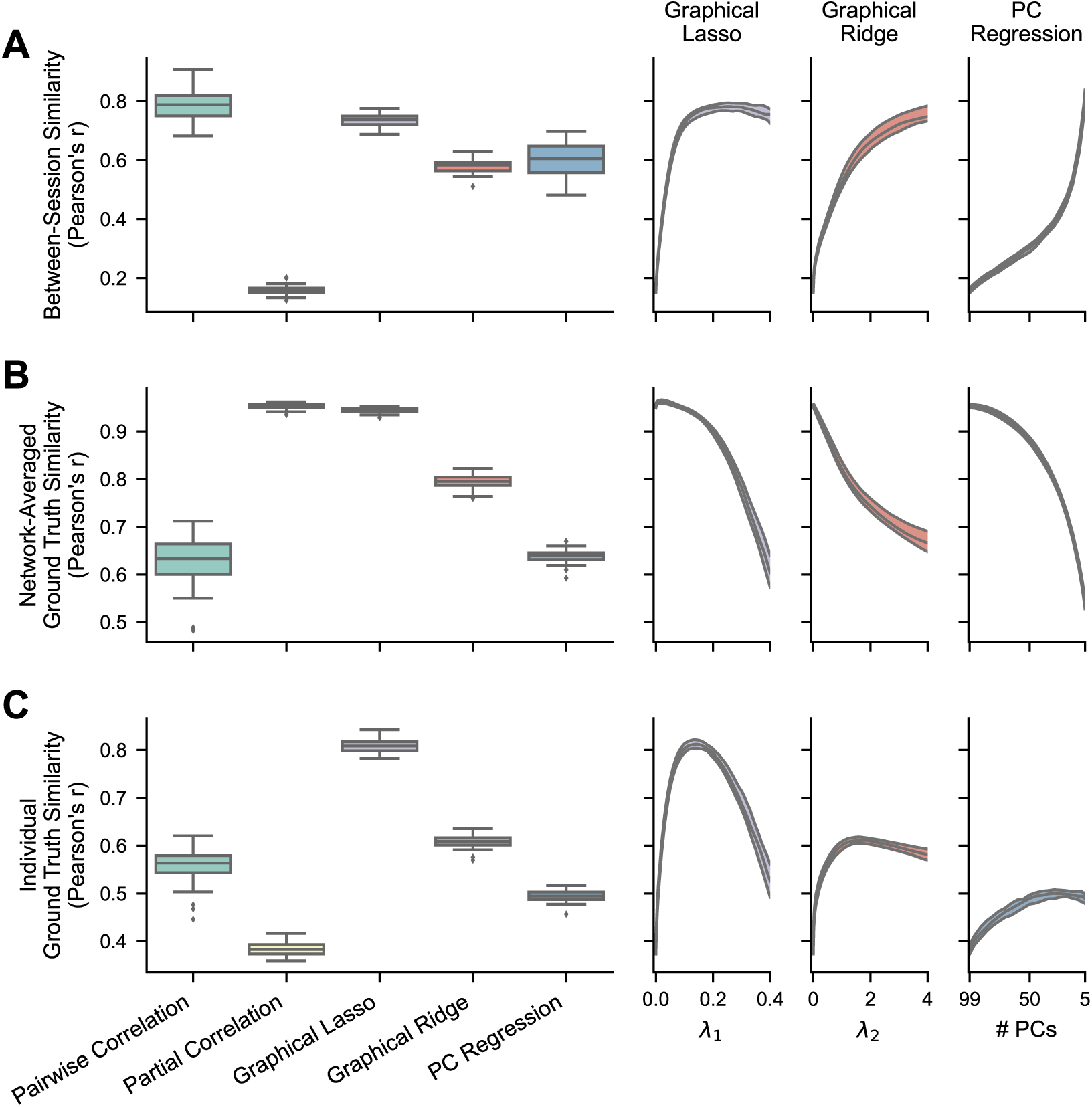
– Reliability and ground truth similarity of FC methods with simulated data. The boxplots show results where the regularization hyperparameters have been optimized for each FC matrix, while the right plots show the medians and IQRs across different hyperparameter values for the regularized methods. For PC regression, number of PCs is plotted in descending order because fewer PCs correspond with more regularization. **A)** Between-session similarity, calculated between one pair of session matrices for each simulated network (n = 50). **B)** Ground truth similarity between group-averaged FC matrices (100 sessions each) and the ground truth for each simulated network (n = 50). Averaging nullifies much of the noise in individual connectivity weights to show validity without the effects of low reliability. **C)** Ground truth similarity between an individual session’s estimated FC matrix and the ground truth for each simulated network (n = 50). The accuracy of single measurements is vastly improved by recovering reliability through regularization.

**Table 2.** – Statistical tests comparing FC methods on reliability and ground truth similarity using simulated data. Between-session similarity, group-averaged ground truth similarity, and individual ground truth similarity were compared using two-tailed, dependent-sample t-tests, their Pearson’s r values first normalized using Fisher’s z transformation (arctanh). For all reported correlation differences, correlations were transformed to Fisher’s z, subtracted, and then transformed back to Pearson’s r. Alpha levels were adjusted to .005 (.05/10) following Bonferroni correction to account for the multiple comparisons within each column.

| Methods | Between-Session Similarity<br>(Pearson's r) | Ground Truth Similarity (Pearson's r) |  |
| --- | --- | --- | --- |
|  |  | Network-Averaged | Individual |
| 1 Pairwise Correlation vs. Partial Correlation | mean $\Delta r = 0.629$,<br>T(49) = 83.3, $p < .00001$ | mean $\Delta r = -0.324$,<br>T(49) = -51.1, $p < .00001$ | mean $\Delta r = 0.172$,<br>T(49) = 37.9, $p < .00001$ |
| 2 Pairwise Correlation vs. Graphical Lasso | mean $\Delta r = 0.051$,<br>T(49) = 6.94, $p < .00001$ | mean $\Delta r = -0.316$,<br>T(49) = -48.5, $p < .00001$ | mean $\Delta r = -0.252$,<br>T(49) = -56.9, $p < .00001$ |
| 3 Pairwise Correlation vs. Graphical Ridge | mean $\Delta r = 0.206$,<br>T(49) = 34.4, $p < .00001$ | mean $\Delta r = -0.166$,<br>T(49) = -31.4, $p < .00001$ | mean $\Delta r = -0.051$,<br>T(49) = -14.1, $p < .00001$ |
| 4 Pairwise Correlation vs. PC Regression | mean $\Delta r = 0.185$,<br>T(49) = 25.7, $p < .00001$ | mean $\Delta r = -0.009$,<br>T(49) = -1.70, $p = 0.096$ | mean $\Delta r = 0.061$,<br>T(49) = 14.6, $p < .00001$ |
| 5 Partial Correlation vs. Graphical Lasso | mean $\Delta r = -0.577$,<br>T(49) = -204, $p < .00001$ | mean $\Delta r = 0.008$,<br>T(49) = 19.9, $p < .00001$ | mean $\Delta r = -0.425$,<br>T(49) = -222, $p < .00001$ |
| 6 Partial Correlation vs. Graphical Ridge | mean $\Delta r = -0.423$,<br>T(49) = -102, $p < .00001$ | mean $\Delta r = 0.158$,<br>T(49) = 101, $p < .00001$ | mean $\Delta r = -0.224$,<br>T(49) = -126, $p < .00001$ |
| 7 Partial Correlation vs. PC Regression | mean $\Delta r = -0.443$,<br>T(49) = -53.7, $p < .00001$ | mean $\Delta r = 0.315$,<br>T(49) = 221, $p < .00001$ | mean $\Delta r = -0.111$,<br>T(49) = -43.8, $p < .00001$ |
| 8 Graphical Lasso vs. Graphical Ridge | mean $\Delta r = 0.154$,<br>T(49) = 42.8, $p < .00001$ | mean $\Delta r = 0.150$,<br>T(49) = 85.1, $p < .00001$ | mean $\Delta r = 0.201$,<br>T(49) = 129, $p < .00001$ |
| 9 Graphical Lasso vs. PC Regression | mean $\Delta r = 0.134$,<br>T(49) = 16.4, $p < .00001$ | mean $\Delta r = 0.307$,<br>T(49) = 196, $p < .00001$ | mean $\Delta r = 0.313$,<br>T(49) = 132, $p < .00001$ |
| 10 Graphical Ridge vs. PC Regression | mean $\Delta r = -0.020$,<br>T(49) = -3.03, $p = 0.004$ | mean $\Delta r = 0.157$,<br>T(49) = 113, $p < .00001$ | mean $\Delta r = 0.112$,<br>T(49) = 82.9, $p < .00001$ |

Meanwhile, partial correlation demonstrated its worth in our test of validity (mean r = 0.952, SD = 0.005; Figure 6B), scoring significantly higher than pairwise correlation on network-averaged ground truth similarity (mean r = 0.628, SD = 0.048; p < .00001, Table 2, row 1). Unregularized partial correlation also scored better than graphical lasso (mean r = 0.944, SD =0.005), graphical ridge (mean r = 0.794, SD = 0.015), and PC regression (mean r = 0.637, SD = 0.013; all p < .00001, Table 2, rows 5-7), although the difference from graphical lasso was very slight. Among the regularized methods, graphical lasso was followed by graphical ridge and then PC regression (all p < .00001, Table 2, rows 8-10). Graphical lasso and graphical ridge achieved greater validity than pairwise correlation (both p < .00001, Table 2, rows 2 and 3), but PC regression was not significantly different (p = .096, Table 2, row 4). In a way, regularization has an opposite impact on validity from reliability, with group-level validity generally decreasing with greater levels of regularization (Fig. 6B right panels). However, this effect was minimal for graphical lasso at the optimal hyperparameter values.

The accuracy of individual FC estimates depends on the underlying estimated network structures being correct (validity) and the individual estimates being minimally perturbed by noise (reliability). Partial correlation produced diminished individual ground truth similarity scores (mean r = 0.384, SD = 0.013; Figure 6C), attributable to its low reliability. These were even lower than those of pairwise correlation (mean r = 0.556, SD = 0.034; p < .00001, Table 2, row 1). However, in improving reliability, all regularized methods were able to increase individual measurement accuracy above that of unregularized partial correlation (graphical lasso: mean r = 0.808, SD = 0.014; graphical ridge: mean r = 0.607, SD = 0.012; PC regression: mean r = 0.495, SD = 0.012; p < .00001, Table 2, rows 5-7). Graphical lasso showed superior accuracy among regularized methods, which is expected from its higher reliability and validity (all p < .00001, Table 2, rows 8 and 9). Graphical lasso and graphical ridge also performed expectedly better than pairwise correlation (both p < .00001, Table 2, rows 2 and 3), but PC regression showed significantly worse accuracy than pairwise correlation (p < .00001; Table 2, row 4). Evaluating their performances at different hyperparameter values, we can see that the optimal levels of regularization are those that balance validity and reliability (Figure 6C right panels). These findings corroborate the results of structural-functional similarity in empirical fMRI data that regularized FC methods can improve individual measurement accuracy. They also substantiate graphical lasso as the most effective of the regularized methods we tested.

We further considered whether properties of the generated networks and simulations could have biased the relative performances of FC methods, so we repeated the analyses with two manipulations in 25 networks each. First, in case we assumed an overly sparse brain network that disproportionally benefitted the sparser FC methods, we generated ground truth networks with twice the density as above (see Figure S4 for an example true network and FC estimates). These denser networks yielded a similar pattern of results (Figure S5; see Table S3 for statistical comparisons). The denser FC methods (graphical ridge, PC regression, and pairwise correlation) may here be closer in performance to the sparser methods (graphical lasso or partial correlation), with PC regression now showing higher reliability than graphical lasso (p = .0005, Table S3, row 9) and graphical ridge showing now significant difference in reliability from graphical lasso (p = .060, Table S3, row 8). There may be a smaller gap between graphical lasso and the denser FC methods in individual measurement accuracy, but graphical lasso still shows the highest ground truth similarity (all p < .00001, Table S3, rows 2, 8, and 9). Second, as the above simulated timeseries were generated with a relatively simple linear model, we included HRF convolution to add an autocorrelation element and make them more resemble BOLD data (see Figure S6 for an example true network and FC estimates). This decreased the performances of all methods somewhat, but their performances relative to each other were unaffected (Figure S7; see Table S4 for statistical comparisons). Graphical lasso remained the strongest contender after these two manipulations as well.

### 3.5. Accuracy of FC with varying amounts of data and noise levels

After confirming that regularization can indeed improve the reliability and individual measurement accuracy of FC, we proceeded to test its benefits to datasets of varied quality. Two factors that can exacerbate model overfitting are low number of datapoints and high degree of noise relative to the signal of interest (Blum et al., 2020; Hastie et al., 2009; Ying, 2019), which remain limitations of fMRI data. We therefore generated 50 additional networks and simulated datasets with differing numbers of timepoints and noise levels, covering a range of low to improbably large amounts of data (0.5, 1-5, 10, and 100 times as many observations as nodes) and three plausible ratios of Gaussian noise relative to the signal (0.25, 0.50, and 1.00). We then computed the FC from each dataset and calculated the ground truth similarities of individual FC estimates.

Our first observation on estimating the FC matrices was that the optimal hyperparameters for the regularized methods vary across conditions, the methods generally preferring a greater degree of regularization when the data contains more noise or fewer timepoints (Figure 7A). For graphical lasso, the average optimal hyperparameter for each condition was strongly correlated with the number of timepoints (Spearman correlation; r_s_ = –0.990, p < .00001) but not with noise level (r_s_ = 0.133, p = .536). For graphical ridge and PC regression, optimal hyperparameter value had a significant monotonic relationship with both number of timepoints (graphical ridge: r_s_ = –0.567, p = .004; PC regression: r_s_ = 0.734, p = .00004) and noise level (graphical ridge: r_s_ = 0.442, p = .030; PC regression: r_s_ = –0.430, p = .036), even though graphical ridge shows a perturbation at 100 timepoints. This indicates a stronger relevance of regularization when datasets have higher noise or fewer timepoints. Note too that this dependence of hyperparameters on timepoints can have practical implications for model selection, necessitating that cross-validation schemes use a similar number of timepoints in training models as will be present in the final model.

**Figure 7.**
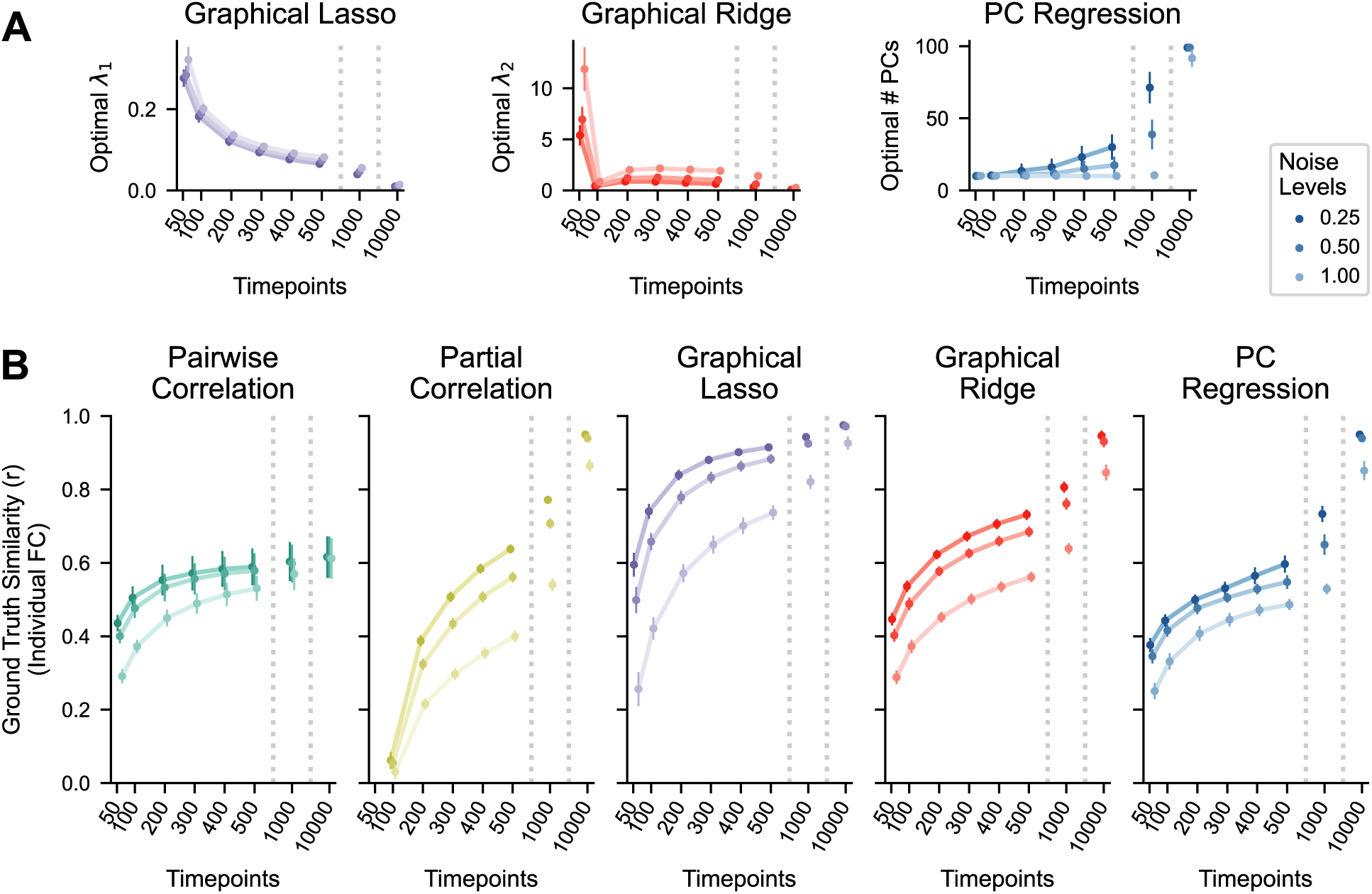
– Performance of FC methods when varying the amounts of data and noise levels in simulated timeseries. **A**) Optimal regularization hyperparameters of each regularized method with different numbers of timepoints (x-axis) and noise ratios (line color). Generally, a larger degree of regularization is preferred when there is more noise or fewer timepoints. For PC regression, number of PCs is plotted in descending order because fewer PCs correspond with more regularization. **B)** Ground truth similarity of individual FC estimates (n = 50; 100 nodes) for different numbers of timepoints (x-axis) and noise ratios (line color). Plots indicate mean and standard deviation.

The extended results on ground truth similarity largely agree with the previous findings but provide greater insights into the methods and their utility to different datasets (Figure 7B; see Table 3 for statistical comparisons). We can see that pairwise correlation maintains relatively stable ground truth similarity across all conditions, showing less susceptibility to low data and high noise but plateauing at a meager level of performance. In contrast, partial correlation performs dismally in low data and high noise conditions, unable to even run with fewer timepoints than variables, but it does successfully recreate the networks when given sufficient data. The amounts of data needed to accomplish this, however, are not often feasible. By adding L1 regularization, graphical lasso can produce strong performances with far fewer timepoints, although results are still limited in cases with high noise and/or very few timepoints. Its improvements over partial correlation were especially large with fewer timepoints, but it still offers marginal gains at 10000 timepoints. Graphical ridge and PC regression are also less susceptible to low data and high noise, although partial correlation surpasses them when more data is available. They also never reach the efficacy of graphical lasso, which scores significantly better than all FC methods across all conditions but that with the least data and most noise (all p < .00001, Table 3, rows 2, 5, 8, and 9). These results again lead us to recommend graphical lasso for FC estimation, for most scan lengths and noise levels. Other solutions may be needed for very short and noisy scans.

**Table 3.**
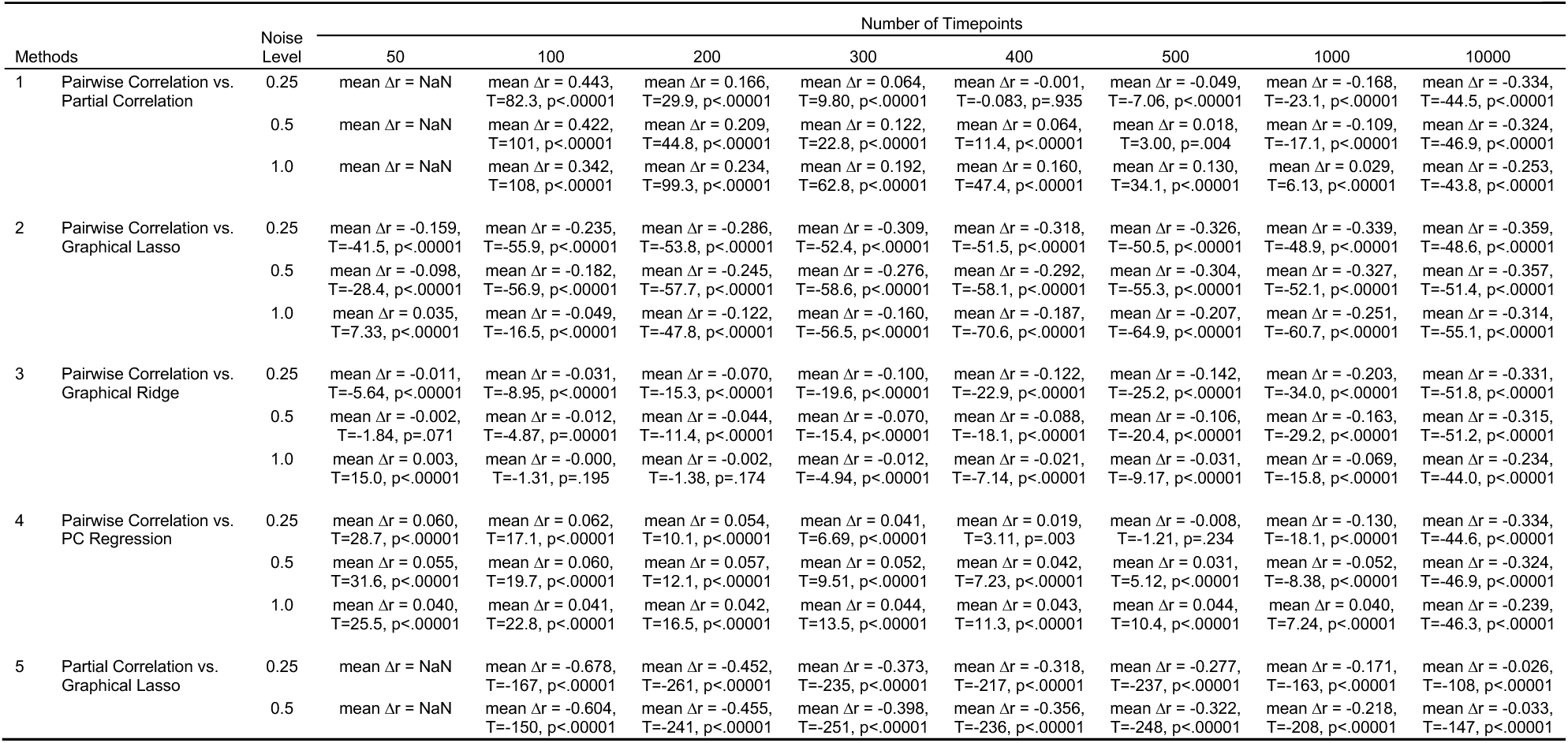
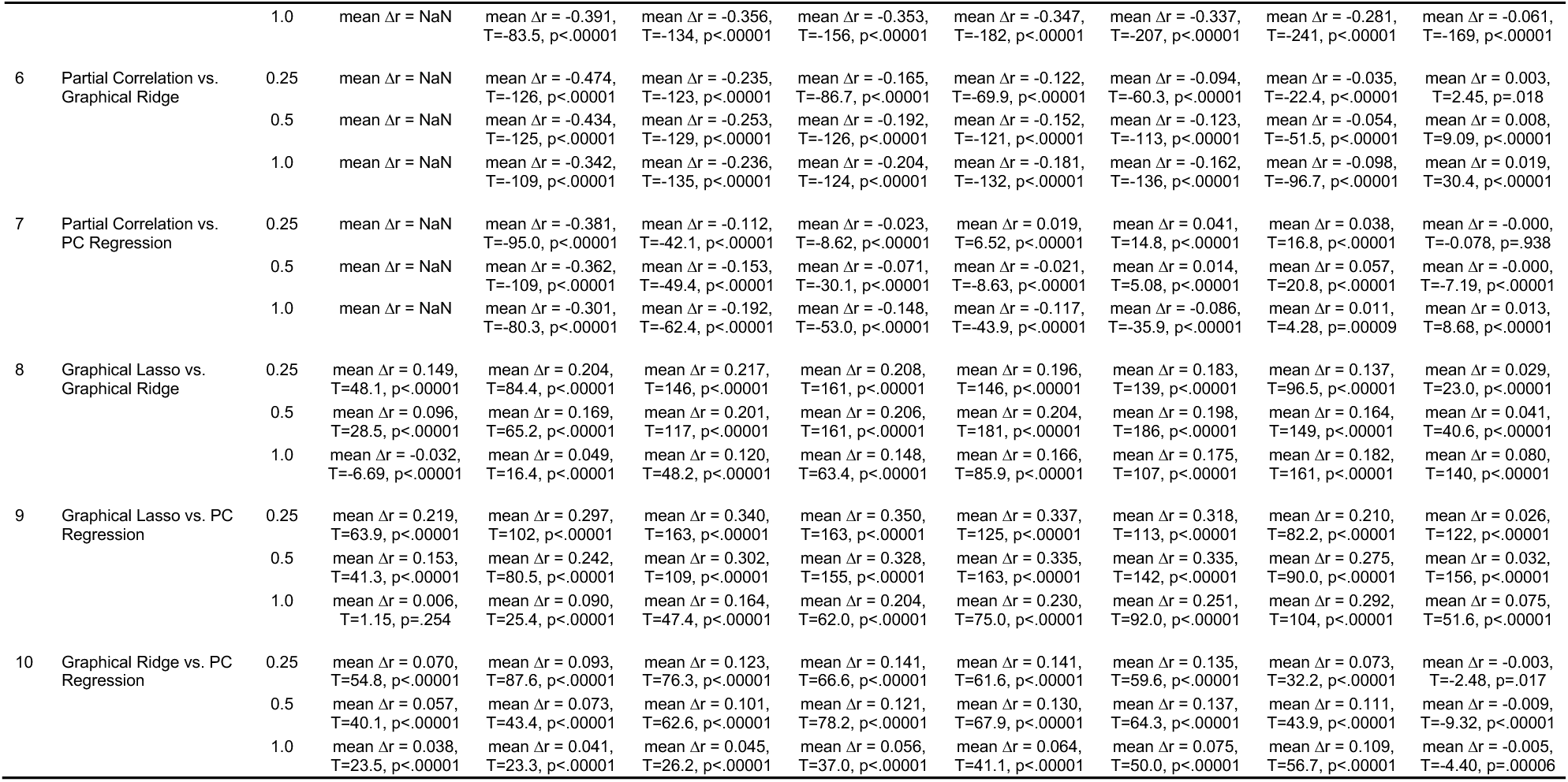
– Statistical tests comparing FC methods on ground truth similarity of individual FC estimates (Pearson’s r) using simulated data with varying amounts of data and noise levels. Individual ground truth similarities were compared using two-tailed, dependent-sample t-tests (49 degrees of freedom), their Pearson’s r values first normalized using Fisher’s z transformation (arctanh). For all reported correlation differences, correlations were transformed to Fisher’s z, subtracted, and then transformed back to Pearson’s r. Alpha levels were adjusted to .00022 (.05/228) following Bonferroni correction to account for the multiple comparisons across FC methods and dataset conditions. Note that NaNs are listed for unregularized partial correlation for the 50 timepoint case, as unregularized partial correlation requires more timepoints than nodes.

### 3.6. Sensitivity of FC to motion artifacts

In addition to the random measurement noise whose impact we tested above, fMRI data is also contaminated with subject motion artifacts (Power et al., 2012). Previous studies have shown partial correlation to effectively mitigate the confounding effects of motion (Mahadevan et al., 2021; Power et al., 2015), but given the pervasiveness of the issue, we sought to also assess the regularized methods tested here. We did so by measuring the QC-FC correlation of each edge, or the correlation across subjects between their mean head motion during scanning and their estimated connectivity weights for each edge of interest (Ciric et al., 2017; Mahadevan et al., 2021).

Our findings replicate those of previous studies, showing partial correlation to be largely robust to motion artifacts with 0% of edges significantly correlated with motion (median absolute QC-FC = 0.045). Pairwise correlation was very susceptible to motion artifacts with 56.4% significant edges (median absolute QC-FC = 0.157), also in agreement with prior findings. Graphical lasso (0.01% significant edges, median absolute QC-FC = 0.048), graphical ridge (0.05% significant edges, median absolute QC-FC = 0.050), and PC regression (0.09% significant edges, median absolute QC-FC = 0.052) were impacted slightly more than unregularized partial correlation, but nonetheless they were also largely insensitive to motion. In case sparsity alone was providing some FC methods with an advantage, we repeated the analyses while only assessing nonzero edges (excluding subjects with very small weights from edge analyses and entire edges with too few remaining subjects; see Methods). The results showed minimal changes, with pairwise correlation having 55.6% significant edges (n = all 64620 edges included in the analysis, median absolute QC-FC = 0.157), partial correlation 0% significant edges (n = 60169, median absolute QC-FC = 0.049), graphical lasso 3.2% significant edges (n = 717, median absolute QC-FC = 0.060), graphical ridge 2.0% significant edges (n = 2771, median absolute QC-FC = 0.065), and PC regression 2.1% significant edges (n = 2995, median absolute QC-FC = 0.065). Their insensitivity to motion artifacts further supports the suitability of regularized partial correlation or similar methods for FC estimation.

### 3.7. Predicting task activations from empirical FC

The remaining analyses further test the accuracy of regularized multivariate FC in representing brain networks and test their advantage for FC-based neuroscience applications. For simplicity, we limited our main results to pairwise correlation, partial correlation, and graphical lasso FC, but the results of graphical ridge and PC regression are presented in Supplementary Figures S8 and S9 and Supplementary Tables S5 and S3, with graphical ridge and PC regression achieving similar performances to graphical lasso.

Here we implemented activity flow modeling to predict held-out neurocognitive function (task activations) from FC matrices (Cole et al., 2016). This model formalizes a functional role for FC, and in this application conveys how well different FC methods captured communication pathways underlying activity flow in the brain. To perform activity flow mapping, the activity of a target node *j* is predicted as the sum of activity from all other source nodes *i*, weighted by the FC coefficients between sources and target (Figure 8A; see Methods). We applied this to the “hit” and “correct rejection” events of a Go/No-go task, and we present the actual and predicted activations for a single subject’s “hit” trials in Figure 8C. The prediction accuracies were calculated for each subject as the Pearson correlation between the predicted and actual activities.

**Figure 8.**
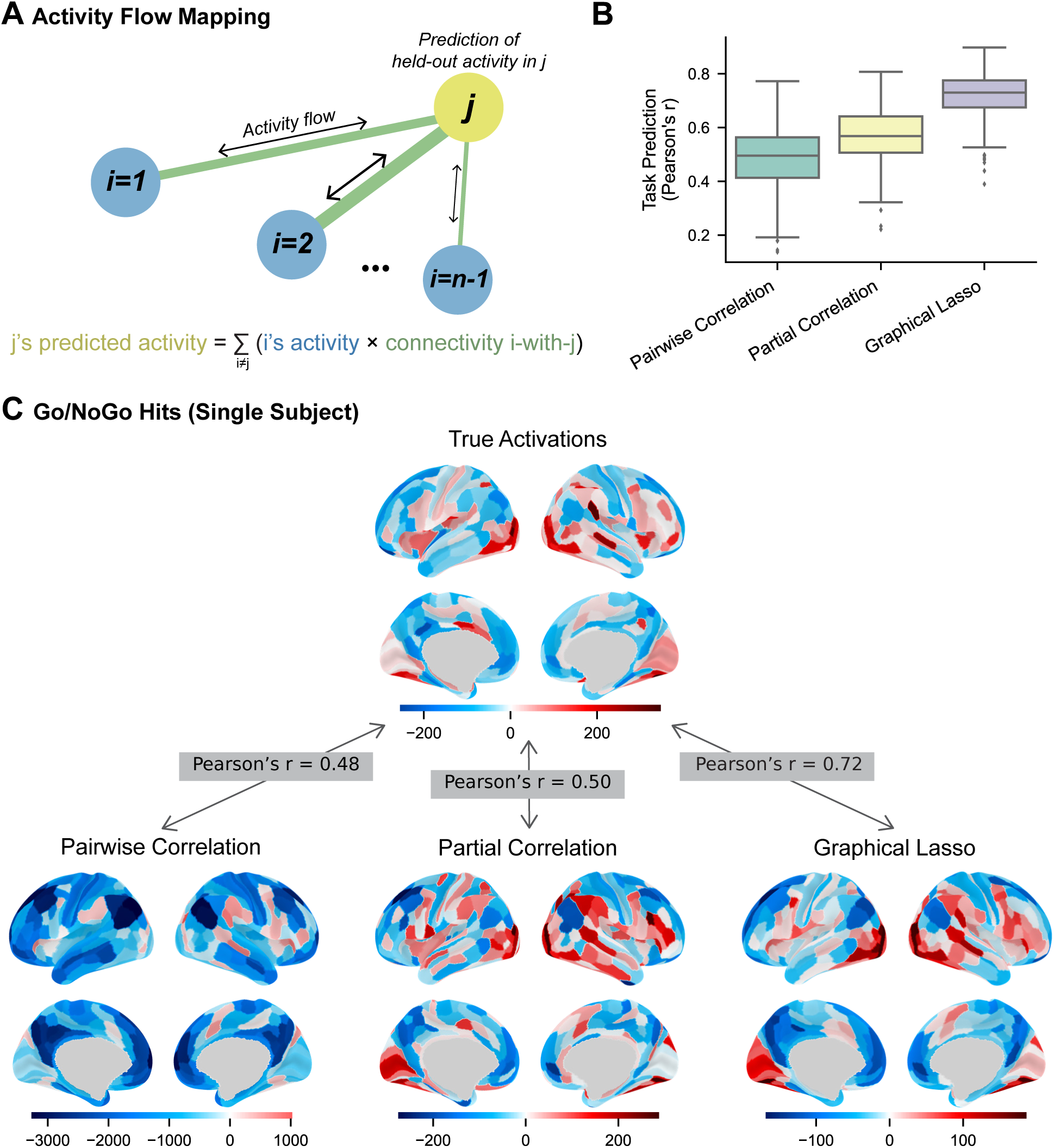
– Predicting task activations using FC estimated from rest fMRI data. **A)** The activity flow mapping paradigm (Cole et al., 2016). Each target region’s (*j*) activity is predicted as the sum of all other regions’ (*i*) activity flows to the target region, which are the products of each source region’s connectivity with the target and the source region’s actual activity. **B)** Prediction accuracy by FC method calculated as the Pearson correlation (r) between regions’ actual and predicted activations, computed in each subject (n = 236) across regions and task conditions. **C)** A single subject’s actual and predicted task activations for the go/no-go hit events.

Pairwise correlation FC generated a mean task prediction score of r = 0.488 (SD = 0.119; Figure 8B). Diverging from the previous tests of FC accuracy, partial correlation produced significantly higher scores than pairwise correlation (mean r = 0.569, SD = 0.106; T(235) = 16.4, p < .00001; all tests two-tailed, dependent-samples; α = .017 (.05/3) for all tests following Bonferroni correction for multiple comparisons). This performance metric appears to be less impacted by low reliability than previous tests, likely because some effects of noisy edges can cancel during the summation of activity flows. These results again support the higher validity of partial correlation than pairwise correlation FC, as the network created by partial correlation better reflects the integration of brain activity over nodes. Activity prediction was further improved by adding regularization through graphical lasso (mean r = 0.719, SD = 0.086), which performed significantly better than both pairwise correlation (T(235) = 48.9, p < .00001) and partial correlation (T(235) = 60.4, p < .00001; α = .017). This pattern of results was reproduced in the replication dataset as well (Supplementary Figure S8 and Supplementary Table S5). Graphical lasso was previously shown to increase reliability while preserving the validity of the underlying estimated network structure, allowing for greater accuracy of individual FC estimates. This improved accuracy allowed for better simulations of activity flowing across brain networks.

### 3.8. Predicting subject age and intelligence from empirical FC

An increasingly common goal in neuroscience research is to link individual differences in brain structure or function with cognitive or other biological characteristics. This can help reveal the functional relevance of brain network organization. However, individual difference studies have been shown to have small effect sizes (Marek et al., 2022), and they therefore may benefit from regularized FC methods that can reduce the impact of measurement noise. If edge weights experience high noise variability compared to true individual variability, then any real individual difference patterns will be obscured, and results will be too easily swayed by random chance. We therefore hypothesized that using more reliable FC estimates would lead to better predictions of individual differences. We tested this by using FC edge weights to predict subject age and intelligence (psychometric *g*; Johnson et al., 2008), comparing prediction accuracies across FC methods. These tests demonstrate the utility of regularization to this research approach while providing additional validation of these methods, confirming their ability to capture the patterns of brain function that underly cognitive differences.

We did indeed find that the more reliable FC methods, pairwise correlation and graphical lasso, predicted individual differences better than the less reliable method, partial correlation. All methods allowed for above-chance prediction of subject age (pairwise correlation: r = 0.739, p < .00001; partial correlation: r = 0.512, p < .00001; graphical lasso: r = 0.695, p < .00001; α = .017 (.05/3) for all tests following Bonferroni correction for multiple comparisons; Figure 9A). Pairwise correlation and graphical lasso both produced significantly higher predicted-to-actual age correlations than partial correlation (pairwise vs. partial correlation: z = 5.34, p < .00001; graphical lasso vs. partial correlation: z = 5.45, p < .00001; α = .017), but their correlations were not significantly different from each other (pairwise correlation vs. graphical lasso: z = 1.59, p = .112; statistical tests described by (Meng et al., 1992). These results were reproduced in the replication dataset (pairwise correlation: r = 0.760, p < .00001; partial correlation: r = 0.579, p < .00001; graphical lasso: r = 0.737, p < .00001; pairwise vs. partial correlation: z = 5.00, p < .00001; graphical lasso vs. partial correlation: z = 5.94, p < .00001; pairwise correlation vs. graphical lasso: z = 0.922, p = .356; α = .017).

**Figure 9.**
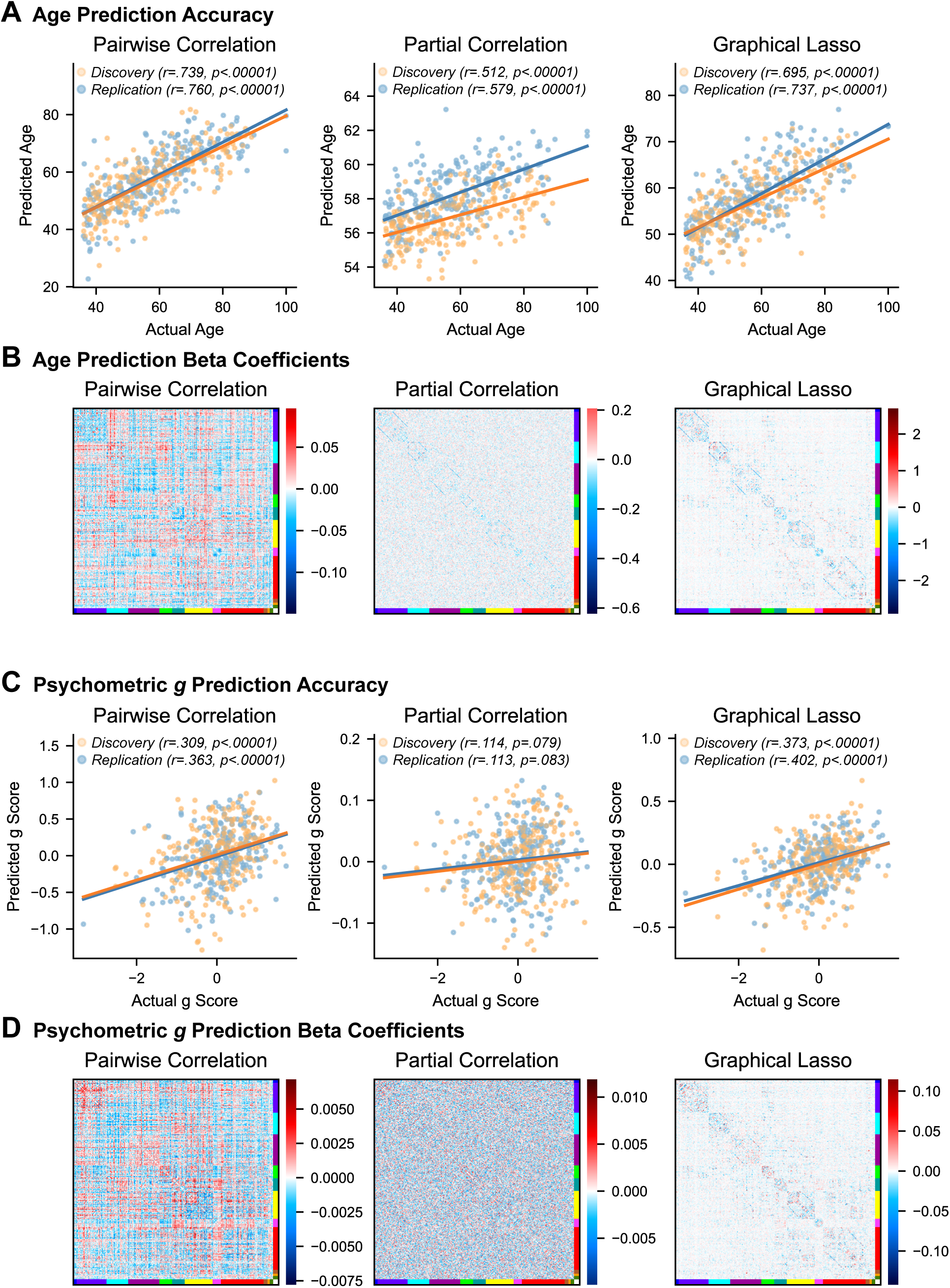
– Predicting individual differences in age and intelligence using estimated FC from rest fMRI data. **A)** Actual and predicted ages of each subject by FC method, from both the discovery and replication datasets. **B)** Average beta coefficients assigned to each connection by the regression models for estimating subject age. Beta coefficients shown here are the averages over cross-validation folds and discovery and replication datasets. Blue indicates that connection strength decreased with age and red indicates that connection strength increased. **C)** Actual and predicted intelligence (psychometric *g*) of each subject by FC method. **D)** Average beta coefficients assigned to each connection by the regression models for estimating psychometric *g*.

The same pattern was observed when predicting intelligence from FC, although prediction accuracies were generally lower than for age, and partial correlation no longer showed a significantly above-chance prediction accuracy (pairwise correlation: r = 0.309, p < .00001; partial correlation: r = 0.114, p = .079; graphical lasso: r = 0.373, p < .00001; α = .017). Again, pairwise correlation and graphical lasso produced significantly better prediction accuracies than partial correlation (pairwise vs. partial correlation: z = 2.48, p = .013; graphical lasso vs. partial correlation: z = 4.08, p = .00004; α = .017), but again their performances were not significantly different from each other (pairwise correlation vs. graphical lasso: z = –1.05, p = .292). The same occurred in the replication dataset (pairwise correlation: r = 0.363, p < .00001; partial correlation: r = 0.113, p = .083; graphical lasso: r = 0.402, p < .00001; pairwise vs. partial correlation: z = 3.16, p = .002; graphical lasso vs. partial correlation: z = 4.43, p < .00001; pairwise correlation vs. graphical lasso: z = –0.639, p = .523; α = .017). Together, these results indicate that regularization improves the ability of partial correlation FC to reflect individual differences in cognition, and they provide additional evidence that graphical lasso FC accurately represents brain connectivity. Further, these results indicate the importance of the increased reliability of regularized partial correlation for matching the behavioral prediction accuracy of pairwise correlation, but now with more interpretable (i.e., valid) connectivity due to the substantial reduction in the number of confounded functional connections.

## 4. Discussion

Accurate estimation of human brain connectivity is a major goal of neuroscience, given evidence that connectivity is a major determinant of neurocognitive function (Bassett & Sporns, 2017). The current field standard approach for FC estimation – pairwise Pearson correlation – is well known to be sensitive to confounded and indirect connections that often do not reflect the brain’s direct functional interactions (Honey et al., 2009; Sanchez-Romero & Cole, 2021; Smith et al., 2011). Direct interactions can offer more mechanistic, causally valid insights into brain communication and network structure (Reid et al., 2019). Therefore, the field may benefit from shifting to a method that removes many of these false connections, such as partial correlation and related multivariate methods. Some other FC methods attempt to provide still stronger causal inferences, for instance by estimating asymmetric directionalities of causal interactions (i.e., *A* → *B* ≠ *A* ← *B*; Sanchez-Romero et al., 2023), but they often require complex models and strong assumptions about the system being studied (Mill et al., 2017; Ramsey et al., 2010; Sanchez-Romero et al., 2019). While not perfect measures of causality, partial correlation and multiple regression FC can offer substantial advances in understanding how brain regions interact, and they do so using comparatively simple, data-driven models. The potential utility of partial correlation is reduced, however, by its low reliability, reported to be much lower than that of pairwise correlation FC (Fiecas et al., 2013; Mahadevan et al., 2021). Here we tested the hypothesis that low reliability in partial correlation (and related methods) is due to excessive model complexity resulting in overfitting to noise. Consistent with our hypothesis, we found that regularization significantly improves the repeat reliability and accuracy of partial correlation FC. These results suggest regularized partial correlation is a strong candidate for replacing pairwise correlation as the next field standard for estimating FC using fMRI. These results also suggest that FC methods used with high temporal resolution data (such as EEG) could benefit from regularization as well, especially given the additional model complexity that comes with modeling temporal lags (e.g., with multivariate autoregressive modeling; Fiecas et al., 2010; Haufe et al., 2010; Mill et al., 2022).

We found that regularized partial correlation FC improved on both pairwise correlation and unregularized partial correlation along multiple dimensions. First, reliability was substantially improved in regularized (e.g., mean r = 0.64 for graphical lasso empirical between-session similarity) relative to unregularized partial correlation (e.g., mean r = 0.11; Figure 3). This result essentially “rescues” partial correlation as a method, given how essential reliability is to FC estimation (Noble et al., 2021). This was especially beneficial in conditions of low amounts of data and high noise (Figure 7), which can further impair reliability (Birn et al., 2013). Second, the validity of FC estimates was much higher for regularized partial correlation relative to pairwise correlation. This was demonstrated in terms of increased similarity of regularized partial correlation FC to empirical structural connectivity (Figure 4) and simulated ground truth FC (Figure 6). Notably, this likely reflects partial correlation reducing the number of confounded (e.g., a false *A*-*B* connection due to an upstream region *C*: *A* ← *C* → *B*) and indirect (e.g., a false *A*-*B* connection due to an intermediate region *C*: *A* → *C* → *B*) connections (Figure 1A), which structural connectivity and ground truth FC are insensitive to. Further supporting validity, regularized partial correlation FC was found to be less sensitive to in-scanner motion than pairwise correlation FC, demonstrating reduced confounding from non-neural factors as well. Finally, given the importance of reliability and validity for a variety of FC applications, we tested for improvements to FC-based applications in cognitive neuroscience. As expected, we found that regularized partial correlation significantly improved FC-based generation of task-evoked fMRI activations (Figure 8) and prediction of individual differences in age and behavior (Figure 9). Together, these results demonstrate the improved reliability, validity, and general utility of regularized partial correlation – especially graphical lasso – relative to pairwise correlation and unregularized partial correlation.

We conceptualize FC as the functional manifestation of grounded physical processes, such that every functional connection must have a corresponding structural connection *at a minimum*. This is the minimum because FC provides more information regarding the interaction of neural populations than structural connectivity. This reflects the impact of various functional factors on time series similarity. For instance, in our simulations we found that regularized methods reflected simulated aggregate synaptic weights, in addition to the structural connections necessary to support those synaptic weights. We may also expect effects from neurotransmitter concentrations (e.g., from arousal) to impact FC, along with other functional factors. Thus, while similarity to structural connectivity was important as a minimal case (to show we have reduced false positives from confounding) we did not expect a full match with structural connectivity given these other factors.

The central finding of this study is that regularization stabilizes the connectivity estimates of partial correlation (and multiple regression) FC, increasing the method’s reliability. We demonstrated this in multiple ways. First, we provided a visual depiction of measurement instability in FC matrices, where an individual subject’s partial correlation FC varied discernibly from the group-averaged matrix (Figure 2). In the individual matrix, random noise obscured an underlying network structure, while in the group matrix, that noise had been reduced by averaging. Regularization reduced this noise at the individual subject level. We then quantified repeat reliability in empirical data using between-session similarity and intraclass correlation, where all regularized methods substantially improved on the reliability of partial correlation FC.

Our findings also agree with those of several previous studies. Brier et al. (2015) showed that Ledoit-Wolf shrinkage (Ledoit & Wolf, 2003), a regularization technique not investigated here, improved the reliability of partial covariance FC. Mejia et al. (2018) also demonstrated that L_2_-regularized partial correlation achieved greater reliability as the L_2_ penalty (degree of regularization) was increased. Our results somewhat contrast with those of Fiecas et al. (2013) and Mahadevan et al. (2021), who tested regularized partial correlation, in that their results indicate a more subtle impact of regularization on reliability. We suspect, however, that this apparent small difference in regularized and unregularized partial correlation is due to their measuring reliability using intraclass correlation for all possible edges. We found that null edges achieve poor ICC scores regardless of their reliability (see Methods for elaboration), meaning that the abundance of invariably low scores from these sparse FC methods would have eclipsed any meaningful changes in ICCs for non-null edges. Our analysis using intraclass correlation, which instead examined a conservative set of non-null edges, showed a clear improvement of ICCs with regularization. In utilizing several regularization methods, two reliability metrics, and empirical and simulated datasets, our study provides comprehensive support that regularization can substantially improve the repeat reliability of multivariate FC methods, which are otherwise prone to overfitting to noise.

We also analyzed the validity and individual measurement accuracy of the FC methods, which we define as the systematic correctness of estimates in aggregate (independent of reliability) and the closeness of individual estimates to the truth (dependent on validity and reliability). Previous studies have tested regularized FC methods based on simulations (Nie et al., 2017; Ryali et al., 2012; Smith et al., 2011), generalization to held-out data (Varoquaux et al., 2010), network modularity (Brier et al., 2015; Mahadevan et al., 2021; Ryali et al., 2012; Varoquaux et al., 2010), sensitivity to task state (Brier et al., 2015; Duff et al., 2013; Sala-Llonch et al., 2019), prediction of individual differences (Pervaiz et al., 2020), and similarity to structural connectivity (Liégeois et al., 2020). We chose to primarily validate FC with structural connectivity and simulated networks because these both (attempt to) represent direct connections only – a property which allows FC to better reflect the causal mechanisms and physical reality of brain networks (Reid et al., 2019).

Structural connectivity estimates direct connections by mapping white matter tracts across the brain volume to their grey matter endpoints (Le Bihan & Johansen-Berg, 2012). These anatomical connections are the basis for brain-wide communication and are expected to exist between regions with direct functional connections. Structural connectivity does not measure precisely the same phenomena as FC, as structural connectivity shows predominantly static pathways while FC is also influenced by dynamic factors such as cognitive state (Cole et al., 2014). Structural connectivity estimated from diffusion MRI is also prone to systematic errors, such as from failing to resolve crossing fibers or underestimating long-distance tracts (Maier-Hein et al., 2017; Rheault et al., 2020; Sotiropoulos & Zalesky, 2019). Nevertheless, the two measures largely overlap (Damoiseaux & Greicius, 2009; Honey et al., 2009; Straathof et al., 2019; van den Heuvel et al., 2009). Because structural connectivity is not susceptible to causally confounded and indirect connectivity, it should better converge with the FC whose methods limit these errors (Damoiseaux & Greicius, 2009; Honey et al., 2009; Liégeois et al., 2020). For instance, (Liégeois et al., 2020) have shown partial correlation FC to be more similar to structural connectivity than is pairwise correlation FC. Further, Wodeyar et al. (2020) found that stroke lesions to white matter tracts are more predictive of changes in partial over pairwise correlation FC in the impacted edges.

With simulations, we had access to the true causal connections against which we could compare FC estimates to gauge their validity and accuracy. However, simulations also require the selection of many model parameters, the choices of which can potentially bias performance among FC methods. We opted for a simple linear model to simulate activity over bidirectional networks, which were generated with modular and approximately scale-free structure – properties the human brain has been shown to have (Bullmore & Sporns, 2009; van den Heuvel & Sporns, 2011). As with structural connectivity, prior simulation studies have shown partial correlation FC to emulate true connectivity more closely than pairwise correlation FC (Nie et al., 2017; Smith et al., 2011). We thus validated the FC methods by comparing them against approximate (structural) or certain (simulated) causal networks to further increase confidence in our results.

One aim of these analyses was to ensure that the regularization techniques did not reduce the underlying validity of FC measures, estimated as the correctness of the group-level means. We hypothesized that unregularized partial correlation FC would exhibit high validity, and indeed, after averaging across estimates, it performed substantially better than pairwise correlation FC at reflecting structural and simulated network connectivity. We found group-level graphical lasso FC to perform similarly to partial correlation, scoring slightly worse when tested with simulations but showing no significant difference when tested against SC. As regularization was only hypothesized to aid reliability, this was the expected result for analyses of group-level FC where the effect of reliability had been controlled via cross-subject averaging. Unexpectedly, however, we found graphical ridge and PC regression to have significantly worse validity, a pattern that emerged for both structural connectivity and simulation analyses. This result also aligns with the FC matrix visualizations (Figure 2), where graphical ridge and PC regression FC show visibly different group-level network structures relative to partial correlation and graphical lasso FC. Evidently, the more diffuse connectivity patterns produced by graphical ridge and PC regression are less valid representations of direct FC than the sparser alternatives.

We also considered whether our analyses could have biased the results toward sparser FC methods by gauging FC against targets more sparse than the brain’s “true” direct FC – SC due to methodological shortcomings and simulated networks due to our design and assumptions. We therefore reran the simulations using networks twice as dense as the originals, and while this somewhat improved validity of graphical ridge and PC regression, graphical lasso remained most valid (Supplementary Figure S5). To our knowledge, this is the only published study to examine the effect of these regularization techniques on group-level validity, controlling for the benefit of improved reliability. Our study suggests the novel finding that graphical ridge and PC regression may reduce the group-level validity of FC estimates while graphical lasso can better maintain the validity achieved by unregularized partial correlation.

Our second aim was to determine whether the improved reliability of the regularized FC methods could grant them greater individual measurement accuracy than pairwise correlation and unregularized partial correlation FC. Individual subject measurement accuracy is a central goal of precision neuroscience (Hermosillo et al., 2024) and is ideal for clinical applications. It is a function of both reliability and the underlying validity of the measurement (see Figure 1). We found that the individual estimates of pairwise correlation FC are hindered by its low validity while the scores of partial correlation FC are hindered by its low reliability. The regularized methods generally perform better, led by graphical lasso FC, which most increased reliability and best preserved underlying measurement validity.

The results were largely consistent between the structural connectivity and simulation analyses, and they agree with results from prior studies. (Liégeois et al., 2020) also compared FC estimates with structural connectivity and showed graphical ridge to be more similar than both partial correlation and pairwise correlation FC. (Smith et al., 2011) and (Nie et al., 2017) showed graphical lasso recreates simulated networks more accurately than pairwise correlation FC, although there was not always a large difference between graphical lasso and unregularized partial correlation. We suspect that this was due to the simulations using very few nodes (usually 5 or 10) relative to the amount of data (usually 200 TRs or more), causing less overfitting to noise than should be expected from an empirical dataset. Regularization had a much larger benefit during their simulations with 50 nodes, where partial correlation was impaired by a more realistic degree of overfitting. (Varoquaux et al., 2010) tested the generalizability (via loglikelihood) of FC models to held-out rest data, showing graphical lasso and graphical ridge both perform better than partial correlation when applied to individual subjects’ data. They also found graphical lasso to exhibit higher generalizability than graphical ridge, further supporting our finding that graphical lasso is the more valid regularized FC method.

All brain recording modalities have limitations, with some weaknesses of fMRI being scanner noise, low sampling rate and constrained scan times, and susceptibility to head motion artifacts. We hypothesized that partial correlation would be highly sensitive to noise and number of timepoints, as these can influence the extent of model overfitting (Blum et al., 2020; Hastie et al., 2009; Ying, 2019), and that regularization would ameliorate this. These hypotheses were supported by our simulation results.

While partial correlation achieved nearly perfect ground truth similarity with lower noise levels and a very large number of timepoints (100 times as many TRs as nodes – an unrealistic amount of empirical fMRI data), there was a steep decline in performance with decreasing timepoints and increasing noise. The decline is especially sharp approaching an equal number of timepoints as nodes, for which the ground truth correlation is almost zero, and partial correlation is unable even to calculate FC where there are fewer timepoints than nodes. Our results demonstrate that regularization can handle such scenarios where there are fewer timepoints than nodes – a common use for regularization as it helps constrain fitting procedures that are ill-posed.

Regularization especially improves accuracy in the lowest timepoint conditions, with graphical lasso offering the largest benefits, across the full range of conditions. Previous studies have shown similar patterns in empirical data, with Liégeois et al. (2020) demonstrating that L2-regularized precision is more similar to SC than is unregularized precision across a range of scan lengths, but especially with the shortest scan durations. Meanwhile, Mejia et al. (2018) found that shorter scans prefer a greater amount of shrinkage than longer scans. While regularized multivariate FC estimates still benefit from increased data and decreased noise, these results indicate that they can also be used with legacy datasets that have shorter rest durations or longer TRs, as well as with data from subject populations that cannot tolerate long MRI scans.

While not considered one of our primary tests of validity, sensitivity to motion artifacts renders FC less valid because it is then capturing non-neural effects. Unlike random noise in coefficient weights, the FC variations introduced by subject head motion are systematic, for instance leading to decreases in long-distance connections and increases in short-range connections (Power et al., 2012). We found that pairwise correlation FC was highly susceptible, with half of all edges showing significant correlation with subject motion, despite also employing intensive motion artifact mitigation strategies (see Methods). In contrast, partial correlation FC had no significant edges. Largely maintaining this result, regularization added only a fraction of a percent of significant edges. This replicates the findings of Mahadevan et al. (2021), who demonstrated high motion sensitivity of pairwise correlation FC but low sensitivity of unregularized and L2-regularized partial correlation FC. While spurious connectivity is undesirable in any study, the effects of motion artifacts on FC can be especially harmful where the variable of interest is associated with head motion. For example, children, older adults, and various clinical populations are prone to more in-scanner movement than healthy young adults, and group differences in FC due to motion could be mistaken for true differences in brain communication. Regularized multivariate FC methods largely eliminate confounding by head motion.

Our final analyses predicting task-evoked activations and individual differences in subject age and intelligence further support our findings of reliability, validity, and accuracy of individual measurements. Moreover, they demonstrate how direct, more causally-valid FC measures can contribute to a mechanistic understanding of cognition and brain function. Activity flow modeling simulates the propagation of task-evoked activations over FC pathways (Cole, 2024; Cole et al., 2016). Activity predictions for each node are generated as the summed activities of all other nodes, modulated by the strengths of their connections to the target node. More accurate FC estimates will generate predictions more similar to the actual activations. When introducing activity flow modeling, Cole et al. (2016) showed that multiple regression FC yielded far more accurate predictions than pairwise correlation FC, suggesting that multiple regression FC better reflects the brain-wide interactions that underlie cognition. Our activity flow analyses also showed the related method of partial correlation FC performing better than pairwise correlation FC. Some positive and negative noisy coefficients of partial correlation FC may have cancelled each other out, but the noise reduction of graphical lasso still offered ample improvement in activity predictions. Graphical ridge and PC regression achieved only slightly worse activity predictions than graphical lasso despite having different network structures (Figure S8), suggesting that their more distributed connectivities can produce similar summations. However, graphical lasso is likely the best choice for activity flow modeling, given its higher validity based on our structural connectivity and simulation analyses. Activity flow over such estimates of direct FC offers a mechanistic explanation of how the connectome shapes the brain activity patterns underlying cognitive functions.

A common goal of neuroscience research is to relate measures of brain function to cognition. Here, we demonstrated the ability of regularized multivariate FC to predict individual differences in subject age and intelligence in models that are both accurate and interpretable. Pairwise correlation and graphical lasso (and the other regularized methods) produced high accuracies that were not significantly different from each other but were all better than unregularized partial correlation. This suggests that model performance depended mainly on reliability and was less affected by redundancy in FC weights. This aligns well with the findings of prior studies. Pervais et al. (2020) found L_1_-regularized partial correlation with group information and L_2_-regularized partial correlation to both predict subject individual differences (e.g., age, intelligence, sex) better than pairwise correlation FC. However, Duff et al. (2013) and Sala-Llonch et al. (2019) tested how well FC applied to resting state or task data could classify different task states, and while L_1_-regularized (Duff et al., 2013) or L_2_-regularized (Sala-Llonch et al., 2019) partial correlation produced overall better performance, pairwise correlation FC achieved better classification for some parcellation schemes. If the most important factor of an analysis is prediction or classification accuracy, for instance for a clinical diagnosis, pairwise correlation FC may be as good a choice as regularized multivariate FC. However, if the analysis aims to elucidate the neural mechanisms and specific pathways that underlie differences in cognition, then regularized multivariate FC offers the benefit of more interpretable (e.g., valid and less numerous) model coefficients.

### 4.1. Limitations and directions for future research

The present study has several limitations, each of which suggest important directions for future research. For example, the present results are based only on empirical and simulated fMRI data, leaving future studies to test the utility of such regularized multivariate methods for FC estimation with different types of brain recordings (e.g., MEG/EEG). Indeed, supporting the possibility that our hypotheses generalize to EEG data, Mill et al. (2022) found evidence that PC regression improves the stability of multivariate autoregressive models. Additionally, (Haufe et al., 2010) showed that a group lasso penalty can also stabilize multivariate autoregressive models, and (Fiecas et al., 2010) found that shrinkage can stabilize partial coherence estimates from EEG data. While these studies show the efficacy of regularized multivariate FC in general for EEG, the field may further benefit from tests comparing different regularized FC methods to establish best practices. For instance, the current study found L_1_ regularization (graphical lasso) most advantageous for fMRI data, but other modalities may benefit more from different methods.

The present study also focused solely on FC at the scale of brain regions, with future research needed to establish the best approaches for regularized multivariate FC at the level of fMRI voxels or other small spatial scales. With voxelwise FC, regularization is not just beneficial but essential when fitting partial correlation or other multivariate FC models, as the number of variables (voxels) typically far exceeds the number the timepoints. Cole et al. (2016) demonstrated the feasibility of PC regression for estimating voxelwise FC and the efficacy of the FC estimates in predicting held-out task activations via activity flow modeling. Some FC methods may have to be applied differently to voxelwise data from how they would be to regionwise data, however. PC regression is more computationally suitable for voxelwise FC because it involves fitting a reduced number of variables, whereas fitting a graphical lasso model with tens of thousands of variables is computationally impractical. Further, we expect that fitting a model with so few timepoints compared to variables would yield low reliability and/or validity. Our simulation analyses may show suitably accurate FC estimates with as few as half the number timepoints as variables (i.e., graphical lasso with low noise; Figure 7), but accuracy declines sharply with decreasing timepoints to variables, and voxelwise FC analyses will typically have a far smaller ratio than even this. One possible solution to aid the estimation of voxelwise FC is to first perform graphical lasso on regionwise data and then limit voxelwise FC calculations to between only those voxels with regional connections (Chakravarthula et al., 2025). Such a strategy imposes a regionwise sparse structure to reduce the number of variables in the voxelwise model. Voxelwise FC can then be calculated with PC regression, for instance, where each target voxel’s FC is fit in an independent model that includes only the voxels of connected regions. Future research is needed to further develop and test regularized multivariate methods for voxel level FC.

As stated previously, we do not know the brain’s true FC, and the targets against which we compared FC estimates (SC and simulation networks) are imperfect models. Even within these strategies, there is a wide range of options for their implementation and our choices could have affected our results. For instance, we only employed SC matrices generated with deterministic tractography, but probabilistic tractography algorithms generally produce denser connectomes. Our SC edges were weighted by normalized streamline count, but a variety of other weighting schemes are available which yield different weight distributions (Nelson et al., 2023). There was an especially wide parameter space when designing our simulations. We attempted to ameliorate this by testing the original simulations with different numbers of timepoints and noise levels, by altering the networks to have twice the original density, and by convolving simulated activity with an HRF. However, we could not test every possible scenario. Future studies will have the opportunity to conduct more comprehensive simulations, such as with different underlying network architectures, more biological detail, and multiple scales simulated (e.g., voxels and brain regions). For example, it is possible that larger modules (regions or networks) produce larger shared signals, which may impact partial correlations and coefficient stability. It will also be important to test network recovery from data simulated from nonlinear processes, as the FC methods studied here measure linear relationships. Such studies could reveal more general conclusions about the utility of different forms of regularization as a function of the specific data analysis scenario.

Perhaps because we did not set out to promote any single regularization approach, we tested more forms of regularization than most prior studies investigating regularization for FC estimation. However, many other forms of regularization have been proposed that we did not test (Mejia et al., 2018; Nie et al., 2017; Ryali et al., 2012; Varoquaux et al., 2010). One promising form of regularization to test in future work will be elastic net (Zou & Hastie, 2005), which combines both L_1_ and L_2_ regularization with potential benefits of both (Ryali et al., 2012). For instance, elastic net is purported to induce sparsity while allowing more sharing of weights between similar variables (Zou & Hastie, 2005). Another adaptation of interest to us is group regularization, where data is pooled across subjects so that the larger quantity of data can improve model fit. Typically, these will estimate the underlying, group-level network structure and the individual variations of each subject from that baseline (Mejia et al., 2018; Varoquaux et al., 2010). Some methods also offer the advantage of not requiring hyperparameter selection, making them easier for researchers to apply (Brier et al., 2015; Ledoit & Wolf, 2003; Nie et al., 2017). For regularization methods that do require hyperparameters, there are various additional model selection methods not tested here that may improve FC accuracy or computational efficiency. We chose to use cross-validated R^2^, but other options include cross-validated loglikelihood (Friedman et al., 2008), extended Bayesian information criterion (Foygel & Drton, 2010), and *Dens* criterion based on sparsity (Wang et al., 2016). Many other forms of FC remain to be tested with regularization as well, given that regularization is applicable to any form of data fitting (e.g., artificial neural network learning) and not only partial correlation and multiple regression. It could also benefit directed FC methods that are prone to overfitting, such as time-lagged multivariate FC with EEG or other high-temporal-resolution data (Antonacci et al., 2024; Mill et al., 2022). Another promising direction for future work is to utilize regularization to counter the excessive flexibility of nonlinear FC methods, reducing the tendency for such methods to overfit to noise.

The methods we found were most effective here involve an additional step – regularization parameter selection – relative to the standard pairwise correlation FC approach. We used cross-validation for parameter selection, with FC estimates used to optimally predict held-out “test” portions of the time series in each cross-validation fold. Cross-validation is commonly used in a variety of machine learning applications, and regularized partial correlation and regression can be considered machine learning approaches. Cross-validation is known to be effective for parameter selection in a variety of circumstances, with our use of external validations here (e.g., simulations) further confirming its effectiveness. Note, however, that cross-validation can add a large computational cost relative to alternatives. In the present study we used 360 time series (one per brain region), and we were able to estimate 30 subjects’ graphical lasso FC (with cross-validation) in 3.75 hours on a 2020 MacBook Pro laptop. Larger node sets (e.g., voxels) would undoubtedly increase the computational cost of cross-validation.

Thus, it will be important for future research to investigate alternate strategies for regularization parameter selection, perhaps based on strategies used in other machine learning contexts.

While we found across multiple validation measures that identifying a more optimal regularization parameter (based on cross-validation) resulted in better results, a range of hyperparameter values produced satisfactory performance, and even a too-small level of regularization often improved results relative to no regularization. Thus, some pressure is taken off of the choice of regularization parameter. For instance, choosing a regularization parameter based on a prior related study could be valid, with improvement of results despite retaining some remaining overfitting to noise. While using cross-validation to identify the regularization parameter is our recommended best practice, using this (or other) alternative approaches to identifying a regularization parameter is unlikely to invalidate the results presented here.

The complexity of regularized partial correlation FC contrasts with many neuroscientists’ preference for reduced FC measure complexity – to minimize assumptions, minimize compute time, and fully comprehend and easily communicate methodological details. However, we showed that the lower complexity of the field-standard Pearson correlation FC method comes with lower validity of the estimated connections. Further, while more compute time is needed for regularized partial correlation (for single instances and for repetitions over different regularization hyperparameters), recent increases in the availability of multi-core processors makes the increased compute time much less severe than just a few years ago. This also opens up an opportunity for future research to reduce processing time by identifying standard regularization parameters for common situations, such as when a particular fMRI sequence, scan duration, and region set are used. It is also possible (given the low variability of hyperparameters across subjects in our study) that compute time could be reduced by using the optimal hyperparameters from a small subset of subjects as the basis for hyperparameters for all remaining subjects in a dataset.

This study demonstrated the resilience of multivariate FC methods to head motion artifacts – achieved by partial correlation and maintained by the regularized methods – but this benefit may apply to other common confounds as well, such as global signal. While our focus here has been on removing the influence of actual neural signal from confounding or intermediary nodes, the influence of non-neural confounds could also be controlled for if they are sufficiently sampled by the other nodes. For instance, head motion can cause correlated/anticorrelated signal fluctuations across the whole brain (Power et al., 2018), and if it is sufficiently represented across a variety of nodes, these artifactual signal fluctuations can be regressed from the targets. This may also apply to fMRI global signal artifacts (Power, Plitt, et al., 2017), which have been notoriously difficult to remove without including the signals of interest within the to-be-regressed-out global signal estimate. This may inappropriately distorting the signal of interest (Power, Laumann, et al., 2017; Saad et al., 2012). As global signal should be widely present in the non-target nodes, it too should be regressed out of target node timeseries along with the residual neural activities. Future studies will be needed to confirm this, but if true, it could obviate the need for global signal regression, accomplishing its goals while avoiding its pitfalls, with no additional preprocessing.

Given that direct FC can be thought of as structural connectivity with functional modifications (e.g., based on the aggregate effects of synaptic strengths), one may wonder if direct FC should replace estimates of structural connectivity. In some cases that may be preferable, but likely not in most cases. This reflects the imperfection of every connectivity method – while direct FC has unique strengths (e.g., accuracy is not distance dependent) so too does diffusion MRI-based structural connectivity (e.g., no time series-based confounding problem). Thus, both direct FC and structural connectivity can continue to be used to refine inferences regarding brain connectivity. Further, given that direct FC reflects both structural connectivity and additional functional factors, any given direct FC result can be supplemented by a structural connectivity result to give clues as to the role of structural connectivity (versus other factors) in generating that direct FC result. For example, a given direct FC estimate may be especially high for a group of subjects, and structural connectivity estimates may reveal that this is due to additional white matter tracts or higher myelination of those tracts (as opposed to other functional factors). Thus, not only does structural connectivity remain relevant when using direct FC, it may be more useful in combination with direct FC than standard pairwise FC, given the theoretical correspondence between structural connectivity and direct FC (i.e., that every direct functional connection should have a corresponding structural connection) and the inferences their comparison makes possible.

Our primary motivation for using partial correlation FC (rather than the more common pairwise correlation FC) has been to reduce the number of false connections due to confounders (Reid et al., 2019). However, it is possible that hidden/unobserved confounders result in false connections even with regularized partial correlation. This possibility is substantially reduced (relative to, e.g., multi-unit recording) with wide-field-of-view imaging methods like fMRI or MEG/EEG. However, one straightforward way for future studies to reduce the chance of false positives from confounders is to include subcortical regions in addition to the cortical regions used here, given that there is substantial interaction between cortical and subcortical regions (Ji et al., 2019). It may also be advantageous to use regularized partial correlation at the highest spatial resolution available (e.g., voxels) rather than brain regions, given the possibility that confounder time series may not be fully accounted for when averaged into a larger neural population’s time series.

### 4.2. Conclusion

The brain is a complex network, producing cognition through distributed processing by many regions. To understand the brain, it is therefore crucial to understand how its regions communicate, as is attempted with FC analyses. However, the quality of inferences depends on the performance of the specific FC method. Here, we explore the issue of instability in multivariate FC methods, which are otherwise advantageous, and demonstrate their enhancement with regularization techniques. We thoroughly examine the performance of these methods, utilizing held-out resting-state fMRI data, structural connectivity, fMRI task activations, behavior, and simulations to assess both validity and reliability. In all, we show the regularized methods (especially graphical lasso) to be robust estimators of functional connections, which have strong potential to improve the quality of future FC studies.

#### Ethics

This study used data from the Human Connectome Project in Aging dataset, released through the NIMH Data Archive (NDA). All subjects gave signed informed consent in accordance with the protocol approved by the institutional review board associated with each data collection site (Washington University St. Louis, University of Minnesota, Massachusetts General Hospital, and University of California, Los Angeles). We followed the terms set in the NDA Data Use Certification, and our use of this data was approved by the Rutgers University institutional review board.

#### Author contributions

**Kirsten L. Peterson:** Conceptualization, Methodology, Software, Validation, Formal analysis, Writing – Original Draft, Writing – Review & Editing, Visualization

**Ruben Sanchez-Romero:** Methodology, Software, Writing – Review & Editing

**Ravi D. Mill:** Methodology, Writing – Review & Editing

**Michael W. Cole:** Conceptualization, Methodology, Writing – Original Draft, Writing – Review & Editing, Supervision, Funding acquisition

#### Data and code availability

Data and code to reproduce these analyses will be made available upon publication of this manuscript.

## Acknowledgements

This work was supported by the US National Science Foundation (NSF) under award 2219323. R.S.-R. was also supported by the US National Institute of Neurological Disorders and Stroke (NINDS) and National Institute on Drug Abuse (NIDA) under grant R01NS120289. The empirical data used here (Human Connectome Project in Aging) was supported by the National Institute On Aging of the National Institutes of Health under Award Number U01AG052564 and by funds provided by the McDonnell Center for Systems Neuroscience at Washington University in St. Louis. We thank the Office of Advanced Research Computing at Rutgers, The State University of New Jersey, for providing access to the Amarel cluster and associated research computing resources that have contributed to the results reported here. This content is solely the responsibility of the authors and does not necessarily represent the official views of any of the funding agencies.

## Supplementary Materials

**Figure S1.**
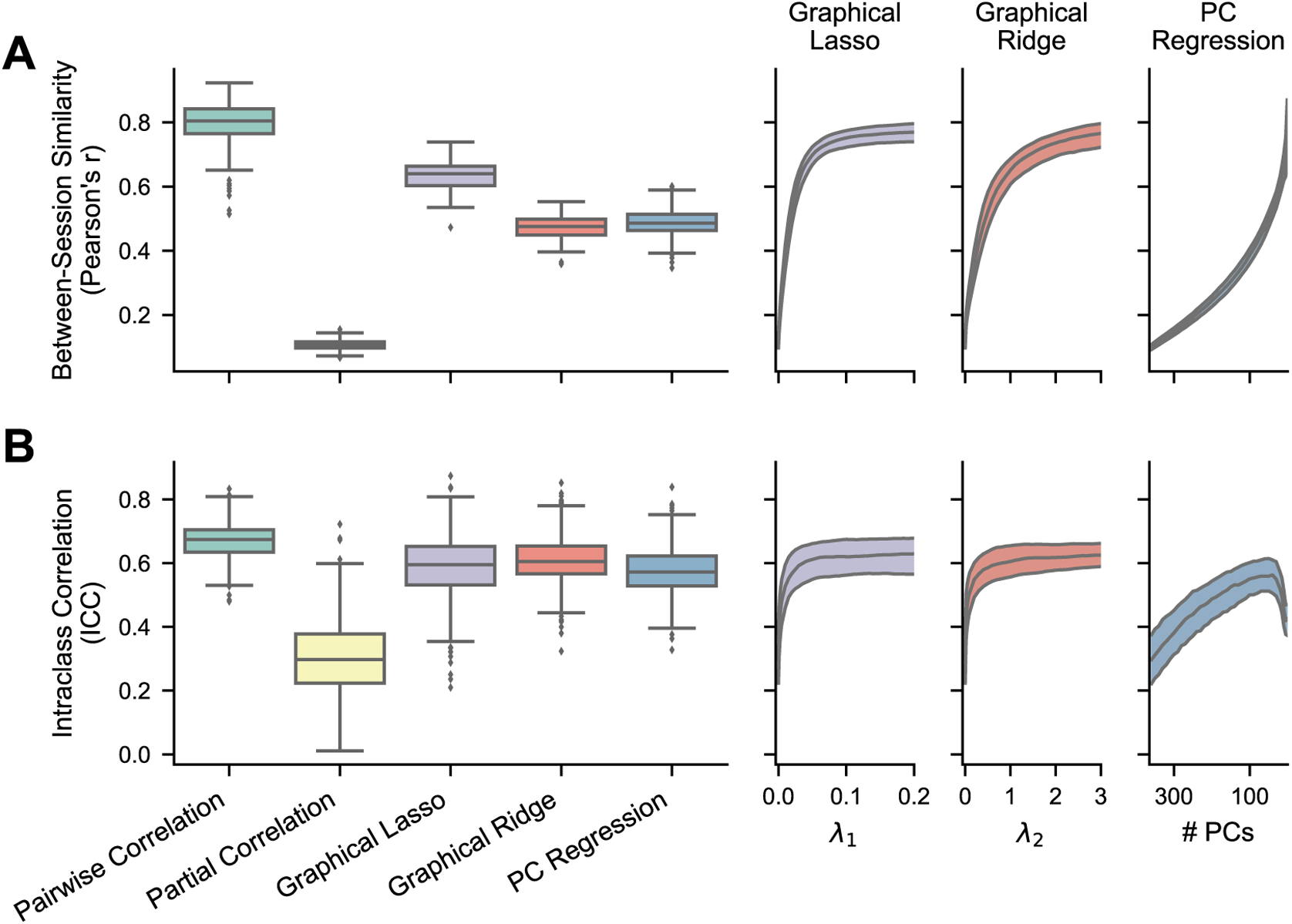
– Reliability of FC methods with empirical fMRI data – replication dataset. The boxplots show results where the regularization hyperparameters have been optimized for each FC matrix, while the right plots show the medians and IQRs across different hyperparameter values for the regularized methods. For PC regression, number of PCs is plotted in descending order because fewer PCs correspond with more regularization. **A)** Between-session similarity, the Pearson correlation between each subject’s session 1 and session 2 FC matrices (n = 236). **B)** Intraclass correlation (Shrout & Fleiss, 1979), calculated for each edge in a conservative subset of nonzero edges (n = 532).

**Table S1.**
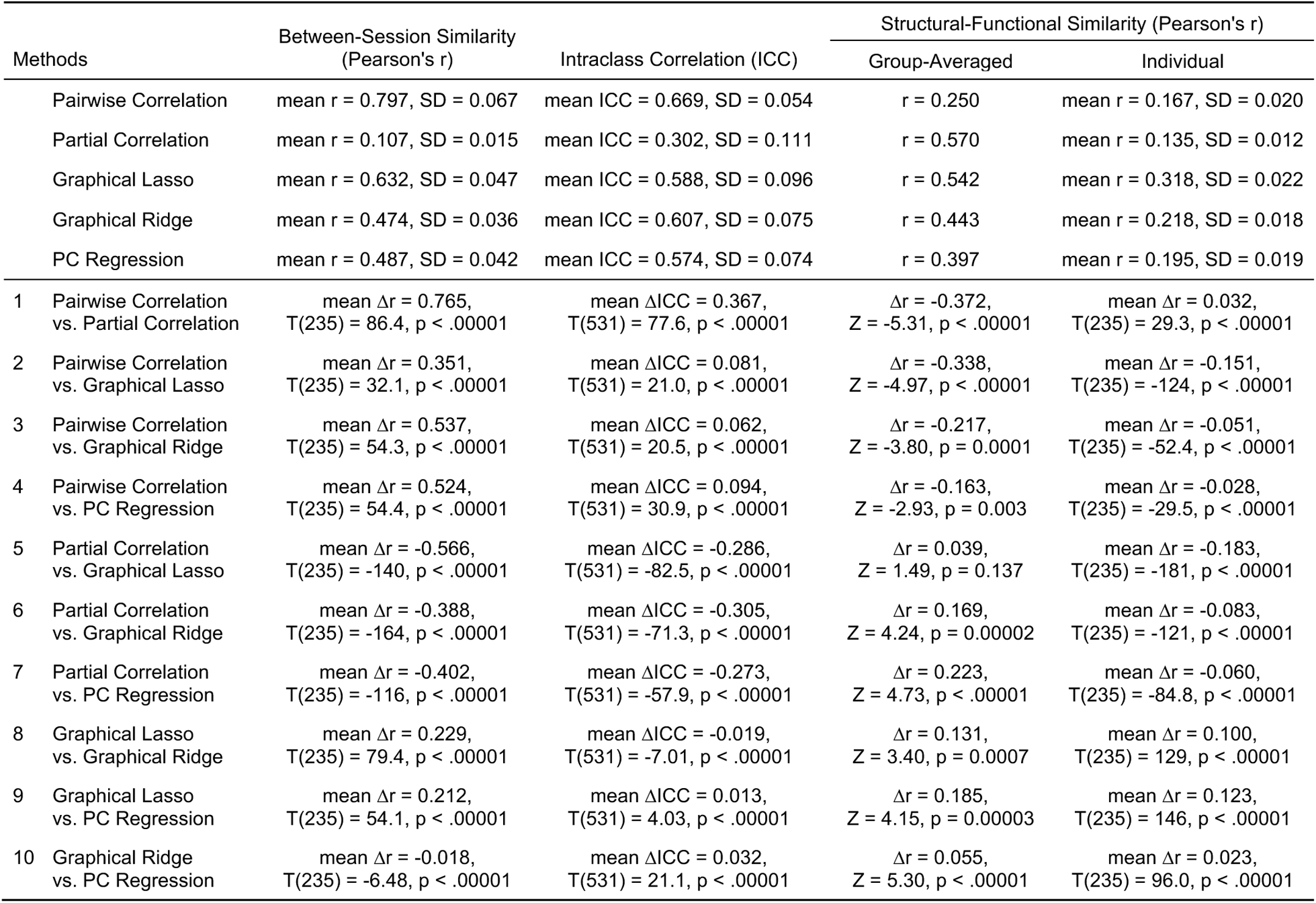
– Statistical tests comparing FC methods on reliability and structural-functional similarity using empirical data – replication dataset. Between-session similarity, intraclass correlation, and individual structural-functional similarity were compared using two-tailed, dependent-sample t-tests. The Pearson’s r values for between-session similarity and individual structural-functional similarity were normalized using Fisher’s z transformation (arctanh) before statistical comparisons. Group-averaged structural-functional similarity (single Pearson’s r per method) was compared using the method described by Meng et al. (1992), which tests for a significant difference between “correlated correlations” (correlations with a shared variable – SC weights in this case). For all reported correlation differences, correlations were transformed to Fisher’s z, subtracted, and then transformed back to Pearson’s r. Alpha levels were adjusted to .005 (.05/10) following Bonferroni correction to account for the multiple comparisons within each column.

**Figure S2.**
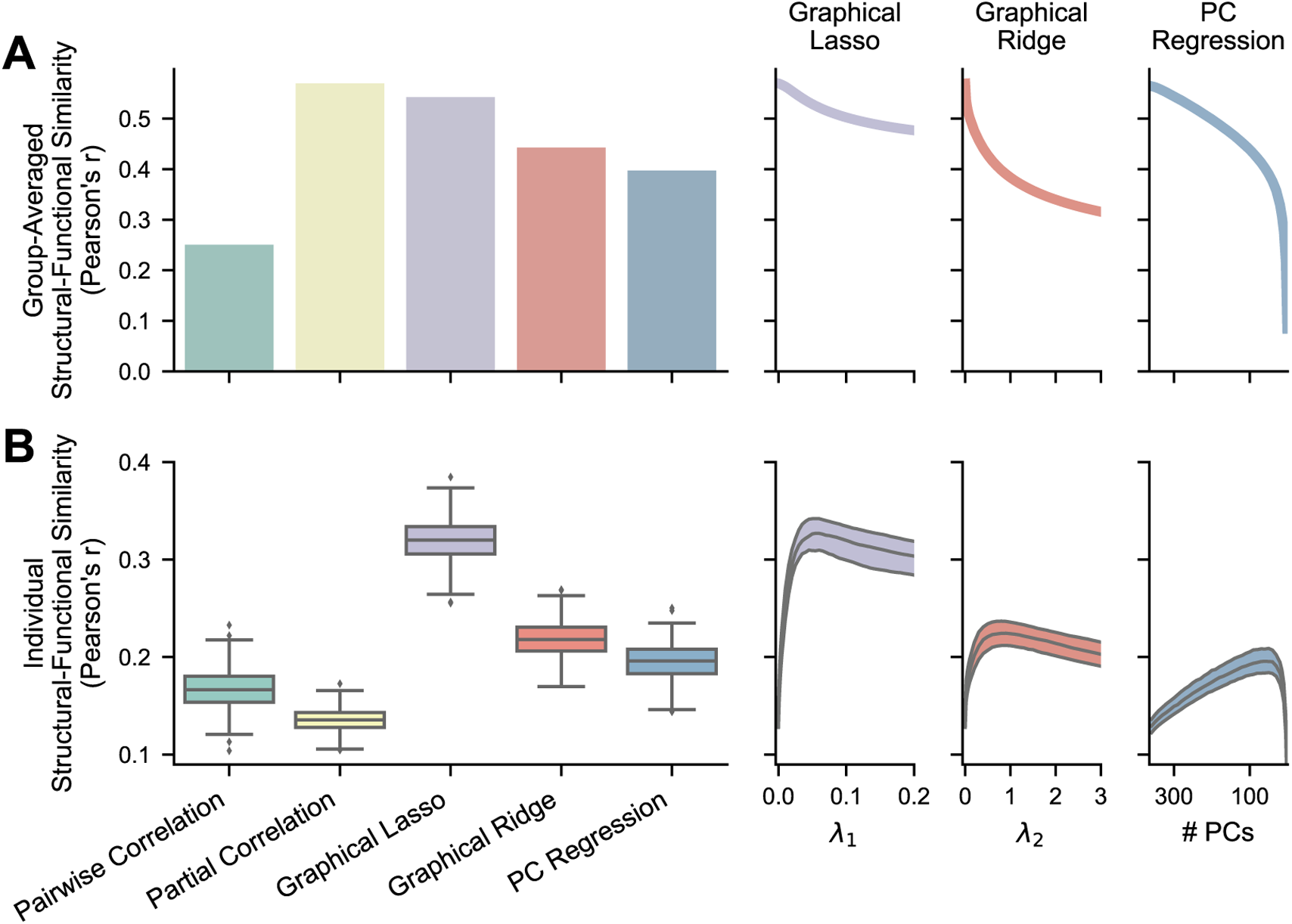
– Similarity between empirical FC and SC (diffusion MRI tractography) – replication dataset. The bar and boxplots show results where the regularization hyperparameters have been optimized for each FC matrix, while the right plots show the single values or the medians and IQRs across different hyperparameter values for the regularized methods. For PC regression, number of PCs is plotted in descending order because fewer PCs correspond with more regularization. **A)** Structural-functional similarity between group-averaged FC and structural matrices (n = 1). Averaging nullifies much of the noise in individual connectivity weights to show validity without the effects of low reliability. **B)** Structural-functional similarity between individual FC matrices and the same subjects’ structural matrices (n = 236). Individual measurement accuracy is vastly improved by recovering reliability.

**Figure S3.**
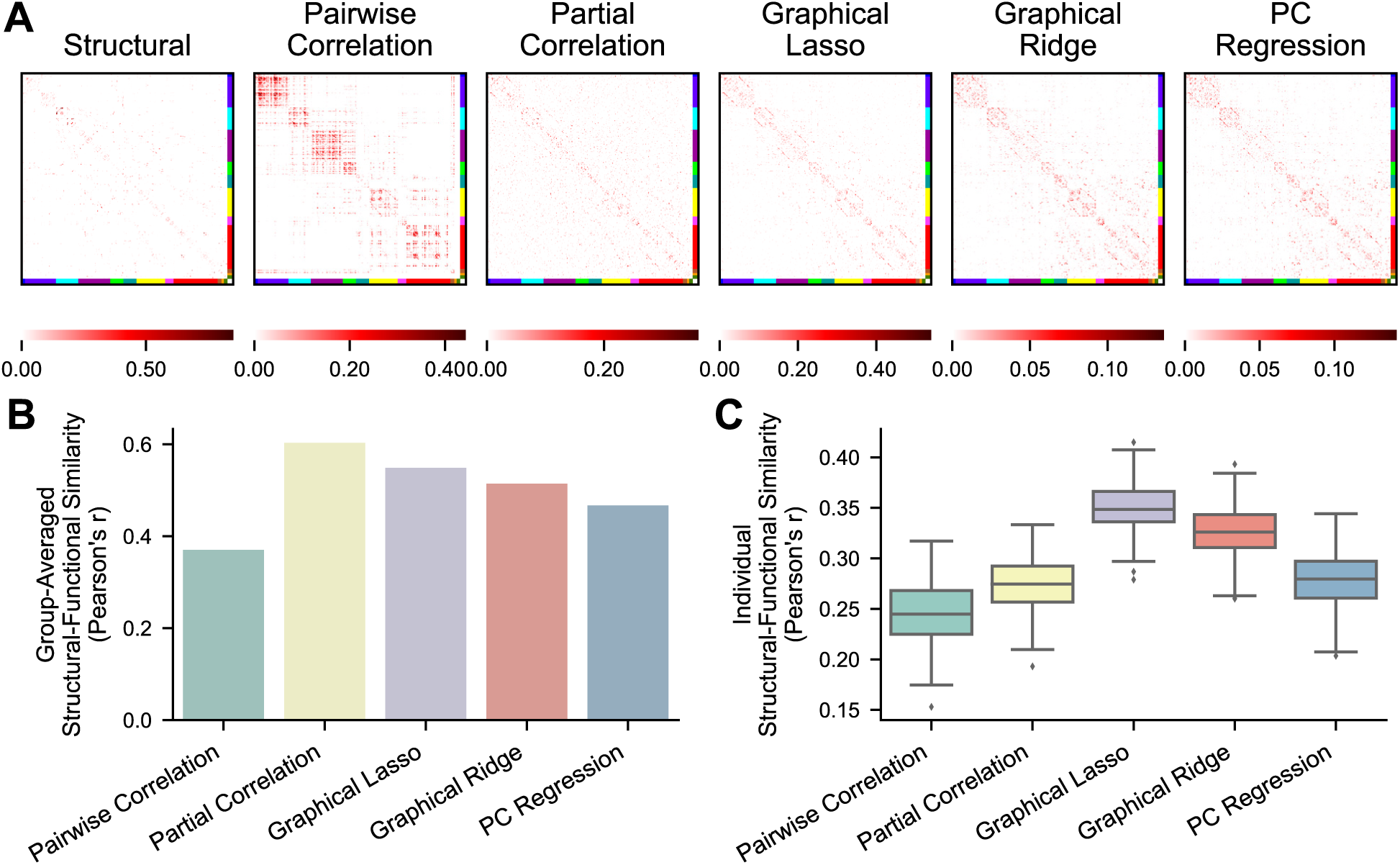
– Similarity between empirical FC and SC (diffusion MRI tractography) – sparsity-matched FC. **A)** Single subject’s SC matrix, and FC matrices thresholded to have the same sparsity as SC. **B)** Structural-functional similarity between group-averaged sparsity-matched FC and structural matrices (n = 1). Averaging nullifies much of the noise in individual connectivity weights to show validity without the effects of low reliability. **C)** Structural-functional similarity between individual sparsity-matched FC matrices and the same subjects’ structural matrices (n = 236). Individual measurement accuracy is vastly improved by recovering reliability.

**Table S2.**
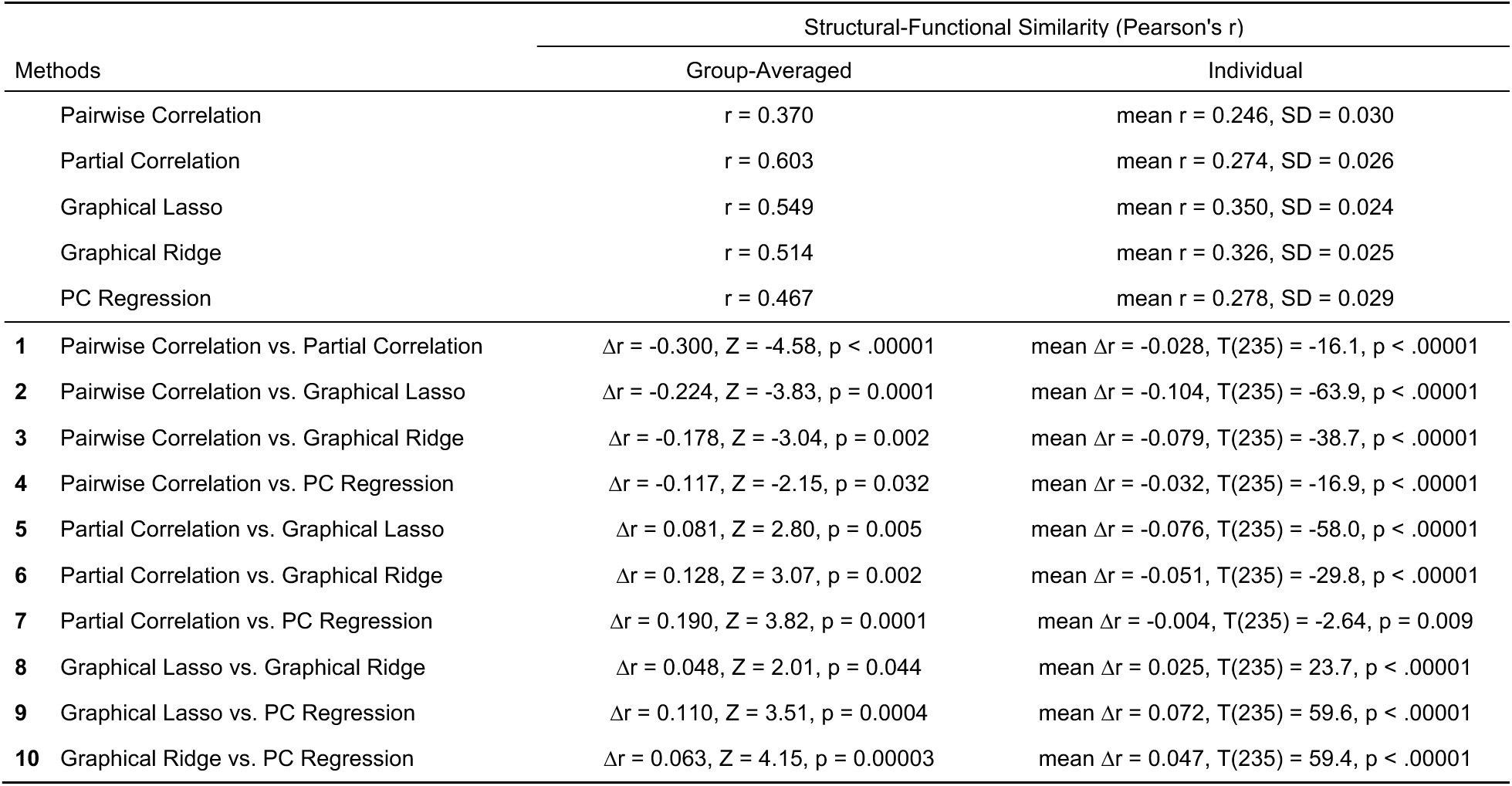
– Statistical tests comparing FC methods on structural-functional similarity with sparsity-matched FC. Individual structural-functional similarity was compared using two-tailed, dependent-sample t-tests. The Pearson’s r values for individual structural-functional similarity were normalized using Fisher’s z transformation (arctanh) before statistical comparisons. Group-averaged structural-functional similarity (single Pearson’s r per method) was compared using the method described by Meng et al. (1992), which tests for a significant difference between “correlated correlations” (correlations with a shared variable – SC weights in this case). For all reported correlation differences, correlations were transformed to Fisher’s z, subtracted, and then transformed back to Pearson’s r. Alpha levels were adjusted to .005 (.05/10) following Bonferroni correction to account for the multiple comparisons within each column.

**Figure S4.**
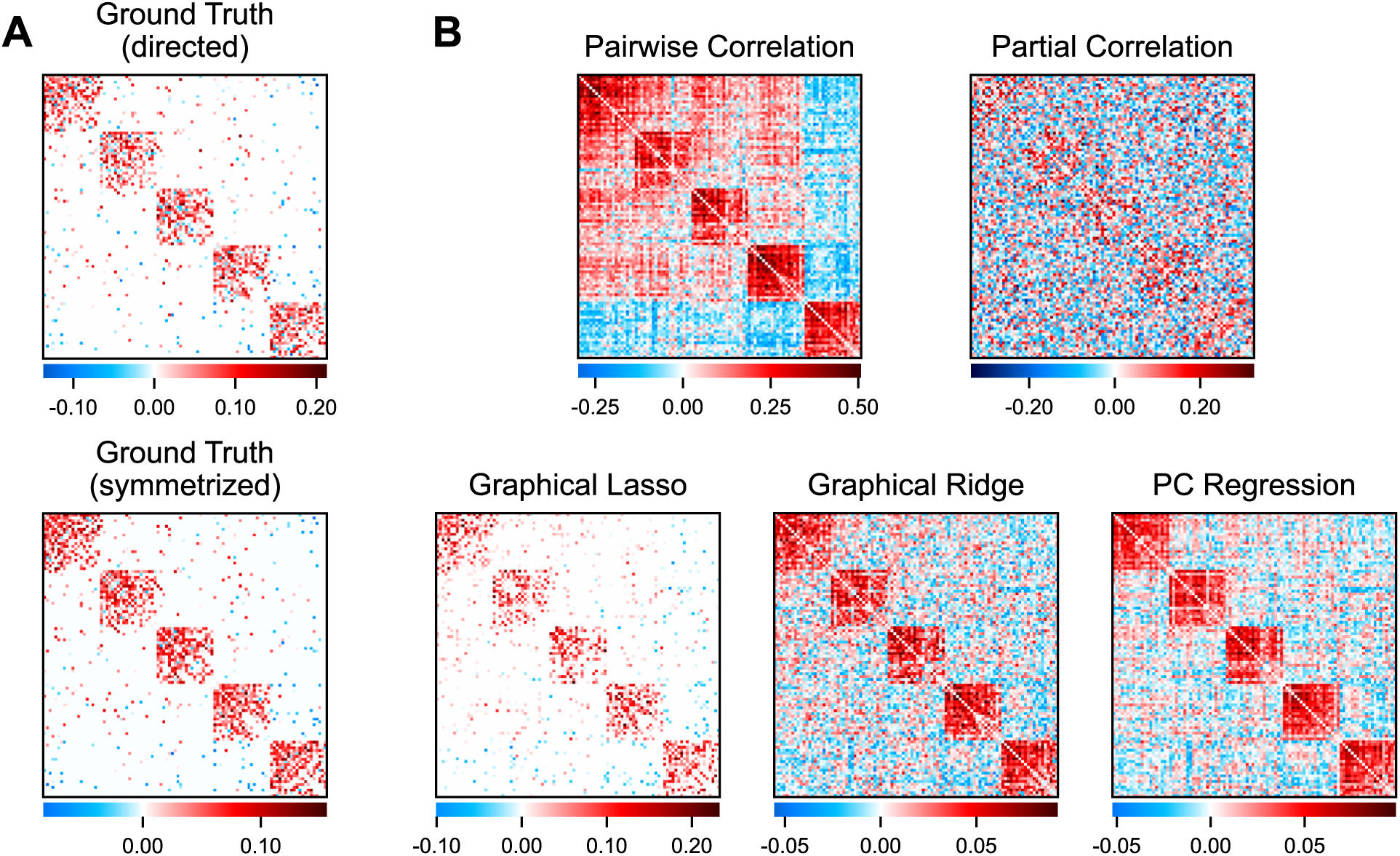
– Simulated networks and FC estimated by different methods – less sparse networks. **A)** An example ground truth network. The directed, asymmetric version (top) was used to generate nodes’ timeseries while the symmetrized version (bottom) was compared with FC estimates. **B)** Individual FC matrices estimated from a simulated timeseries using each FC method.

**Figure S5.**
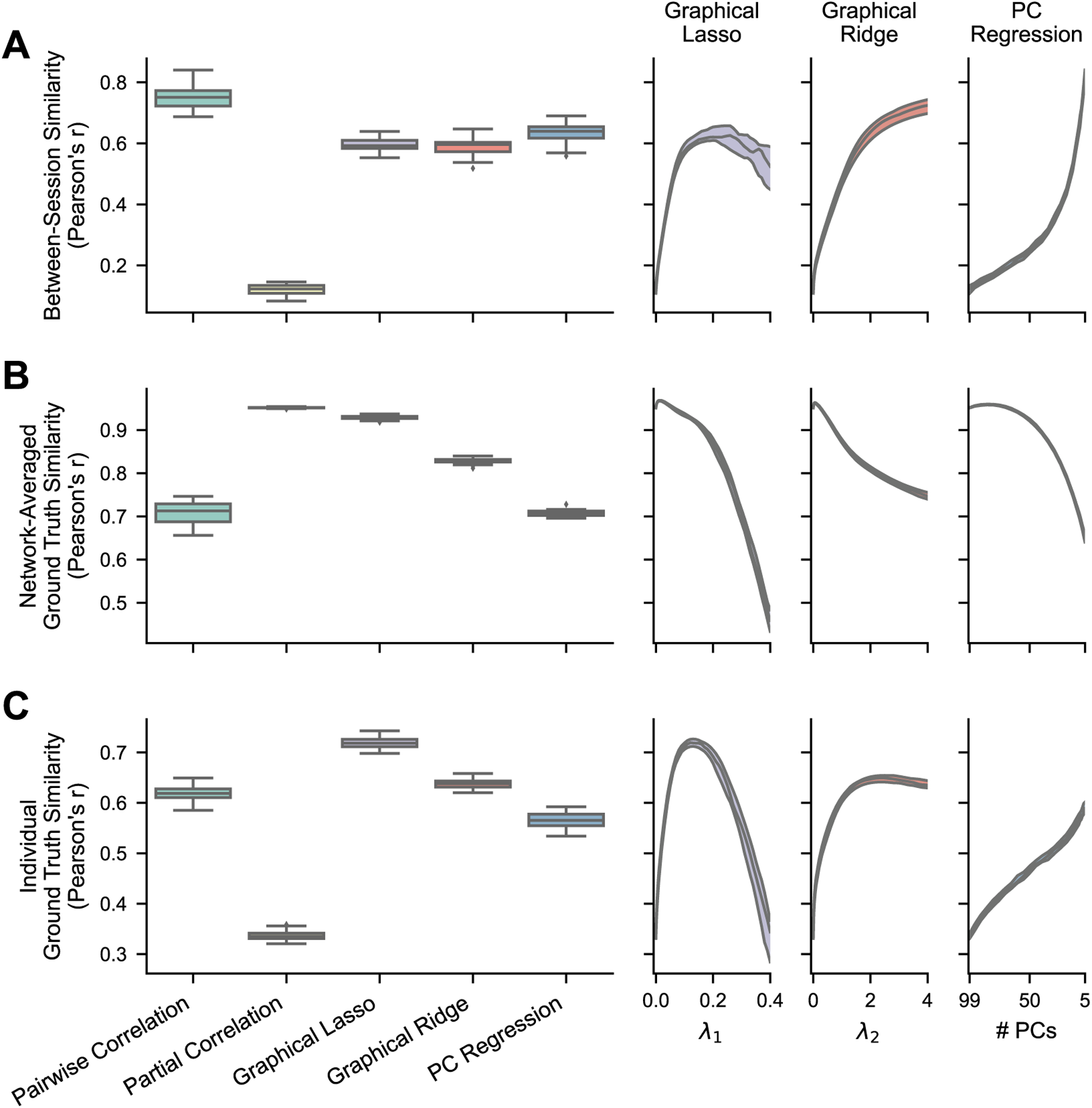
– Reliability and ground truth similarity of FC methods with simulated data – less sparse networks. The boxplots show results where the regularization hyperparameters have been optimized for each FC matrix, while the right plots show the medians and IQRs across different hyperparameter values for the regularized methods. For PC regression, number of PCs is plotted in descending order because fewer PCs correspond with more regularization. **A)** Between-session similarity, calculated between one pair of session matrices for each simulated network (n = 25). **B)** Ground truth similarity between group-averaged FC matrices (100 sessions each) and the ground truth for each simulated network (n = 25). Averaging nullifies much of the noise in individual connectivity weights to show validity without the effects of low reliability. **C)** Ground truth similarity between an individual session’s estimated FC matrix and the ground truth for each simulated network (n = 25). The accuracy of single measurements is vastly improved by recovering reliability through regularization.

**Table S3.**
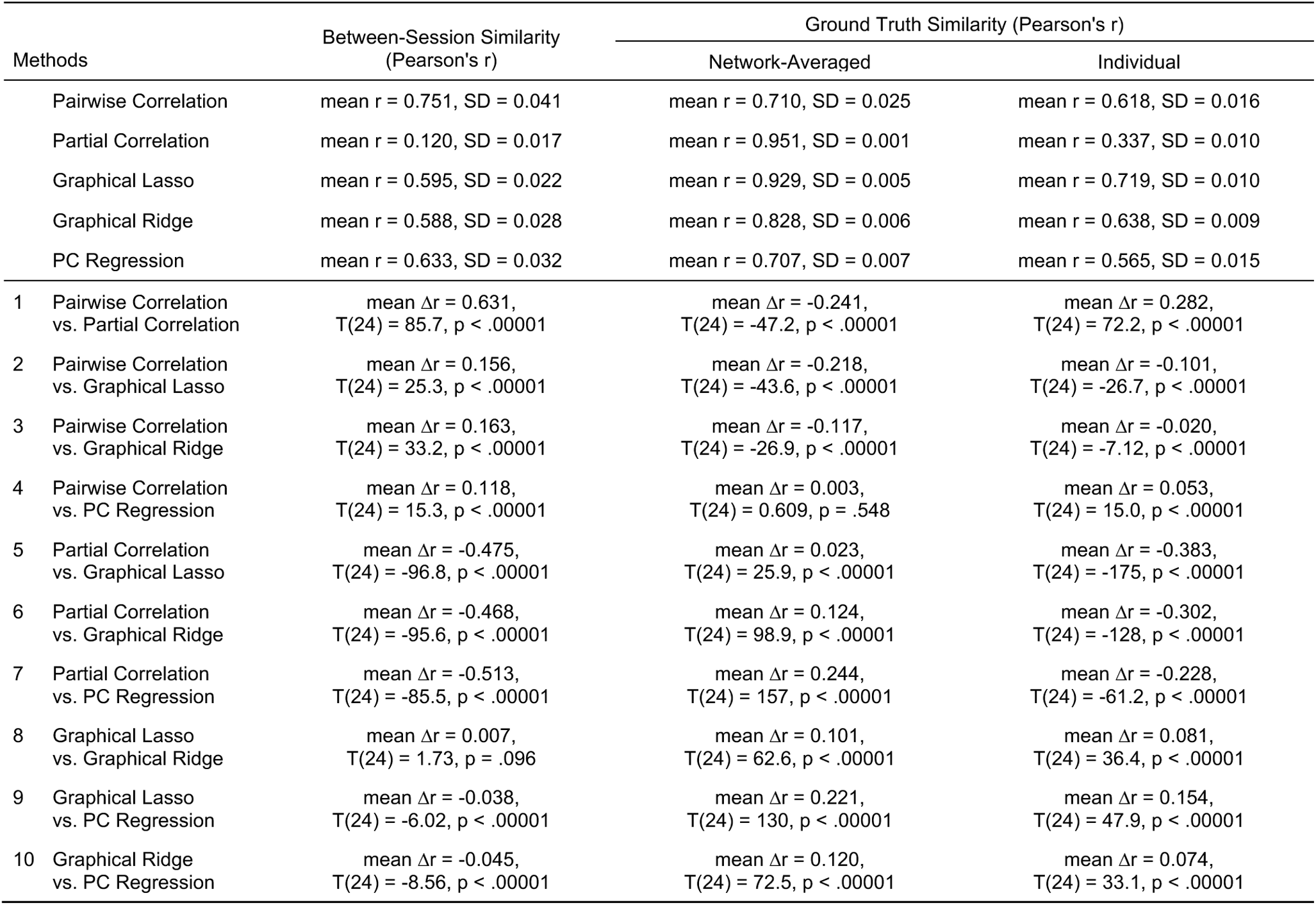
– Statistical tests comparing FC methods on reliability and ground truth similarity using simulated data – less sparse networks. Between-session similarity, group-averaged ground truth similarity, and individual ground truth similarity were compared using two-tailed, dependent sample t-tests, their Pearson’s r values first normalized using Fisher’s z transformation (arctanh). for all reported correlation differences, correlations were transformed to Fisher’s z, subtracted, and then transformed back to Pearson’s r. Alpha levels were adjusted to .005 (.05/10) following Bonferroni correction to account for the multiple comparisons within each column.

**Figure S6.**
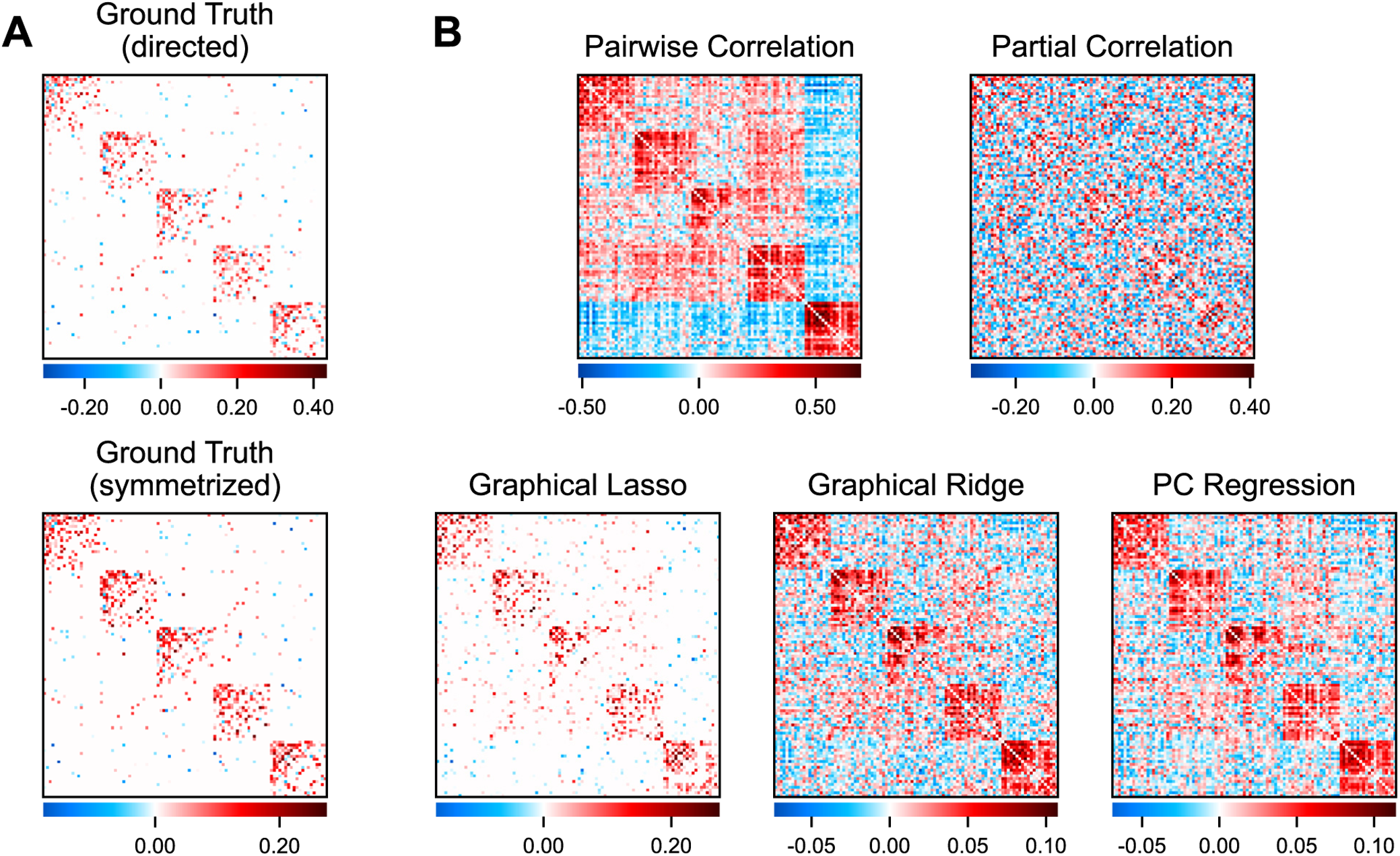
– Simulated networks and FC estimated by different methods – simulated activity convolved with HRF. **A)** An example ground truth network. The directed, asymmetric version (top) was used to generate nodes’ timeseries while the symmetrized version (bottom) was compared with FC estimates. **B)** Individual FC matrices estimated from a simulated timeseries using each FC method.

**Figure S7.**
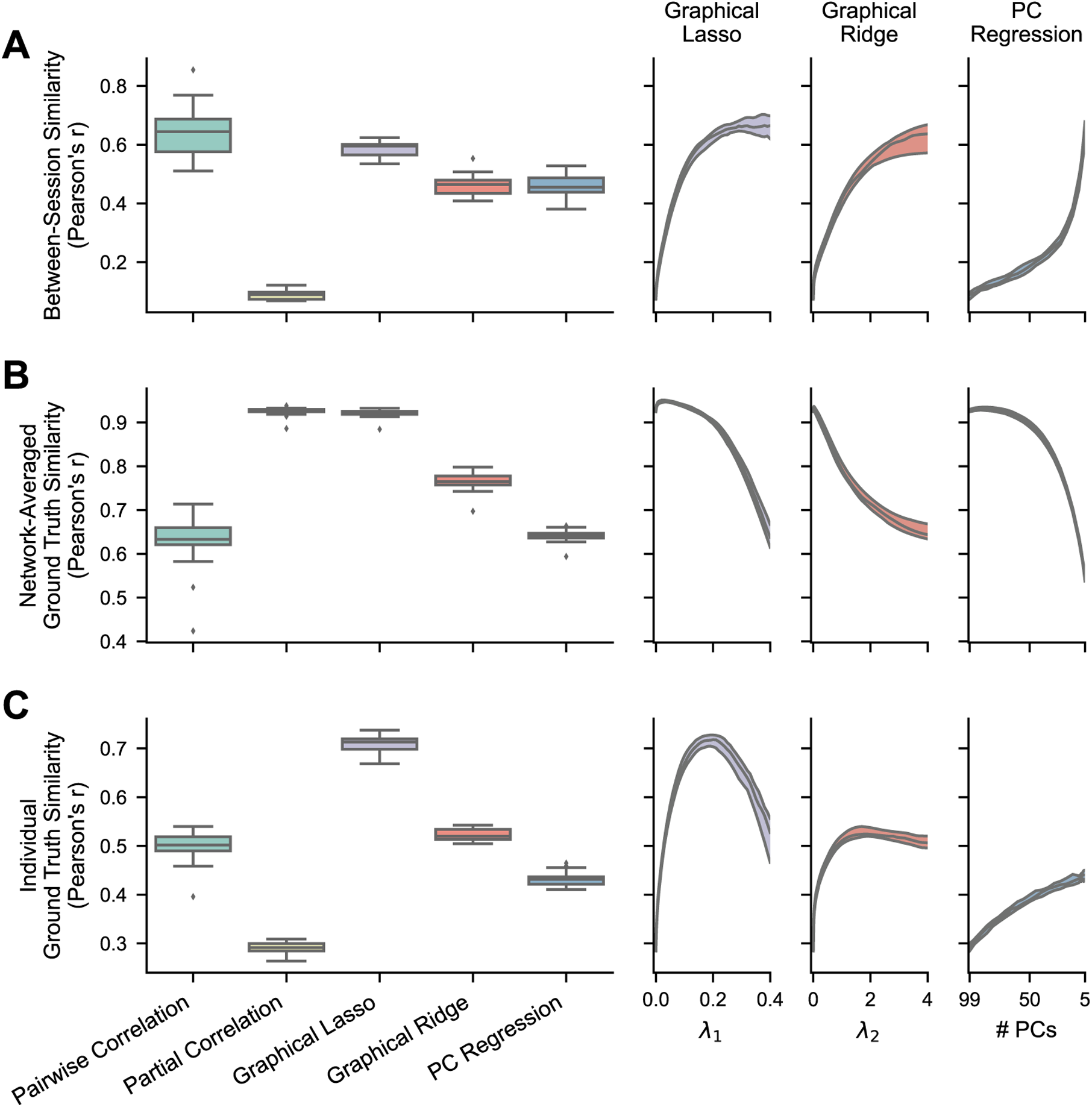
– Reliability and ground truth similarity of FC methods with simulated data – simulated activity convolved with HRF. The boxplots show results where the regularization hyperparameters have been optimized for each FC matrix, while the right plots show the medians and IQRs across different hyperparameter values for the regularized methods. For PC regression, number of PCs is plotted in descending order because fewer PCs correspond with more regularization. **A)** Between-session similarity, calculated between one pair of session matrices for each simulated network (n = 25). **B)** Ground truth similarity between group-averaged FC matrices (100 sessions each) and the ground truth for each simulated network (n = 25). Averaging nullifies much of the noise in individual connectivity weights to show validity without the effects of low reliability. **C)** Ground truth similarity between an individual session’s estimated FC matrix and the ground truth for each simulated network (n = 25). The accuracy of single measurements is vastly improved by recovering reliability through regularization.

**Table S4.**
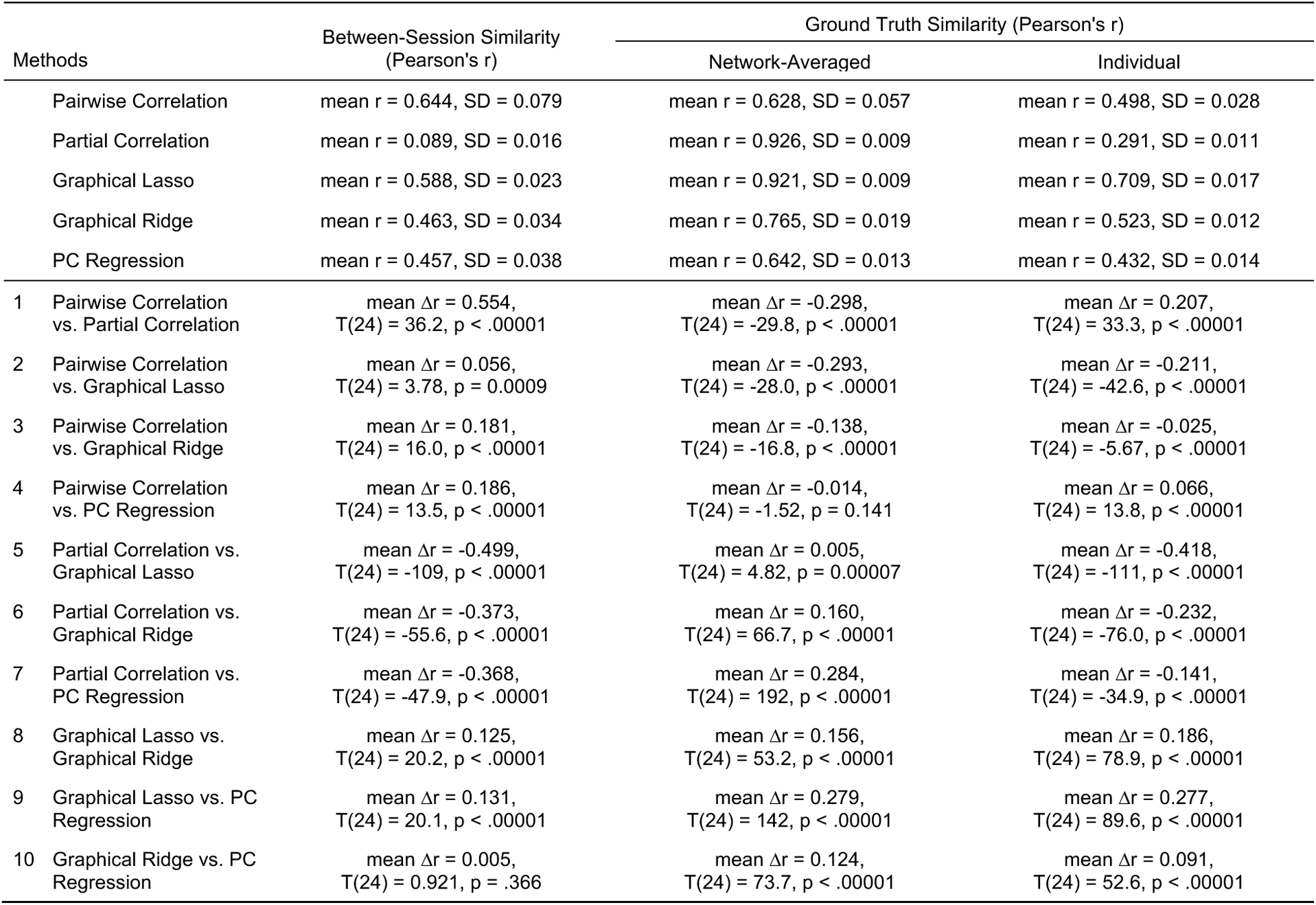
– Statistical tests comparing FC methods on reliability and ground truth similarity using simulated data – simulated activity convolved with HRF. Between-session similarity, group-averaged ground truth similarity, and individual ground truth similarity were compared using two-tailed, dependent-sample t-tests, their Pearson’s r values first normalized using Fisher’s z transformation (arctanh). For all reported correlation differences, correlations were transformed to Fisher’s z, subtracted, and then transformed back to Pearson’s r. Alpha levels were adjusted to .005 (.05/10) following Bonferroni correction to account for the multiple comparisons within each column.

**Figure S8.**
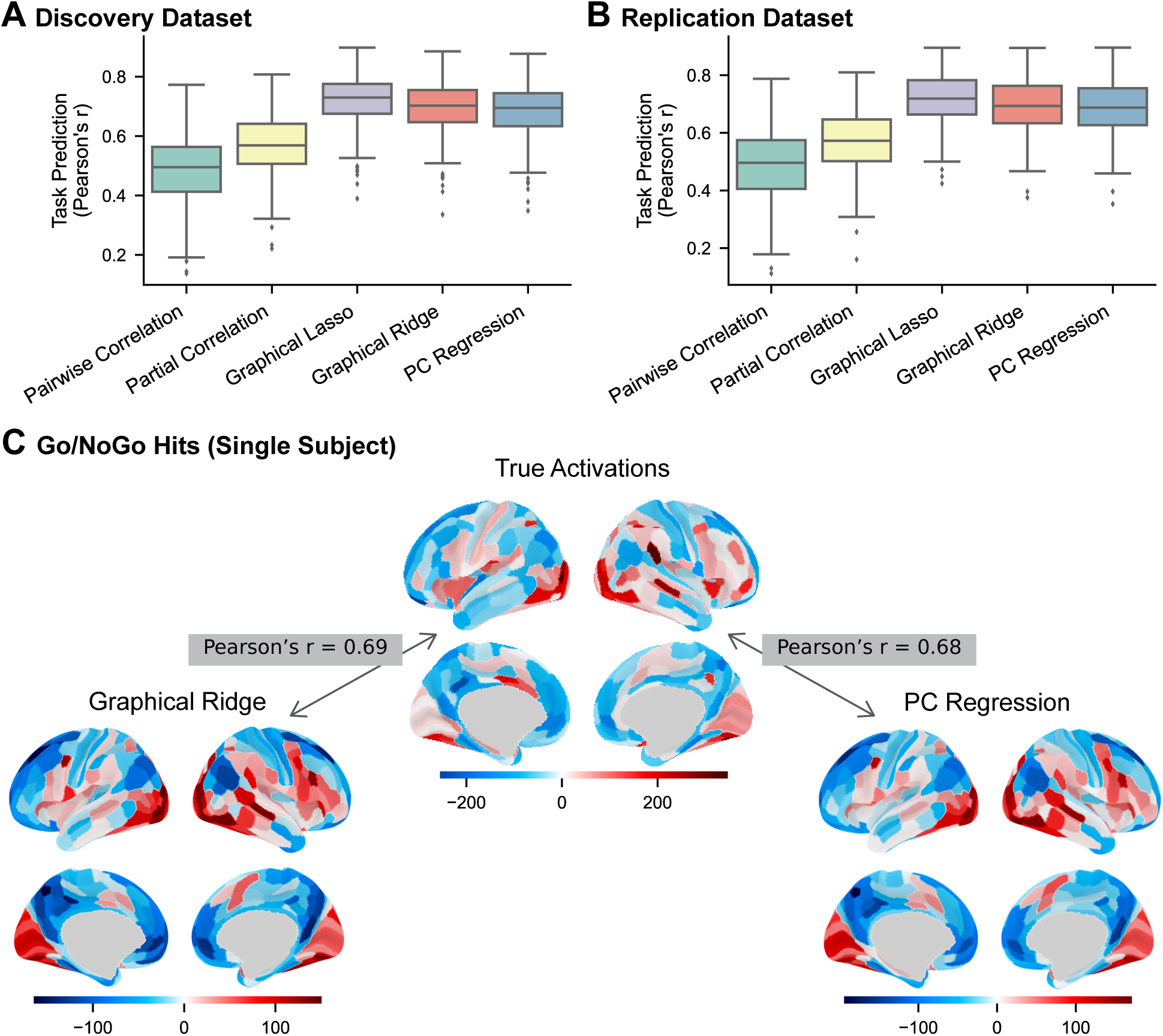
– Predicting task activations using FC estimated from rest fMRI data – discovery and replication datasets, all FC methods. **A-B**) Prediction accuracy by FC method calculated as the Pearson correlation (r) between regions’ actual and predicted activations, computed in each subject (n = 236) across regions and task conditions. Results are from the discovery dataset **(A)** and replication dataset **(B)**. **C)** A single subject’s actual and predicted (from graphical ridge and PC regression FC) task activations for the go/no-go hit events.

**Table S5.**
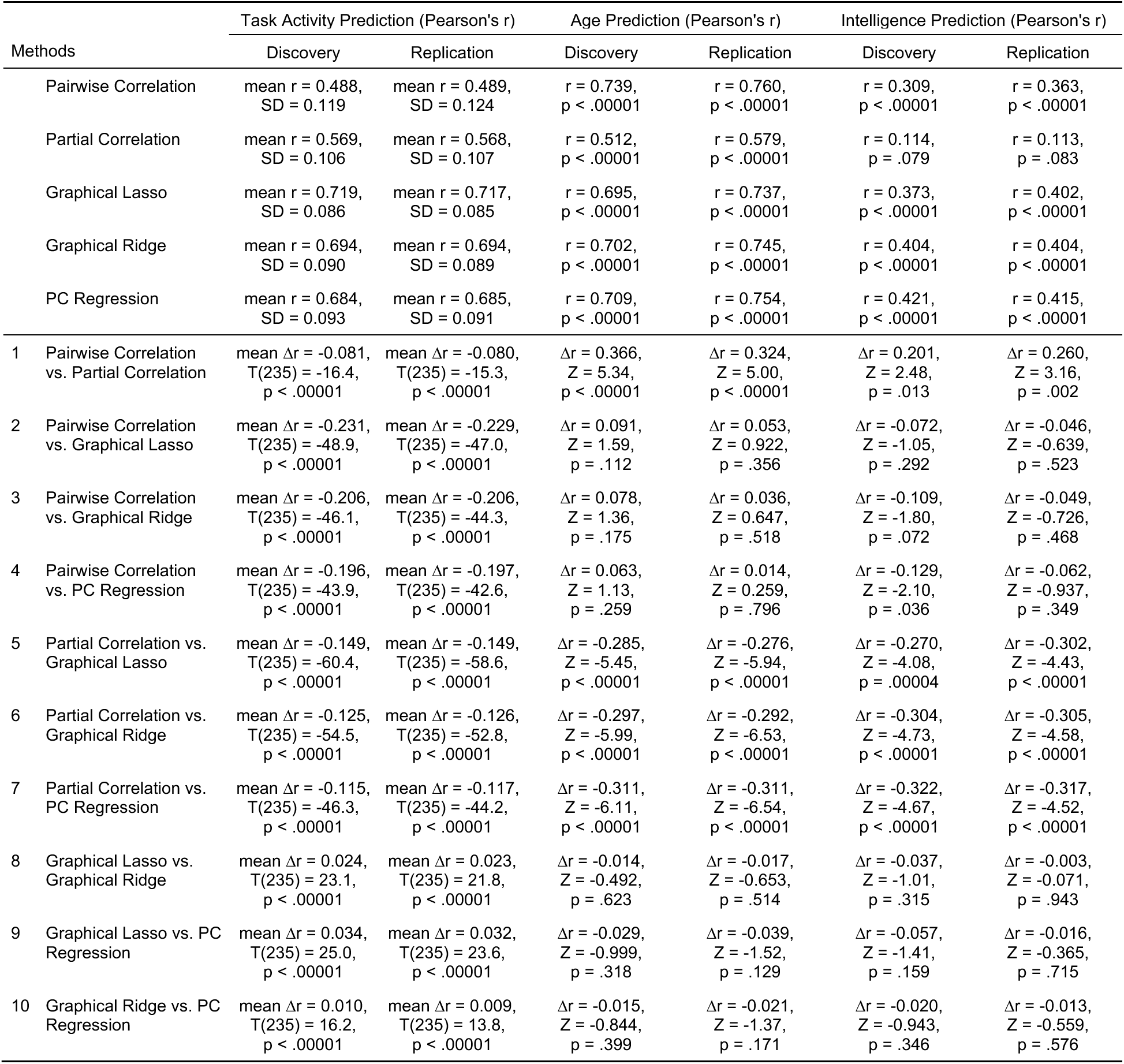
– Statistical tests comparing FC methods on task activation, subject age, and intelligence prediction accuracy – discovery and replication datasets. For task activation prediction accuracies, scores for all subjects (Pearson’s r) were compared using two-tailed, dependent-sample t-tests. For age and intelligence, predicted-to-actual correlations (single Pearson’s r per method) were compared between methods using the two-tailed test described by Meng, Rosenthal, and Rubin (1992), which tests for a significant difference between “correlated correlations” (correlations with a shared variable – actual subject age and psychometric *g* in this case). The Pearson’s r values were normalized using Fisher’s z transformation (arctanh) before statistical comparisons. For reported correlation differences, correlations were transformed to Fisher’s z, subtracted, and then transformed back to Pearson’s r. Alpha levels were adjusted to .005 (.05/10) following Bonferroni correction to account for the multiple comparisons within each column.

**Figure S9.**
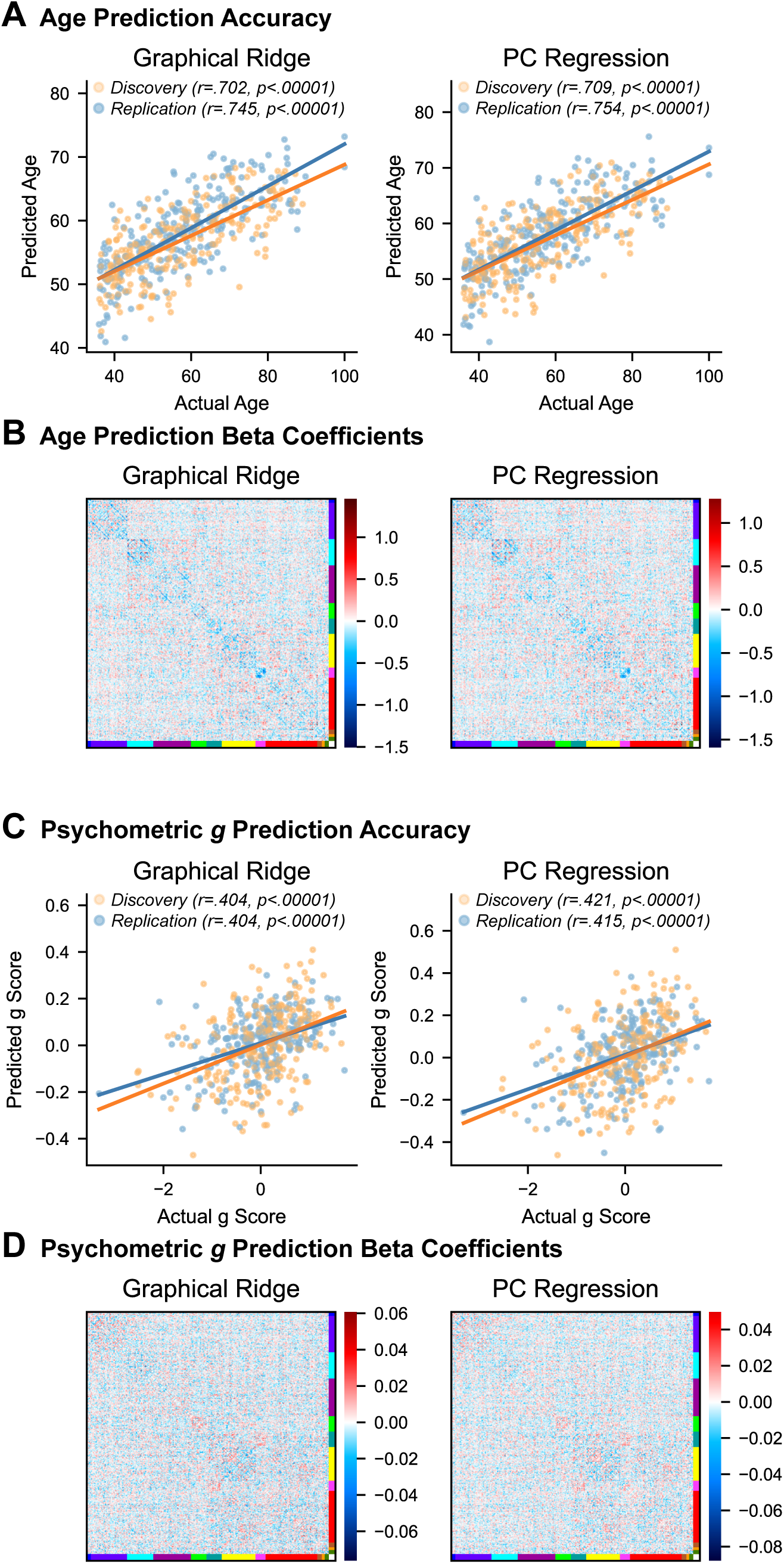
– Predicting individual differences in age and intelligence using estimated FC from rest fMRI data – graphical ridge and PC regression. **A)** Actual and predicted ages of each subject by FC method, from both the discovery and replication datasets. **B)** Average beta coefficients assigned to each connection by the regression models for estimating subject age. Beta coefficients shown here are the averages over cross-validation folds and discovery and replication datasets. Blue indicates that connection strength decreased with age and red indicates that connection strength increased. **C)** Actual and predicted intelligence (psychometric *g*) of each subject by FC method. **D)** Average beta coefficients assigned to each connection by the regression models for estimating psychometric *g*. See Figure 9 for results using pairwise correlation, partial correlation, and graphical lasso for FC estimation.

