## Supplementary Materials for "Regularized partial correlation provides reliable functional connectivity estimates while correcting for widespread confounding"

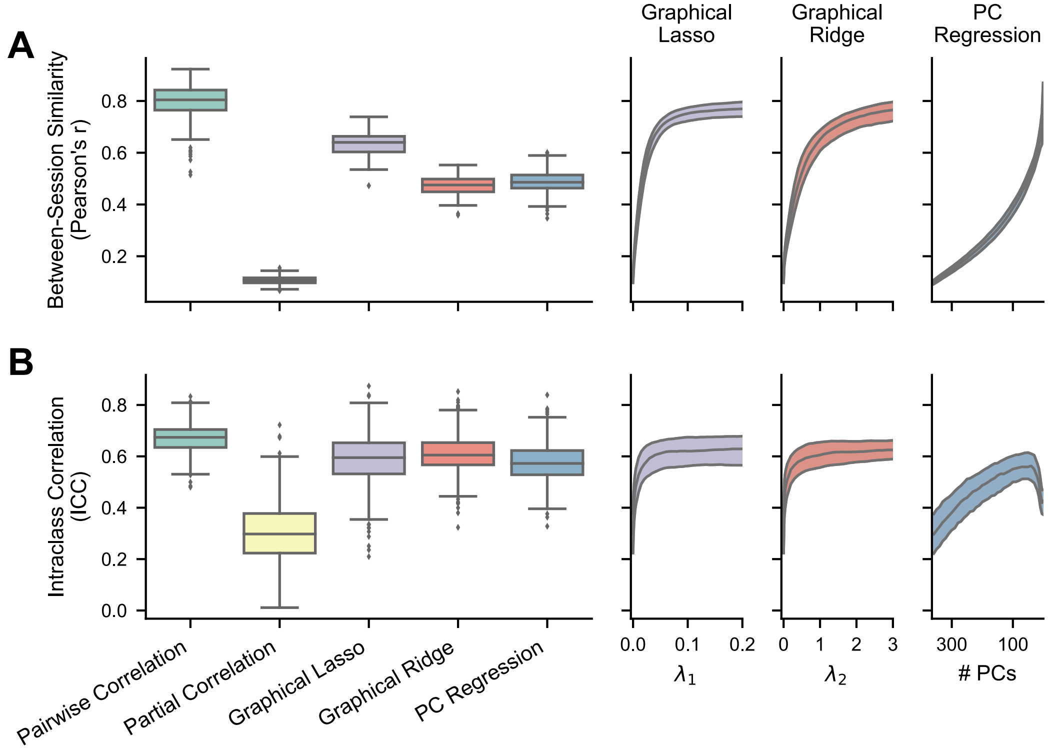


**Figure S1 – Reliability of FC methods with empirical fMRI data – replication dataset.** The boxplots show results where the regularization hyperparameters have been optimized for each FC matrix, while the right plots show the medians and IQRs across different hyperparameter values for the regularized methods. For PC regression, number of PCs is plotted in descending order because fewer PCs correspond with more regularization. **A)** Between-session similarity, the Pearson correlation between each subject's session 1 and session 2 FC matrices (n = 236). **B)** Intraclass correlation (Shrout & Fleiss, 1979), calculated for each edge in a conservative subset of nonzero edges (n = 532).

**Table S1 – Statistical tests comparing FC methods on reliability and structural-functional similarity using empirical data – replication dataset.** Between-session similarity, intraclass correlation, and individual structural-functional similarity were compared using two-tailed, dependent-sample t-tests. The Pearson's r values for between-session similarity and individual structural-functional similarity were normalized using Fisher's z transformation (arctanh) before statistical comparisons. Group-averaged structural-functional similarity (single Pearson's r per method) was compared using the method described by Meng et al. (1992), which tests for a significant difference between "correlated correlations" (correlations with a shared variable – SC weights in this case). For all reported correlation differences, correlations were transformed to Fisher's z, subtracted, and then transformed back to Pearson's r. Alpha levels were adjusted to .005 (.05/10) following Bonferroni correction to account for the multiple comparisons within each column.

| Methods | | Between-Session Similarity (Pearson's r) | Intraclass Correlation (ICC) | Structural-Functional Similarity (Pearson's r) | |
|---|---|---|---|---|---|
| | | | | Group-Averaged | Individual |
| | Pairwise Correlation | mean r = 0.797, SD = 0.067 | mean ICC = 0.669, SD = 0.054 | r = 0.250 | mean r = 0.167, SD = 0.020 |
| | Partial Correlation | mean r = 0.107, SD = 0.015 | mean ICC = 0.302, SD = 0.111 | r = 0.570 | mean r = 0.135, SD = 0.012 |
| | Graphical Lasso | mean r = 0.632, SD = 0.047 | mean ICC = 0.588, SD = 0.096 | r = 0.542 | mean r = 0.318, SD = 0.022 |
| | Graphical Ridge | mean r = 0.474, SD = 0.036 | mean ICC = 0.607, SD = 0.075 | r = 0.443 | mean r = 0.218, SD = 0.018 |
| | PC Regression | mean r = 0.487, SD = 0.042 | mean ICC = 0.574, SD = 0.074 | r = 0.397 | mean r = 0.195, SD = 0.019 |
| 1 | Pairwise Correlation vs. Partial Correlation | mean Δr = 0.765, T(235) = 86.4, p < .00001 | mean ΔICC = 0.367, T(531) = 77.6, p < .00001 | Δr = -0.372, Z = -5.31, p < .00001 | mean Δr = 0.032, T(235) = 29.3, p < .00001 |
| 2 | Pairwise Correlation vs. Graphical Lasso | mean Δr = 0.351, T(235) = 32.1, p < .00001 | mean ΔICC = 0.081, T(531) = 21.0, p < .00001 | Δr = -0.338, Z = -4.97, p < .00001 | mean Δr = -0.151, T(235) = -124, p < .00001 |
| 3 | Pairwise Correlation vs. Graphical Ridge | mean Δr = 0.537, T(235) = 54.3, p < .00001 | mean ΔICC = 0.062, T(531) = 20.5, p < .00001 | Δr = -0.217, Z = -3.80, p = 0.0001 | mean Δr = -0.051, T(235) = -52.4, p < .00001 |
| 4 | Pairwise Correlation vs. PC Regression | mean Δr = 0.524, T(235) = 54.4, p < .00001 | mean ΔICC = 0.094, T(531) = 30.9, p < .00001 | Δr = -0.163, Z = -2.93, p = 0.003 | mean Δr = -0.028, T(235) = -29.5, p < .00001 |
| 5 | Partial Correlation vs. Graphical Lasso | mean Δr = -0.566, T(235) = -140, p < .00001 | mean ΔICC = -0.286, T(531) = -82.5, p < .00001 | Δr = 0.039, Z = 1.49, p = 0.137 | mean Δr = -0.183, T(235) = -181, p < .00001 |
| 6 | Partial Correlation vs. Graphical Ridge | mean Δr = -0.388, T(235) = -164, p < .00001 | mean ΔICC = -0.305, T(531) = -71.3, p < .00001 | Δr = 0.169, Z = 4.24, p = 0.00002 | mean Δr = -0.083, T(235) = -121, p < .00001 |
| 7 | Partial Correlation vs. PC Regression | mean Δr = -0.402, T(235) = -116, p < .00001 | mean ΔICC = -0.273, T(531) = -57.9, p < .00001 | Δr = 0.223, Z = 4.73, p < .00001 | mean Δr = -0.060, T(235) = -84.8, p < .00001 |
| 8 | Graphical Lasso vs. Graphical Ridge | mean Δr = 0.229, T(235) = 79.4, p < .00001 | mean ΔICC = -0.019, T(531) = -7.01, p < .00001 | Δr = 0.131, Z = 3.40, p = 0.0007 | mean Δr = 0.100, T(235) = 129, p < .00001 |
| 9 | Graphical Lasso vs. PC Regression | mean Δr = 0.212, T(235) = 54.1, p < .00001 | mean ΔICC = 0.013, T(531) = 4.03, p < .00001 | Δr = 0.185, Z = 4.15, p = 0.00003 | mean Δr = 0.123, T(235) = 146, p < .00001 |
| 10 | Graphical Ridge vs. PC Regression | mean Δr = -0.018, T(235) = -6.48, p < .00001 | mean ΔICC = 0.032, T(531) = 21.1, p < .00001 | Δr = 0.055, Z = 5.30, p < .00001 | mean Δr = 0.023, T(235) = 96.0, p < .00001 |

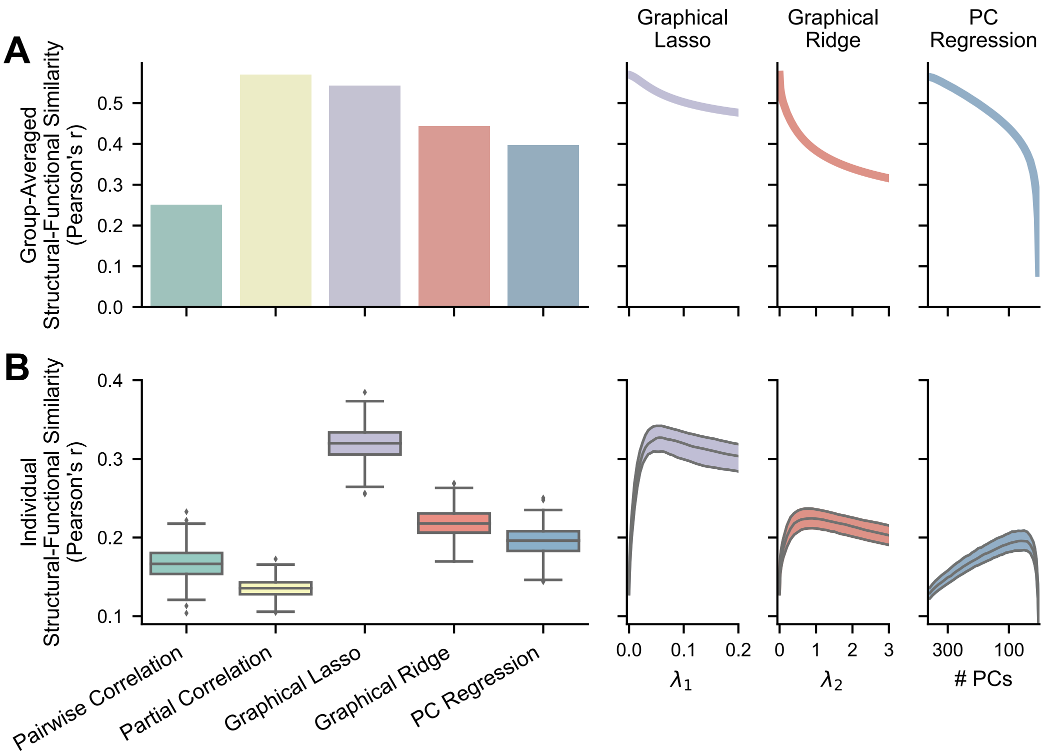


**Figure S2 - Similarity between empirical FC and SC (diffusion MRI tractography) – replication dataset.** The bar and boxplots show results where the regularization hyperparameters have been optimized for each FC matrix, while the right plots show the single values or the medians and IQRs across different hyperparameter values for the regularized methods. For PC regression, number of PCs is plotted in descending order because fewer PCs correspond with more regularization. **A)** Structural-functional similarity between group-averaged FC and structural matrices (n = 1). Averaging nullifies much of the noise in individual connectivity weights to show validity without the effects of low reliability. **B)** Structural-functional similarity between individual FC matrices and the same subjects' structural matrices (n = 236). Individual measurement accuracy is vastly improved by recovering reliability.

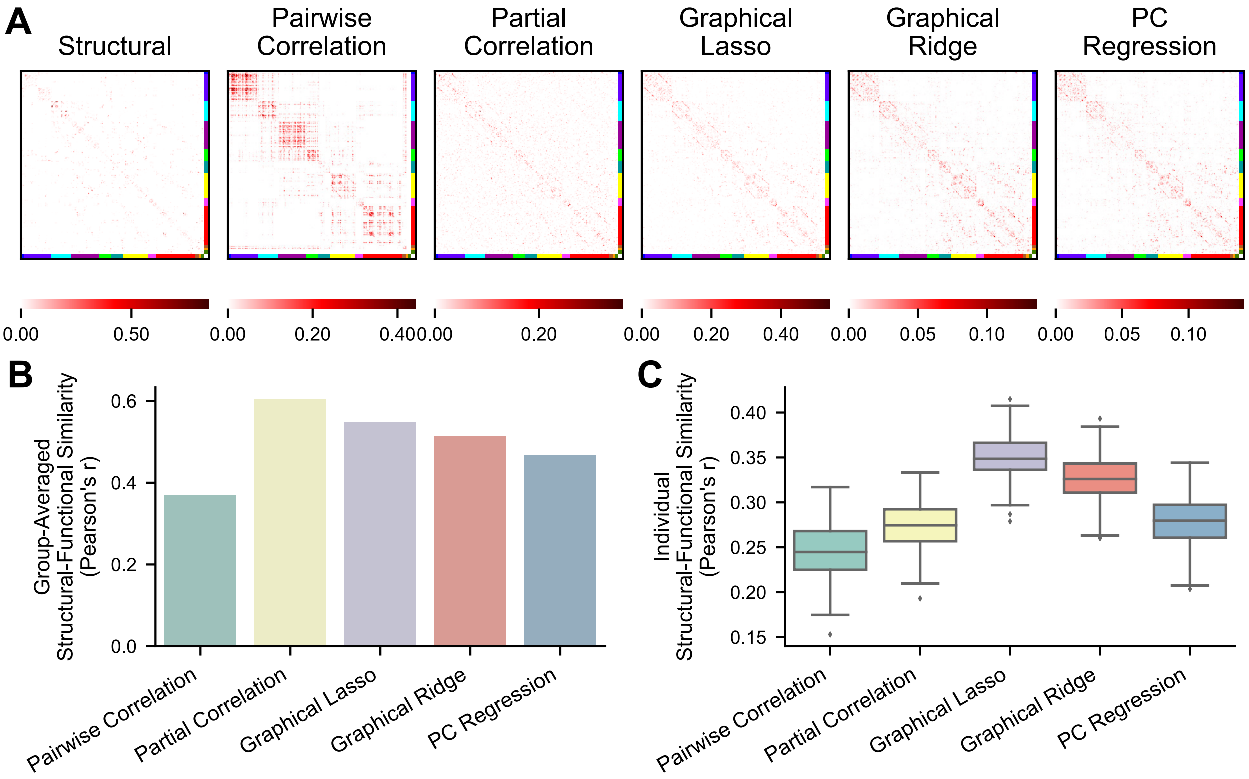


**Figure S3 - Similarity between empirical FC and SC (diffusion MRI tractography) – sparsity-matched FC. A)** Single subject's SC matrix, and FC matrices thresholded to have the same sparsity as SC. **B)** Structural-functional similarity between group-averaged sparsity-matched FC and structural matrices (n = 1). Averaging nullifies much of the noise in individual connectivity weights to show validity without the effects of low reliability. **C)** Structural-functional similarity between individual sparsity-matched FC matrices and the same subjects' structural matrices (n = 236). Individual measurement accuracy is vastly improved by recovering reliability.

**Table S2 – Statistical tests comparing FC methods on structural-functional similarity with sparsity-matched FC.** Individual structural-functional similarity was compared using two-tailed, dependent-sample t-tests. The Pearson's r values for individual structural-functional similarity were normalized using Fisher's z transformation (arctanh) before statistical comparisons. Group-averaged structural-functional similarity (single Pearson's r per method) was compared using the method described by Meng et al. (1992), which tests for a significant difference between "correlated correlations" (correlations with a shared variable – SC weights in this case). For all reported correlation differences, correlations were transformed to Fisher's z, subtracted, and then transformed back to Pearson's r. Alpha levels were adjusted to .005 (.05/10) following Bonferroni correction to account for the multiple comparisons within each column.

| | | Structural-Functional Similarity (Pearson's r) | |
|---|---|---|---|
| Methods | | Group-Averaged | Individual |
| | Pairwise Correlation | r = 0.370 | mean r = 0.246, SD = 0.030 |
| | Partial Correlation | r = 0.603 | mean r = 0.274, SD = 0.026 |
| | Graphical Lasso | r = 0.549 | mean r = 0.350, SD = 0.024 |
| | Graphical Ridge | r = 0.514 | mean r = 0.326, SD = 0.025 |
| | PC Regression | r = 0.467 | mean r = 0.278, SD = 0.029 |
| **1** | Pairwise Correlation vs. Partial Correlation | Δr = -0.300, Z = -4.58, p < .00001 | mean Δr = -0.028, T(235) = -16.1, p < .00001 |
| **2** | Pairwise Correlation vs. Graphical Lasso | Δr = -0.224, Z = -3.83, p = 0.0001 | mean Δr = -0.104, T(235) = -63.9, p < .00001 |
| **3** | Pairwise Correlation vs. Graphical Ridge | Δr = -0.178, Z = -3.04, p = 0.002 | mean Δr = -0.079, T(235) = -38.7, p < .00001 |
| **4** | Pairwise Correlation vs. PC Regression | Δr = -0.117, Z = -2.15, p = 0.032 | mean Δr = -0.032, T(235) = -16.9, p < .00001 |
| **5** | Partial Correlation vs. Graphical Lasso | Δr = 0.081, Z = 2.80, p = 0.005 | mean Δr = -0.076, T(235) = -58.0, p < .00001 |
| **6** | Partial Correlation vs. Graphical Ridge | Δr = 0.128, Z = 3.07, p = 0.002 | mean Δr = -0.051, T(235) = -29.8, p < .00001 |
| **7** | Partial Correlation vs. PC Regression | Δr = 0.190, Z = 3.82, p = 0.0001 | mean Δr = -0.004, T(235) = -2.64, p = 0.009 |
| **8** | Graphical Lasso vs. Graphical Ridge | Δr = 0.048, Z = 2.01, p = 0.044 | mean Δr = 0.025, T(235) = 23.7, p < .00001 |
| **9** | Graphical Lasso vs. PC Regression | Δr = 0.110, Z = 3.51, p = 0.0004 | mean Δr = 0.072, T(235) = 59.6, p < .00001 |
| **10** | Graphical Ridge vs. PC Regression | Δr = 0.063, Z = 4.15, p = 0.00003 | mean Δr = 0.047, T(235) = 59.4, p < .00001 |

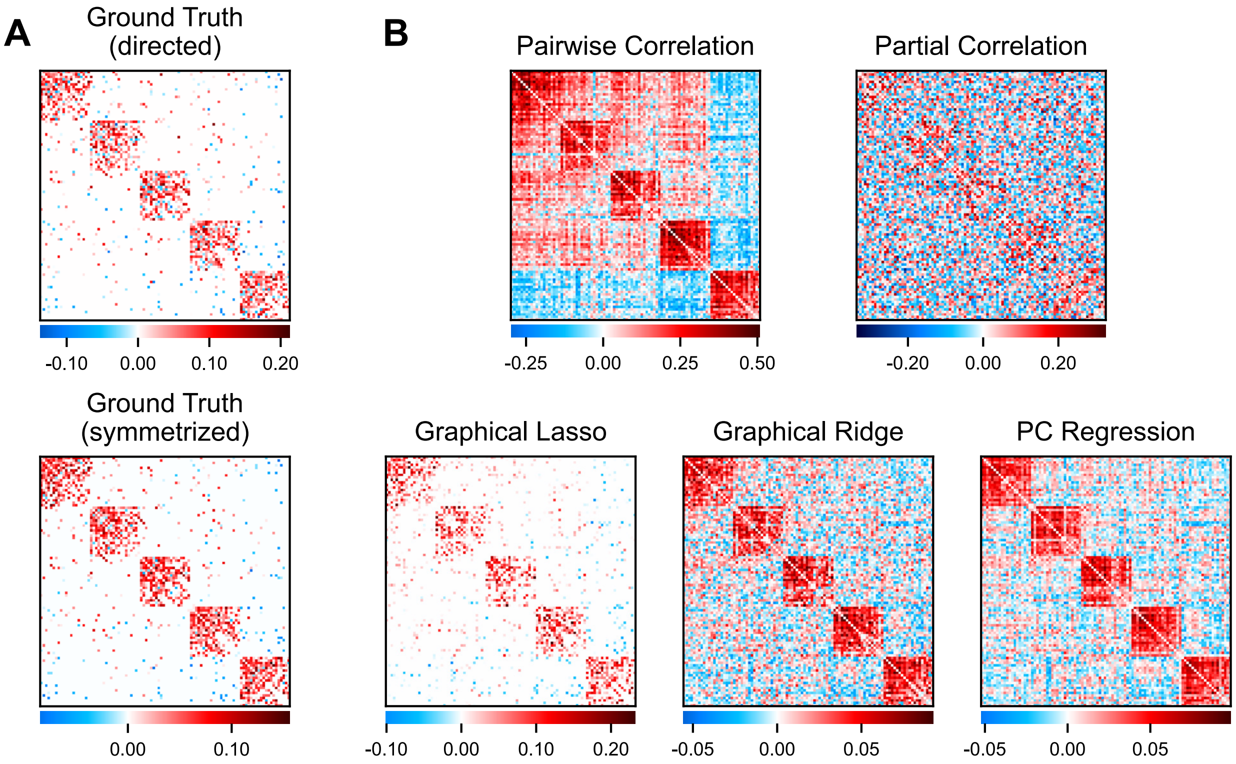


**Figure S4 – Simulated networks and FC estimated by different methods – less sparse networks. A)** An example ground truth network. The directed, asymmetric version (top) was used to generate nodes' timeseries while the symmetrized version (bottom) was compared with FC estimates. **B)** Individual FC matrices estimated from a simulated timeseries using each FC method.

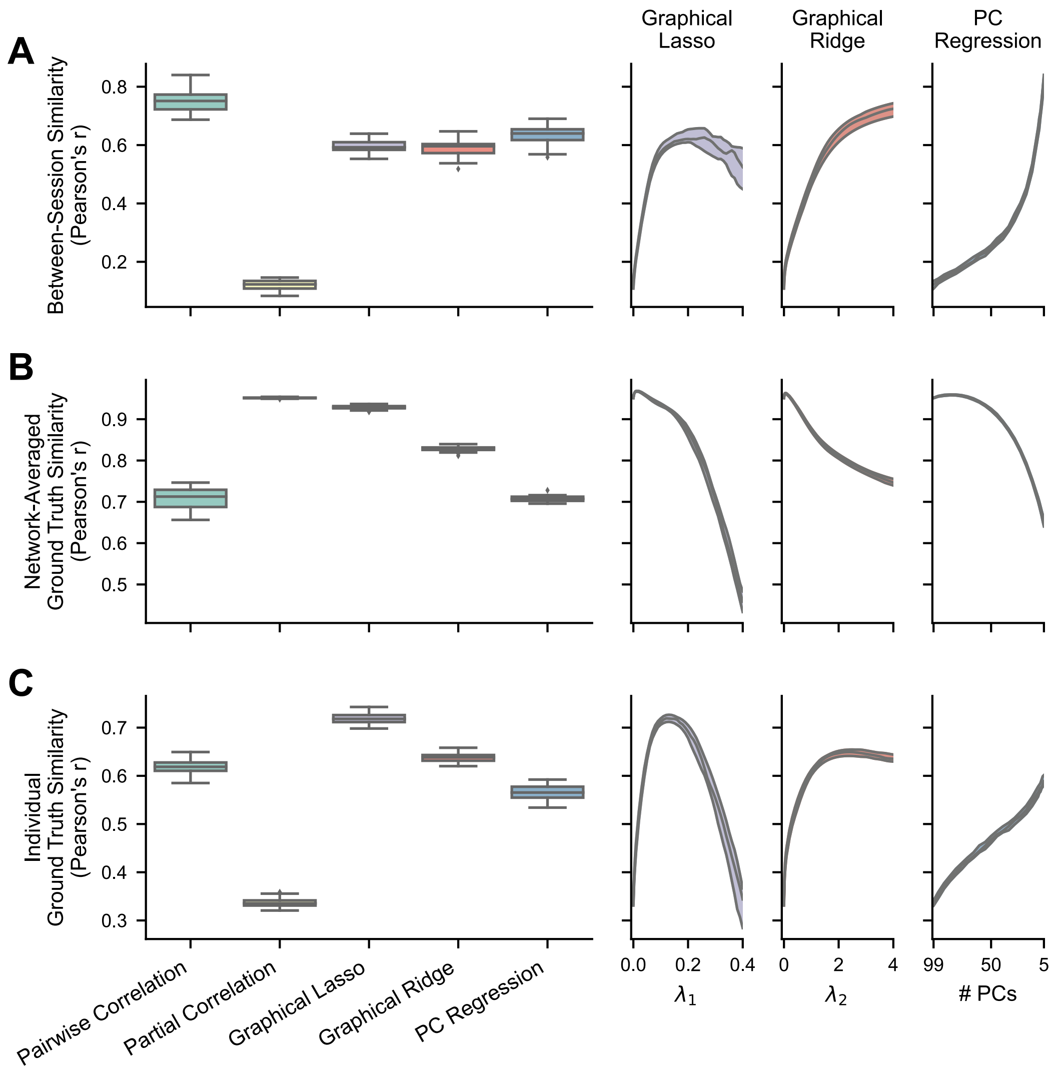


**Figure S5 – Reliability and ground truth similarity of FC methods with simulated data – less sparse networks.** The boxplots show results where the regularization hyperparameters have been optimized for each FC matrix, while the right plots show the medians and IQRs across different hyperparameter values for the regularized methods. For PC regression, number of PCs is plotted in descending order because fewer PCs correspond with more regularization. **A)** Between-session similarity, calculated between one pair of session matrices for each simulated network (n = 25). **B)** Ground truth similarity between group-averaged FC matrices (100 sessions each) and the ground truth for each simulated network (n = 25). Averaging nullifies much of the noise in individual connectivity weights to show validity without the effects of low reliability. **C)** Ground truth similarity between an individual session's estimated FC matrix and the ground truth for each simulated network (n = 25). The accuracy of single measurements is vastly improved by recovering reliability through regularization.

**Table S3 – Statistical tests comparing FC methods on reliability and ground truth similarity using simulated data – less sparse networks.** Between-session similarity, group-averaged ground truth similarity, and individual ground truth similarity were compared using two-tailed, dependent-sample t-tests, their Pearson's r values first normalized using Fisher's z transformation (arctanh). For all reported correlation differences, correlations were transformed to Fisher's z, subtracted, and then transformed back to Pearson's r. Alpha levels were adjusted to .005 (.05/10) following Bonferroni correction to account for the multiple comparisons within each column.

| | Methods | Between-Session Similarity (Pearson's r) | Ground Truth Similarity (Pearson's r) | |
|---|---|---|---|---|
| | | | Network-Averaged | Individual |
| | Pairwise Correlation | mean r = 0.751, SD = 0.041 | mean r = 0.710, SD = 0.025 | mean r = 0.618, SD = 0.016 |
| | Partial Correlation | mean r = 0.120, SD = 0.017 | mean r = 0.951, SD = 0.001 | mean r = 0.337, SD = 0.010 |
| | Graphical Lasso | mean r = 0.595, SD = 0.022 | mean r = 0.929, SD = 0.005 | mean r = 0.719, SD = 0.010 |
| | Graphical Ridge | mean r = 0.588, SD = 0.028 | mean r = 0.828, SD = 0.006 | mean r = 0.638, SD = 0.009 |
| | PC Regression | mean r = 0.633, SD = 0.032 | mean r = 0.707, SD = 0.007 | mean r = 0.565, SD = 0.015 |
| 1 | Pairwise Correlation vs. Partial Correlation | mean Δr = 0.631, T(24) = 85.7, $p < .00001$ | mean Δr = -0.241, T(24) = -47.2, $p < .00001$ | mean Δr = 0.282, T(24) = 72.2, $p < .00001$ |
| 2 | Pairwise Correlation vs. Graphical Lasso | mean Δr = 0.156, T(24) = 25.3, $p < .00001$ | mean Δr = -0.218, T(24) = -43.6, $p < .00001$ | mean Δr = -0.101, T(24) = -26.7, $p < .00001$ |
| 3 | Pairwise Correlation vs. Graphical Ridge | mean Δr = 0.163, T(24) = 33.2, $p < .00001$ | mean Δr = -0.117, T(24) = -26.9, $p < .00001$ | mean Δr = -0.020, T(24) = -7.12, $p < .00001$ |
| 4 | Pairwise Correlation vs. PC Regression | mean Δr = 0.118, T(24) = 15.3, $p < .00001$ | mean Δr = 0.003, T(24) = 0.609, $p = .548$ | mean Δr = 0.053, T(24) = 15.0, $p < .00001$ |
| 5 | Partial Correlation vs. Graphical Lasso | mean Δr = -0.475, T(24) = -96.8, $p < .00001$ | mean Δr = 0.023, T(24) = 25.9, $p < .00001$ | mean Δr = -0.383, T(24) = -175, $p < .00001$ |
| 6 | Partial Correlation vs. Graphical Ridge | mean Δr = -0.468, T(24) = -95.6, $p < .00001$ | mean Δr = 0.124, T(24) = 98.9, $p < .00001$ | mean Δr = -0.302, T(24) = -128, $p < .00001$ |
| 7 | Partial Correlation vs. PC Regression | mean Δr = -0.513, T(24) = -85.5, $p < .00001$ | mean Δr = 0.244, T(24) = 157, $p < .00001$ | mean Δr = -0.228, T(24) = -61.2, $p < .00001$ |
| 8 | Graphical Lasso vs. Graphical Ridge | mean Δr = 0.007, T(24) = 1.73, $p = .096$ | mean Δr = 0.101, T(24) = 62.6, $p < .00001$ | mean Δr = 0.081, T(24) = 36.4, $p < .00001$ |
| 9 | Graphical Lasso vs. PC Regression | mean Δr = -0.038, T(24) = -6.02, $p < .00001$ | mean Δr = 0.221, T(24) = 130, $p < .00001$ | mean Δr = 0.154, T(24) = 47.9, $p < .00001$ |
| 10 | Graphical Ridge vs. PC Regression | mean Δr = -0.045, T(24) = -8.56, $p < .00001$ | mean Δr = 0.120, T(24) = 72.5, $p < .00001$ | mean Δr = 0.074, T(24) = 33.1, $p < .00001$ |

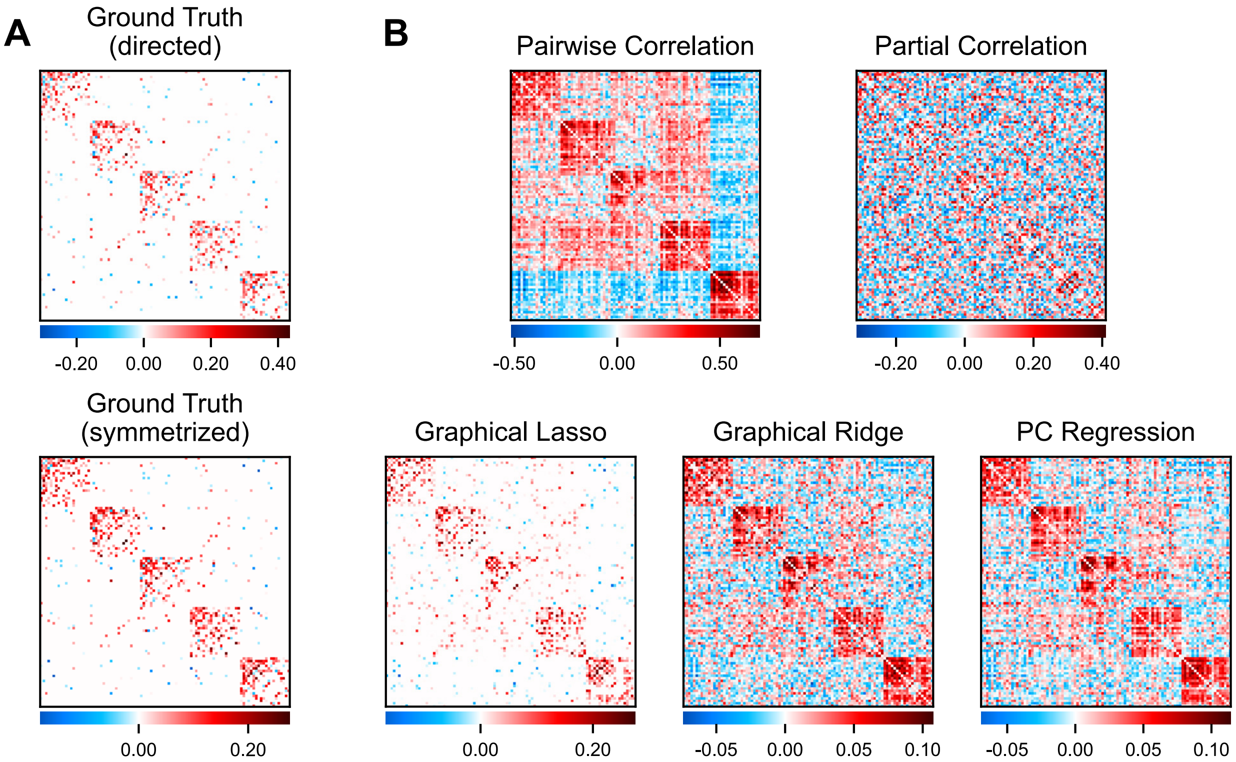


**Figure S6 – Simulated networks and FC estimated by different methods – simulated activity convolved with HRF. A)** An example ground truth network. The directed, asymmetric version (top) was used to generate nodes' timeseries while the symmetrized version (bottom) was compared with FC estimates. **B)** Individual FC matrices estimated from a simulated timeseries using each FC method.

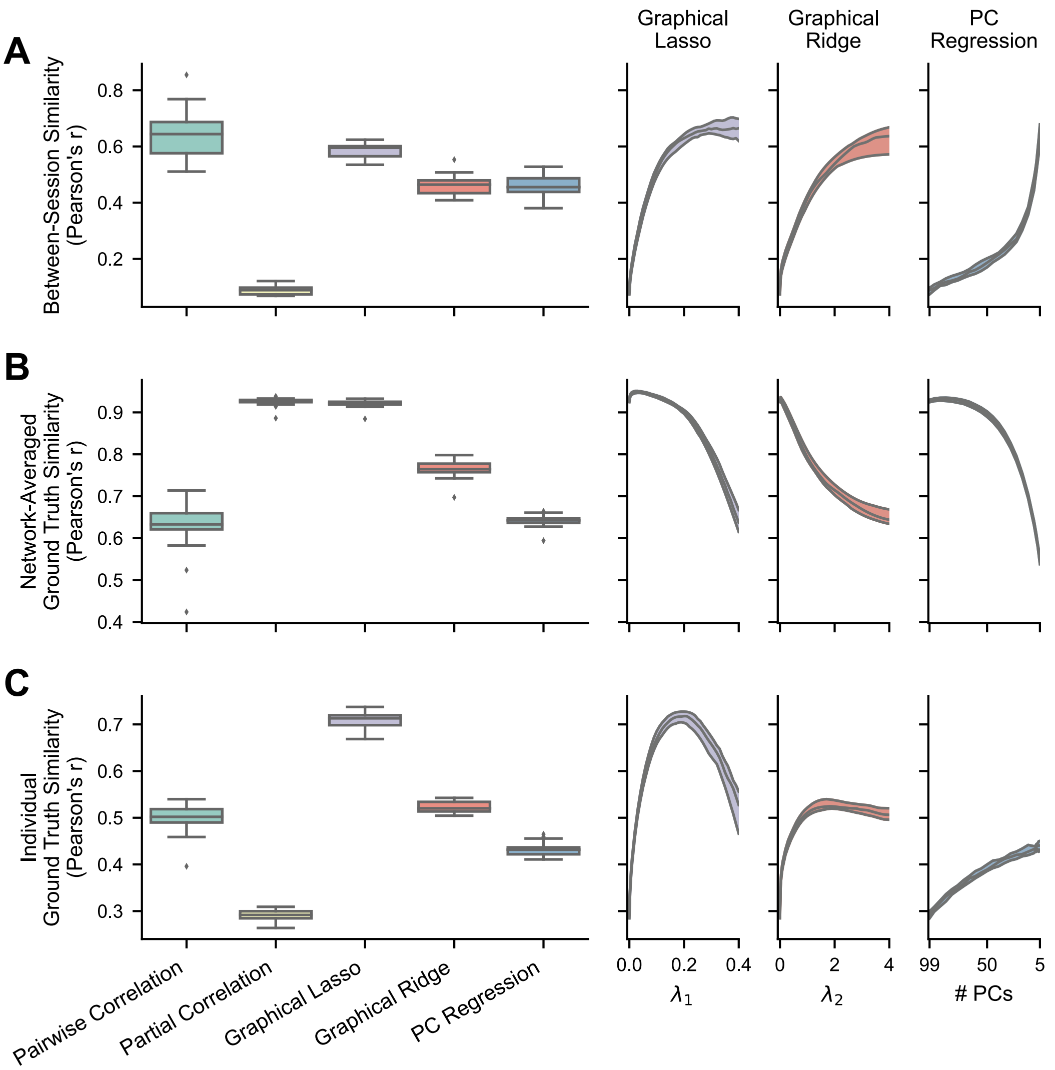


**Figure S7 – Reliability and ground truth similarity of FC methods with simulated data – simulated activity convolved with HRF.** The boxplots show results where the regularization hyperparameters have been optimized for each FC matrix, while the right plots show the medians and IQRs across different hyperparameter values for the regularized methods. For PC regression, number of PCs is plotted in descending order because fewer PCs correspond with more regularization. **A)** Between-session similarity, calculated between one pair of session matrices for each simulated network (n = 25). **B)** Ground truth similarity between group-averaged FC matrices (100 sessions each) and the ground truth for each simulated network (n = 25). Averaging nullifies much of the noise in individual connectivity weights to show validity without the effects of low reliability. **C)** Ground truth similarity between an individual session's estimated FC matrix and the ground truth for each simulated network (n = 25). The accuracy of single measurements is vastly improved by recovering reliability through regularization.

**Table S4 – Statistical tests comparing FC methods on reliability and ground truth similarity using simulated data – simulated activity convolved with HRF.** Between-session similarity, group-averaged ground truth similarity, and individual ground truth similarity were compared using two-tailed, dependent-sample t-tests, their Pearson's r values first normalized using Fisher's z transformation (arctanh). For all reported correlation differences, correlations were transformed to Fisher's z, subtracted, and then transformed back to Pearson's r. Alpha levels were adjusted to .005 (.05/10) following Bonferroni correction to account for the multiple comparisons within each column.

| | Methods | Between-Session Similarity (Pearson's r) | Ground Truth Similarity (Pearson's r) | |
|---|---|---|---|---|
| | | | Network-Averaged | Individual |
| | Pairwise Correlation | mean r = 0.644, SD = 0.079 | mean r = 0.628, SD = 0.057 | mean r = 0.498, SD = 0.028 |
| | Partial Correlation | mean r = 0.089, SD = 0.016 | mean r = 0.926, SD = 0.009 | mean r = 0.291, SD = 0.011 |
| | Graphical Lasso | mean r = 0.588, SD = 0.023 | mean r = 0.921, SD = 0.009 | mean r = 0.709, SD = 0.017 |
| | Graphical Ridge | mean r = 0.463, SD = 0.034 | mean r = 0.765, SD = 0.019 | mean r = 0.523, SD = 0.012 |
| | PC Regression | mean r = 0.457, SD = 0.038 | mean r = 0.642, SD = 0.013 | mean r = 0.432, SD = 0.014 |
| 1 | Pairwise Correlation vs. Partial Correlation | mean Δr = 0.554, T(24) = 36.2, p < .00001 | mean Δr = -0.298, T(24) = -29.8, p < .00001 | mean Δr = 0.207, T(24) = 33.3, p < .00001 |
| 2 | Pairwise Correlation vs. Graphical Lasso | mean Δr = 0.056, T(24) = 3.78, p = 0.0009 | mean Δr = -0.293, T(24) = -28.0, p < .00001 | mean Δr = -0.211, T(24) = -42.6, p < .00001 |
| 3 | Pairwise Correlation vs. Graphical Ridge | mean Δr = 0.181, T(24) = 16.0, p < .00001 | mean Δr = -0.138, T(24) = -16.8, p < .00001 | mean Δr = -0.025, T(24) = -5.67, p < .00001 |
| 4 | Pairwise Correlation vs. PC Regression | mean Δr = 0.186, T(24) = 13.5, p < .00001 | mean Δr = -0.014, T(24) = -1.52, p = 0.141 | mean Δr = 0.066, T(24) = 13.8, p < .00001 |
| 5 | Partial Correlation vs. Graphical Lasso | mean Δr = -0.499, T(24) = -109, p < .00001 | mean Δr = 0.005, T(24) = 4.82, p = 0.00007 | mean Δr = -0.418, T(24) = -111, p < .00001 |
| 6 | Partial Correlation vs. Graphical Ridge | mean Δr = -0.373, T(24) = -55.6, p < .00001 | mean Δr = 0.160, T(24) = 66.7, p < .00001 | mean Δr = -0.232, T(24) = -76.0, p < .00001 |
| 7 | Partial Correlation vs. PC Regression | mean Δr = -0.368, T(24) = -47.9, p < .00001 | mean Δr = 0.284, T(24) = 192, p < .00001 | mean Δr = -0.141, T(24) = -34.9, p < .00001 |
| 8 | Graphical Lasso vs. Graphical Ridge | mean Δr = 0.125, T(24) = 20.2, p < .00001 | mean Δr = 0.156, T(24) = 53.2, p < .00001 | mean Δr = 0.186, T(24) = 78.9, p < .00001 |
| 9 | Graphical Lasso vs. PC Regression | mean Δr = 0.131, T(24) = 20.1, p < .00001 | mean Δr = 0.279, T(24) = 142, p < .00001 | mean Δr = 0.277, T(24) = 89.6, p < .00001 |
| 10 | Graphical Ridge vs. PC Regression | mean Δr = 0.005, T(24) = 0.921, p = .366 | mean Δr = 0.124, T(24) = 73.7, p < .00001 | mean Δr = 0.091, T(24) = 52.6, p < .00001 |

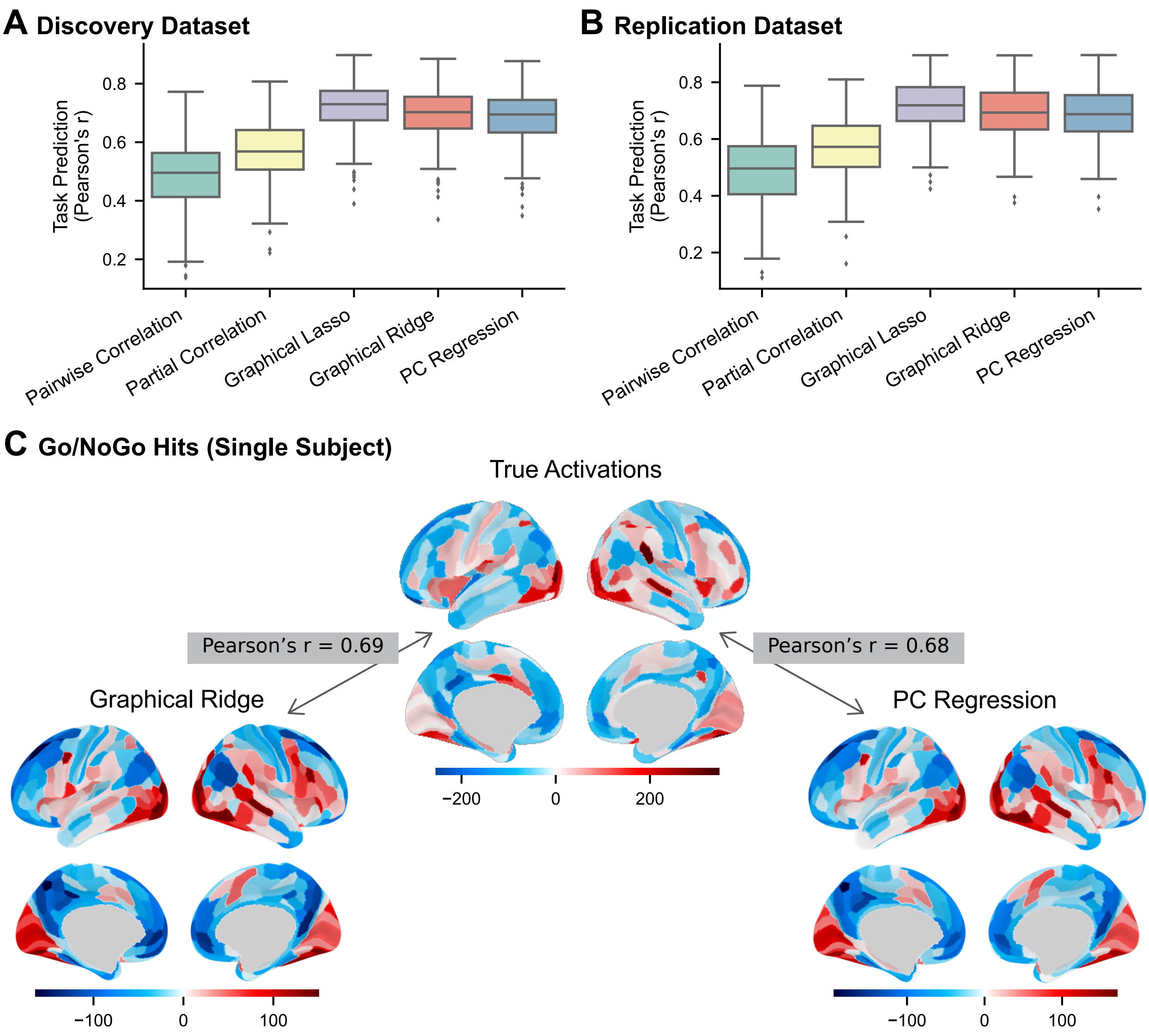


**Figure S8 – Predicting task activations using FC estimated from rest fMRI data – discovery and replication datasets, all FC methods. A-B)** Prediction accuracy by FC method calculated as the Pearson correlation (r) between regions' actual and predicted activations, computed in each subject (n = 236) across regions and task conditions. Results are from the discovery dataset **(A)** and replication dataset **(B)**. **C)** A single subject's actual and predicted (from graphical ridge and PC regression FC) task activations for the go/no-go hit events.

**Table S5 – Statistical tests comparing FC methods on task activation, subject age, and intelligence prediction accuracy – discovery and replication datasets.** For task activation prediction accuracies, scores for all subjects (Pearson's r) were compared using two-tailed, dependent-sample t-tests. For age and intelligence, predicted-to-actual correlations (single Pearson's r per method) were compared between methods using the two-tailed test described by Meng, Rosenthal, and Rubin (1992), which tests for a significant difference between "correlated correlations" (correlations with a shared variable – actual subject age and psychometric *g* in this case). The Pearson's r values were normalized using Fisher's z transformation (arctanh) before statistical comparisons. For reported correlation differences, correlations were transformed to Fisher's z, subtracted, and then transformed back to Pearson's r. Alpha levels were adjusted to .005 (.05/10) following Bonferroni correction to account for the multiple comparisons within each column.

| Methods | | Task Activity Prediction (Pearson's r) | | Age Prediction (Pearson's r) | | Intelligence Prediction (Pearson's r) | |
|---|---|---|---|---|---|---|---|
| | | Discovery | Replication | Discovery | Replication | Discovery | Replication |
| | Pairwise Correlation | mean r = 0.488, SD = 0.119 | mean r = 0.489, SD = 0.124 | r = 0.739, p < .00001 | r = 0.760, p < .00001 | r = 0.309, p < .00001 | r = 0.363, p < .00001 |
| | Partial Correlation | mean r = 0.569, SD = 0.106 | mean r = 0.568, SD = 0.107 | r = 0.512, p < .00001 | r = 0.579, p < .00001 | r = 0.114, p = .079 | r = 0.113, p = .083 |
| | Graphical Lasso | mean r = 0.719, SD = 0.086 | mean r = 0.717, SD = 0.085 | r = 0.695, p < .00001 | r = 0.737, p < .00001 | r = 0.373, p < .00001 | r = 0.402, p < .00001 |
| | Graphical Ridge | mean r = 0.694, SD = 0.090 | mean r = 0.694, SD = 0.089 | r = 0.702, p < .00001 | r = 0.745, p < .00001 | r = 0.404, p < .00001 | r = 0.404, p < .00001 |
| | PC Regression | mean r = 0.684, SD = 0.093 | mean r = 0.685, SD = 0.091 | r = 0.709, p < .00001 | r = 0.754, p < .00001 | r = 0.421, p < .00001 | r = 0.415, p < .00001 |
| 1 | Pairwise Correlation vs. Partial Correlation | mean Δr = -0.081, T(235) = -16.4, p < .00001 | mean Δr = -0.080, T(235) = -15.3, p < .00001 | Δr = 0.366, Z = 5.34, p < .00001 | Δr = 0.324, Z = 5.00, p < .00001 | Δr = 0.201, Z = 2.48, p = .013 | Δr = 0.260, Z = 3.16, p = .002 |
| 2 | Pairwise Correlation vs. Graphical Lasso | mean Δr = -0.231, T(235) = -48.9, p < .00001 | mean Δr = -0.229, T(235) = -47.0, p < .00001 | Δr = 0.091, Z = 1.59, p = .112 | Δr = 0.053, Z = 0.922, p = .356 | Δr = -0.072, Z = -1.05, p = .292 | Δr = -0.046, Z = -0.639, p = .523 |
| 3 | Pairwise Correlation vs. Graphical Ridge | mean Δr = -0.206, T(235) = -46.1, p < .00001 | mean Δr = -0.206, T(235) = -44.3, p < .00001 | Δr = 0.078, Z = 1.36, p = .175 | Δr = 0.036, Z = 0.647, p = .518 | Δr = -0.109, Z = -1.80, p = .072 | Δr = -0.049, Z = -0.726, p = .468 |
| 4 | Pairwise Correlation vs. PC Regression | mean Δr = -0.196, T(235) = -43.9, p < .00001 | mean Δr = -0.197, T(235) = -42.6, p < .00001 | Δr = 0.063, Z = 1.13, p = .259 | Δr = 0.014, Z = 0.259, p = .796 | Δr = -0.129, Z = -2.10, p = .036 | Δr = -0.062, Z = -0.937, p = .349 |
| 5 | Partial Correlation vs. Graphical Lasso | mean Δr = -0.149, T(235) = -60.4, p < .00001 | mean Δr = -0.149, T(235) = -58.6, p < .00001 | Δr = -0.285, Z = -5.45, p < .00001 | Δr = -0.276, Z = -5.94, p < .00001 | Δr = -0.270, Z = -4.08, p = .00004 | Δr = -0.302, Z = -4.43, p < .00001 |
| 6 | Partial Correlation vs. Graphical Ridge | mean Δr = -0.125, T(235) = -54.5, p < .00001 | mean Δr = -0.126, T(235) = -52.8, p < .00001 | Δr = -0.297, Z = -5.99, p < .00001 | Δr = -0.292, Z = -6.53, p < .00001 | Δr = -0.304, Z = -4.73, p < .00001 | Δr = -0.305, Z = -4.58, p < .00001 |
| 7 | Partial Correlation vs. PC Regression | mean Δr = -0.115, T(235) = -46.3, p < .00001 | mean Δr = -0.117, T(235) = -44.2, p < .00001 | Δr = -0.311, Z = -6.11, p < .00001 | Δr = -0.311, Z = -6.54, p < .00001 | Δr = -0.322, Z = -4.67, p < .00001 | Δr = -0.317, Z = -4.52, p < .00001 |
| 8 | Graphical Lasso vs. Graphical Ridge | mean Δr = 0.024, T(235) = 23.1, p < .00001 | mean Δr = 0.023, T(235) = 21.8, p < .00001 | Δr = -0.014, Z = -0.492, p = .623 | Δr = -0.017, Z = -0.653, p = .514 | Δr = -0.037, Z = -1.01, p = .315 | Δr = -0.003, Z = -0.071, p = .943 |
| 9 | Graphical Lasso vs. PC Regression | mean Δr = 0.034, T(235) = 25.0, p < .00001 | mean Δr = 0.032, T(235) = 23.6, p < .00001 | Δr = -0.029, Z = -0.999, p = .318 | Δr = -0.039, Z = -1.52, p = .129 | Δr = -0.057, Z = -1.41, p = .159 | Δr = -0.016, Z = -0.365, p = .715 |
| 10 | Graphical Ridge vs. PC Regression | mean Δr = 0.010, T(235) = 16.2, p < .00001 | mean Δr = 0.009, T(235) = 13.8, p < .00001 | Δr = -0.015, Z = -0.844, p = .399 | Δr = -0.021, Z = -1.37, p = .171 | Δr = -0.020, Z = -0.943, p = .346 | Δr = -0.013, Z = -0.559, p = .576 |

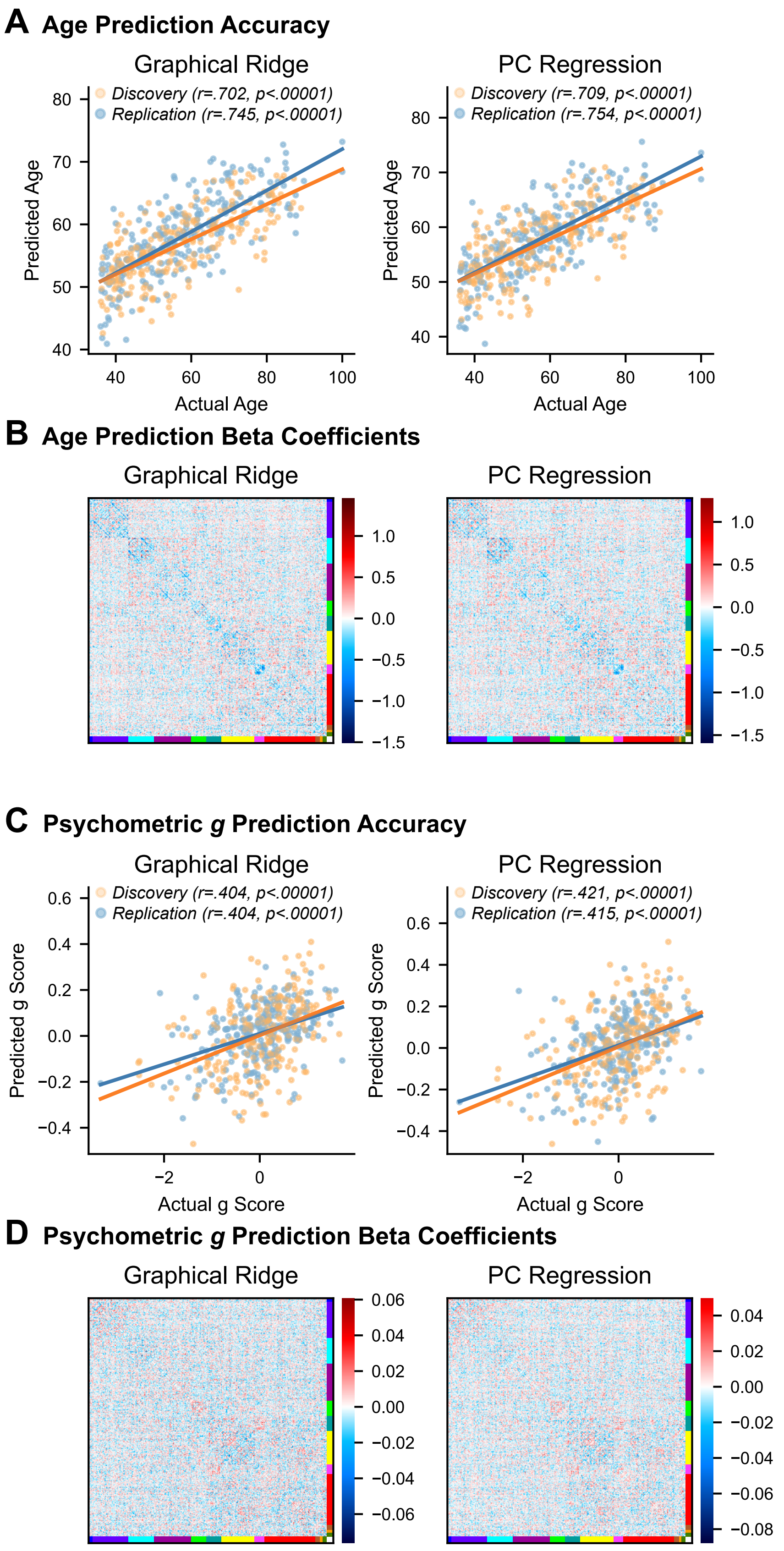
A Age Prediction Accuracy
Graphical Ridge
PC Regression
Discovery (r=.702, p<.00001)
Replication (r=.745, p<.00001)
Discovery (r=.709, p<.00001)
Replication (r=.754, p<.00001)
Predicted Age
Actual Age
B Age Prediction Beta Coefficients
Graphical Ridge
PC Regression
C Psychometric g Prediction Accuracy
Graphical Ridge
PC Regression
Discovery (r=.404, p<.00001)
Replication (r=.404, p<.00001)
Discovery (r=.421, p<.00001)
Replication (r=.415, p<.00001)
Predicted g Score
Actual g Score
D Psychometric g Prediction Beta Coefficients
Graphical Ridge
PC Regression

**Figure S9 – Predicting individual differences in age and intelligence using estimated FC from rest fMRI data – graphical ridge and PC regression. A)** Actual and predicted ages of each subject by FC method, from both the discovery and replication datasets. **B)** Average beta coefficients assigned to each connection by the regression models for estimating subject age. Beta coefficients shown here are the averages over cross-validation folds and discovery and replication datasets. Blue indicates that connection strength decreased with age and red indicates that connection strength increased. **C)** Actual and predicted intelligence (psychometric *g*) of each subject by FC method. **D)** Average beta coefficients assigned to each connection by the regression models for estimating psychometric *g*. See Figure 9 for results using pairwise correlation, partial correlation, and graphical lasso for FC estimation.
